# New genome reveals molecular signatures of adaptation to nocturnality in moth-like butterflies (Hedylidae)

**DOI:** 10.1101/2023.11.21.568084

**Authors:** Rachit Pratap Singh, Yi-Ming Weng, Yash Sondhi, David Plotkin, Paul B. Frandsen, Akito Y. Kawahara

## Abstract

Nearly all animals have a preferred period of daily activity (diel-niche), which is strongly influenced by the light environment. Sensory systems, particularly vision, are adapted to light, and evolutionary transitions to novel light environments, especially light limited ones, can impose strong constraints on eye evolution, color, and motion vision. The adaptive changes in sensory abilities of animals during these transitions, both at the genetic and neural levels, are largely unexplored. Butterflies and moths, with their diverse diel-niche shifts, are an ideal group for investigating the gene evolution linked to these transitions. While most butterflies are day-flying, hedylid butterflies are unique in being primarily nocturnal, and they represent an important evolutionary shift from diurnality to nocturnality in this clade. Here, we sequence the first high-quality Hedylidae genome and functionally annotate genes to understand genomic changes associated with shifts in diel niche. Comparing Hedylidae visual genes against day- and night-flying Lepidoptera species revealed that visual genes are highly conserved, with no major losses. However, hedylid butterfly opsins were more similar to nocturnal moths than their diurnal congeners. Tests on the evolutionary rates (dN/dS) confirmed that color vision opsins were under strong selection, similar to nocturnal moths. We propose that a convergent event of sequence evolution took place when these butterflies became nocturnal, approximately 98 million years ago.

## Introduction

The natural world is subject to a continuous day-night cycle, with drastic changes in both intensity and spectral composition of the light environment^1,2^. For animals, this cycle represents a critical temporal framework within which they conduct their activities. A clear dichotomy often exists between diurnal (day-active) and nocturnal (night-active) animals, with many insects^3,4^, birds^5^ and mammals^6^ restricted to either of the activity periods, often called diel-niche. While binning into nocturnal and diurnal categories is a simplification, given that animals occupy varied activity periods, this allows the examination of diel patterns and how lifestyles shift over an evolutionary time scale^7,8^.

Shifting from bright to dim environments presents unique challenges for an animal’s sensory systems, especially vision. Mammals, for instance, have evolved a wide range of eye shapes, such as large corneas and pupils that maximize light intake during nocturnal forays^9,10^. In birds, eye size is linked to light habitat and foraging behaviour^11^. Smaller animals such as insects are often limited by absolute eye size. Insects have developed intricate visual systems, featuring compound eyes with distinct arrangements of ommatidia. While mammals can use muscles to contract their pupils, insect eyes have a system of migrating pigment and graded refractive indices that can bend and manipulate light, pooling it as required to increase sensitivity or resolution. These eyes are often categorized into two broad classes: superposition and apposition eyes, each tailored to suit the animal’s preferred light environment^12,13^.

Adapting to dimly lit environments also entails subtle adjustments of the visual and nervous system. For example, many animals, including humans, trade off color vision for more sensitive monochromatic vision at night^14,15^. Similarly, nervous systems slow down and often sacrifice spatial and temporal resolution to increase sensitivity^16^. Studying visual systems usually necessitates time-intensive techniques such as behavioral observations or electrophysiology. However, the advent of more accessible genome and transcriptome sequencing methods have allowed for increased exploration of these patterns. For example, through the examination of genes responsible for vision, it was found that nocturnal owls have a reduced set of color vision genes compared to diurnal birds^17^. The same variation has been reported in fishes, where species adapted to bright environments have a much greater set of color vision genes than species that live in the deep sea^18^. In insects, multiple cases of gene duplications and losses have been observed, owing to the strong selective pressure imposed by light availability^19,20^. Butterflies and moths (Lepidoptera) are a prime example, where duplications and color vision gene diversification is much more prevalent in diurnal species as compared to nocturnal species^21–23^. This diversification of color vision genes aligns with the over 100 diel transitions recorded in Lepidoptera, featuring multiple evolutionary switches between nocturnality and diurnality^24^.

Butterflies have emerged as a valuable model system to explore the evolution of color vision in insects. Hedylidae, commonly known as American moth-butterflies, comprise a single genus, *Macrosoma*, with 36 described species^25^ and stand out among their butterfly counterparts in that nearly all species are nocturnal. Marked by moth-like attributes such as filiform antennae and nocturnal flight, hedylids were long classified within the moth superfamily Geometroidea^26^. Hedylids also possess tympanal ears on their wings, which they use to detect echolocation and defend themselves from nocturnal bat predation^27,28^. Recent phylogenetic analyses have revealed that Hedylidae likely diverged from their sister group, the skippers (Hesperiidae), less than 30 Mya^29,30^. Studies exploring the sensory ecology of hedylids suggest they possess refracting superposition eyes, which evolved exclusively in Hedylidae and Hesperiidae among butterflies^31,32^. Because they are nested in a diurnal clade, hedylids are an ideal model to study how shifts to a new light environment influence gene evolution. We present the first annotated genome assembly of the hedylid *Macrosoma leucophasiata,* generated with PacBio Hi-Fi sequencing. Leveraging this new resource, we examined the visual gene repertoire and conducted a comparative analysis of curated phototransduction genes using genome assemblies of candidate Lepidoptera species. Since the repertoire size of phototransduction genes is thought to be generally consistent across species with different diel niches, we expected to see the gene repertoire in *M. leucophasiata* to follow the same tendency. We focused our hypotheses on opsin genes, given their strong genotype-phenotype links and key role in optimizing vision. Firstly, we tested if opsin genes in species sharing the same diel niche show sequence similarity and whether hedylid opsins are more closely related to butterflies or moths. To infer the selection pressure on these genes, we also tested for positive selection. These analyses, with a focus on Hedylidae, provide new insights into the molecular adaptations associated with the switch to a nocturnal niche in butterflies.

## Results

### Contig-scale functionally annotated reference genome assembly for Hedylidae

We sequenced the genome of *Macrosoma leucophasiata* (**Supplementary Figure 1**) using PacBio high-fidelity (HiFi) sequencing. A total of 2.85 million HiFi reads were obtained, resulting in 32x read coverage. A preliminary survey using Genomescope 2.0 estimated the genome size to be 452 Mbp by k-mer analysis, with a heterozygosity rate of 1.61% (**Supplementary Figure 2**). The curated Hifiasm assembly resulted in a genome size of 616 Mbp comprising 66 contigs and N50 value of 22.3 Mbp (**Supplementary Data 1**). The draft assembly was generated after filtering haplotigs via purging and non-target sequence removal using BlobTools^33^. We identified and removed 152 contigs containing 2.82 Mbp (0.11%) of contamination linked to Euglenozoa from the draft assembly. Similarly, haplotig purging led to further removal of duplicated and mismatched contigs (**Supplementary Data 2**). We assessed BUSCO completeness at each step to ensure consistency and prevent the loss of meaningful information. Assembly statistics and BUSCO scores are shown in **Table 1**.

**Table 1:**
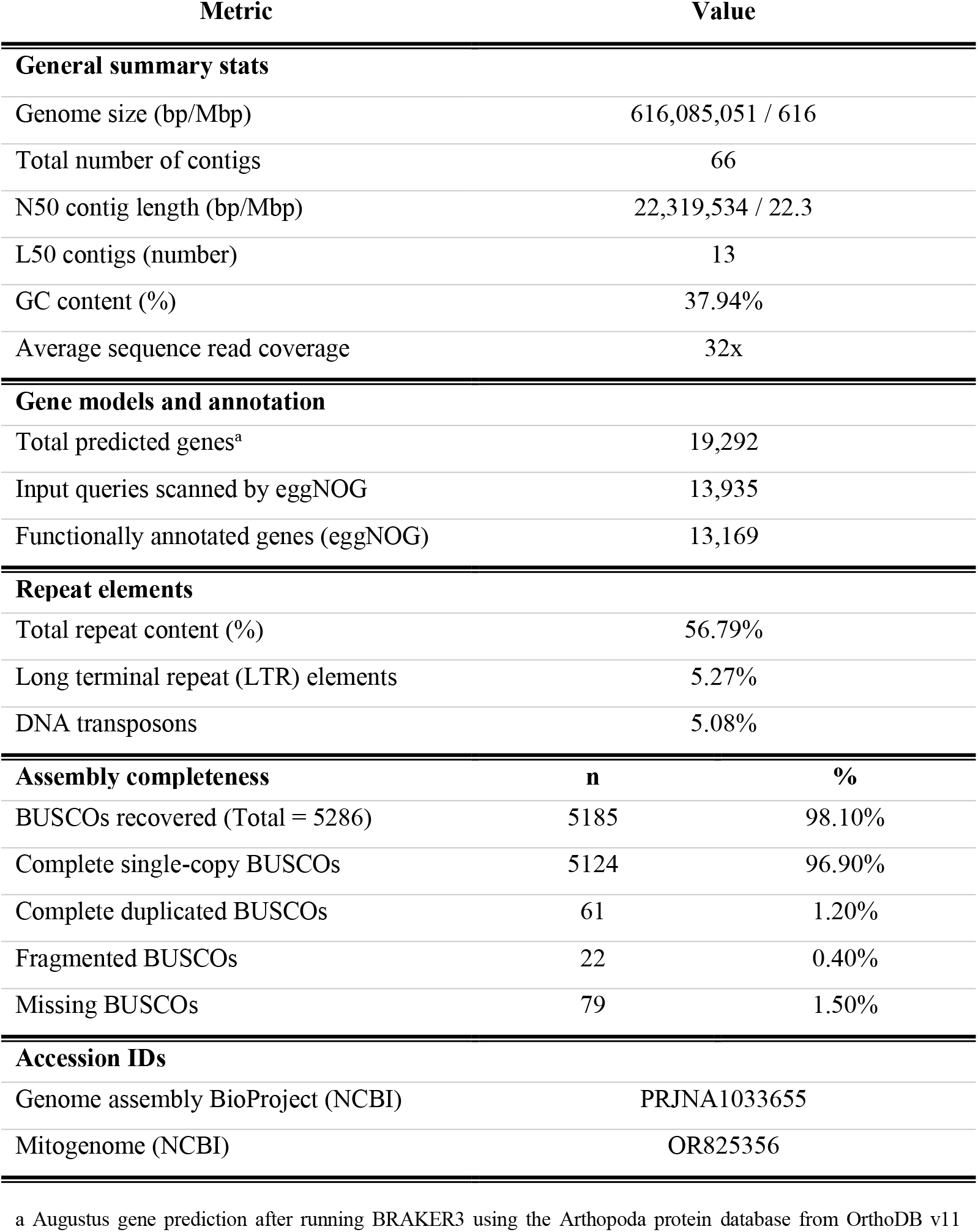
Summary and key characteristics of *Macrosoma leucophasiata* (Hedylidae) genome assembly and annotation.

| Metric | Value |  |
| --- | --- | --- |
| General summary stats |  |  |
| Genome size (bp/Mbp) | 616,085,051 / 616 |  |
| Total number of contigs | 66 |  |
| N50 contig length (bp/Mbp) | 22,319,534 / 22.3 |  |
| L50 contigs (number) | 13 |  |
| GC content (%) | 37.94% |  |
| Average sequence read coverage | 32x |  |
| Gene models and annotation |  |  |
| Total predicted genes<sup>a</sup> | 19,292 |  |
| Input queries scanned by eggNOG | 13,935 |  |
| Functionally annotated genes (eggNOG) | 13,169 |  |
| Repeat elements |  |  |
| Total repeat content (%) | 56.79% |  |
| Long terminal repeat (LTR) elements | 5.27% |  |
| DNA transposons | 5.08% |  |
| Assembly completeness |  |  |
|  | n | % |
| BUSCOs recovered (Total = 5286) | 5185 | 98.10% |
| Complete single-copy BUSCOs | 5124 | 96.90% |
| Complete duplicated BUSCOs | 61 | 1.20% |
| Fragmented BUSCOs | 22 | 0.40% |
| Missing BUSCOs | 79 | 1.50% |
| Accession IDs |  |  |
| Genome assembly BioProject (NCBI) | PRJNA1033655 |  |
| Mitogenome (NCBI) | OR825356 |  |

<sup>a</sup> Augustus gene prediction after running BRAKER3 using the Arthropoda protein database from OrthoDB v11

The circular mitochondrial genome of *M. leucophasiata* was also assembled into a single contig with a length of 15,209 bp. The mitogenome was annotated with 36 protein-coding genes, 22 t-RNA genes and two ribosomal RNA genes. The mitogenome is similar in size to the mitogenome of *M. conifera* (15,344 bp)^34^.

To assess the extent of repetitive sequences in the final assembly of *M. leucophasiata*, we modeled and soft-masked repeat regions using RepeatModeler2^35^ and RepeatMasker^36^. A library of 1082 transposable elements (TE) repeat families was generated corresponding to the Dfam database^37^. Repeat sequences accounted for 56.79% (349 Mbp) of the assembly. For gene annotation, we used the resulting soft-masked genome to run the BRAKER3 pipeline with protein evidence from the OrthoDB v11^38^ catalog for Arthropoda to obtain the gene model. A total of 19,292 protein-coding genes were predicted in the resulting gene model, and were used for functional annotation (**Supplementary Data 3**). We annotated gene function primarily using eggNOG-mapper^39^ and by performing sequence-similarity searches against the Swiss-Prot arthropodan database using DIAMOND v2.0.9^40^. In eggNOG, a total of 13,169 (68.26% of the predicted genes) were functionally annotated (**Table 1**). Gene ontology (GO) terms for 6,598 (∼50%) of the annotated genes were obtained (**Supplementary Data 4**).

### Gene family evolution analysis suggests conservation in vision-related genes with variable gene tree topologies in Opsins

Our ML tree based on 3,376 BUSCO single-copy orthologs was well-supported (100% support for SH-aLRT and ultrafast bootstrap, **Figure 1**). Relationships of butterfly families are consistent with most previous studies where Hedylidae and Hesperiidae are sister groups, and Papilionidae (swallowtails) are the first branching family within the Papilionoidea superfamily^29,30^. We identified 26,990 hierarchical orthogroups (HOGs) with OrthoFinder (**Supplementary Data 5**) from the primary transcripts of Augustus gene models of the 20 species generated from the BRAKER3 pipeline (**Supplementary Table 1**). We evaluated gene repertoire size evolution on the branch leading to our focal species, *M. leucophasiata*, using CAFE^41^ and found 64 HOGs at this branch are under rapid gene expansion and eight are under rapid gene contraction. These HOGs include many retrotransposon-related genes, odorant receptors, and cytochrome P450 genes. However, none of the candidate vision-related HOGs had a significant repertoire size change (**Supplementary Data 6**).

**Figure 1.**
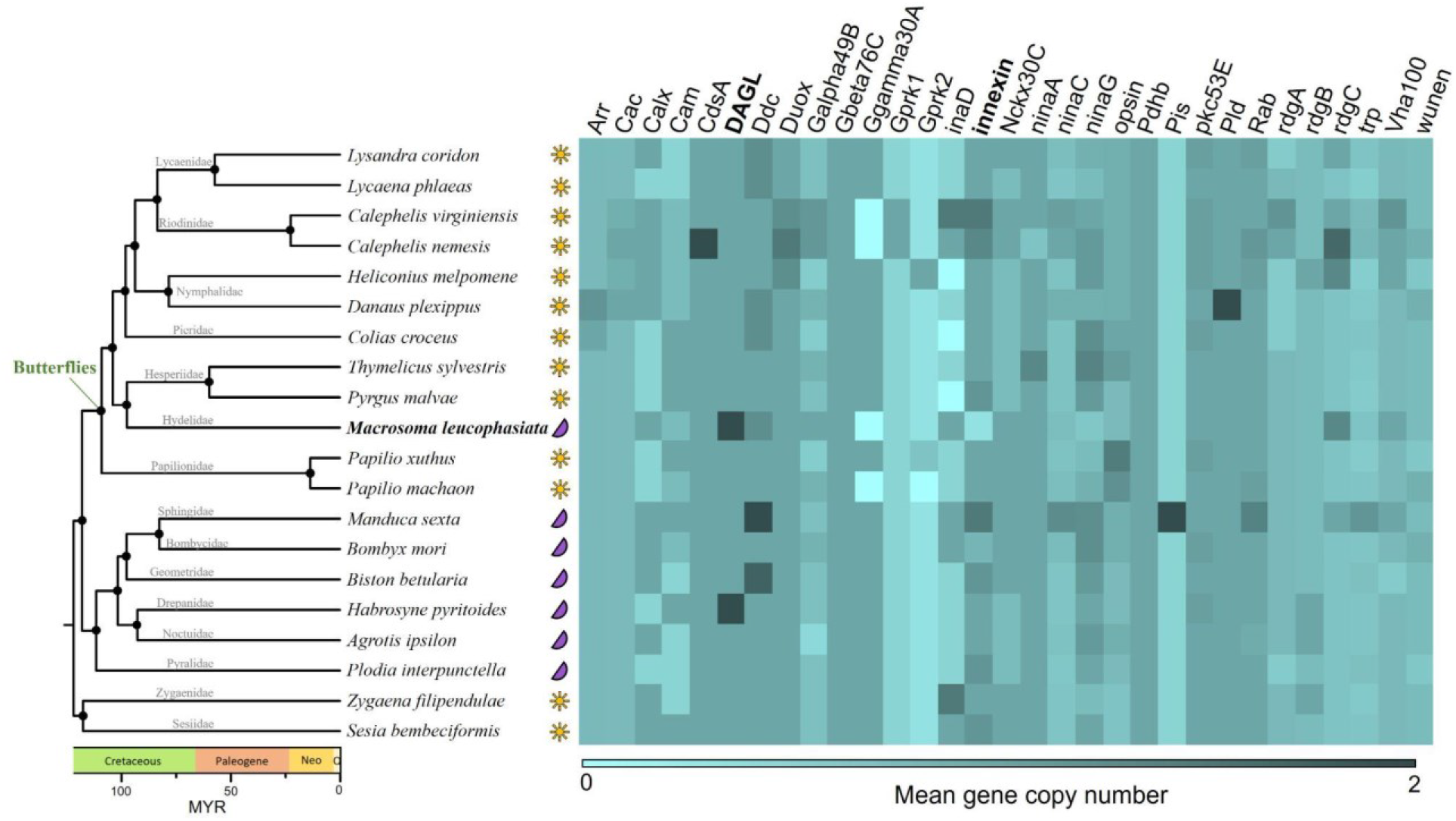
Analysis of visual genes shows high conservation with day and night flying butterflies with gene losses in the *DAGL* and *innexin* in *M. leucophasiata*. Maximum likelihood tree of the 20 selected lepidopteran species (Representing six butterfly and eight moth families) based on 3,376 BUSCO single-copy orthologs (left), and the gene count matrix of phototransduction-related gene families (right). The diel niche of each species is indicated by orange suns (diurnal) and purple moons (nocturnal). The mean gene count (repertoire size) is shown in the colored heatmap.

We searched the vision gene sequences against the database of 32 phototransduction-related gene families (from Macias-Muñoz et al.^42^) using BLASTp, and identified 142, 149, and 180 putative phototransduction-related genes, clustered in 179 HOGs, for *Danaus plexippus*, *Heliconius melpomene*, and *Manduca sexta*, respectively. A total of 3,503 genes from the 20 species in these 179 HOGs were considered putative phototransduction-related gene orthologs. In addition, keyword searches in the EggNOG annotations of the remaining 17 species resulted in the extraction of 23 additional HOGs. The unfiltered vision model from the 20 species thus consisted of 3,721 genes in 202 HOGs (**Supplementary Data 7**). A total of 345 genes were removed and three HOGs lost all their genes after filtering out the putative vision genes without corresponding function (defined from the EggNOG annotation; **Supplementary Data 8**). Among these genes in the filtered gene set, 90% (2995 genes) are single copy and 10% are duplicated. Among duplicated orthologs, 45 (1.35%) genes from 26 HOGs met the first criterion where the sum of sequence lengths are in the range of mean ± standard deviation of other single copy sequences, 40 (1.2%) from 22 HOGs passed the second criterion, and 34 (1%) gene from 21 HOGs passed all criteria and were assembled into a single sequence. The false duplication correction resulted in a vision gene count matrix consisting of 3,316 genes in 199 HOGs for these 20 species (**Supplementary Data 9**). According to this gene count matrix, repertoire sizes of the vision genes are generally consistently single-copy across the 20 species, with greater copy number variations found in the *trp* and *ninaC* families (**Supplementary Data 9**). Noticeably, four out of six orthologs of the innexin gene family are missing in *M. leucophasiata* (**Figure 1**).

The 32 phototransduction-related gene family trees differed, but genes from butterfly species generally tended to cluster together (**Supplementary Data 10**). Notably, three out of the seven opsin genes (UVRh, BRh, and LWRh) in *M. leucophasiata* were grouped with nocturnal moths rather than butterflies with fairly strong branch supports (>80 supports from SH-aLRT and ultrafast bootstrap) (**Figure 2 & Supplementary Data 11**). Surprisingly, this pattern is only found in a few other genes (**Supplementary Data 10**).

**Figure 2.**
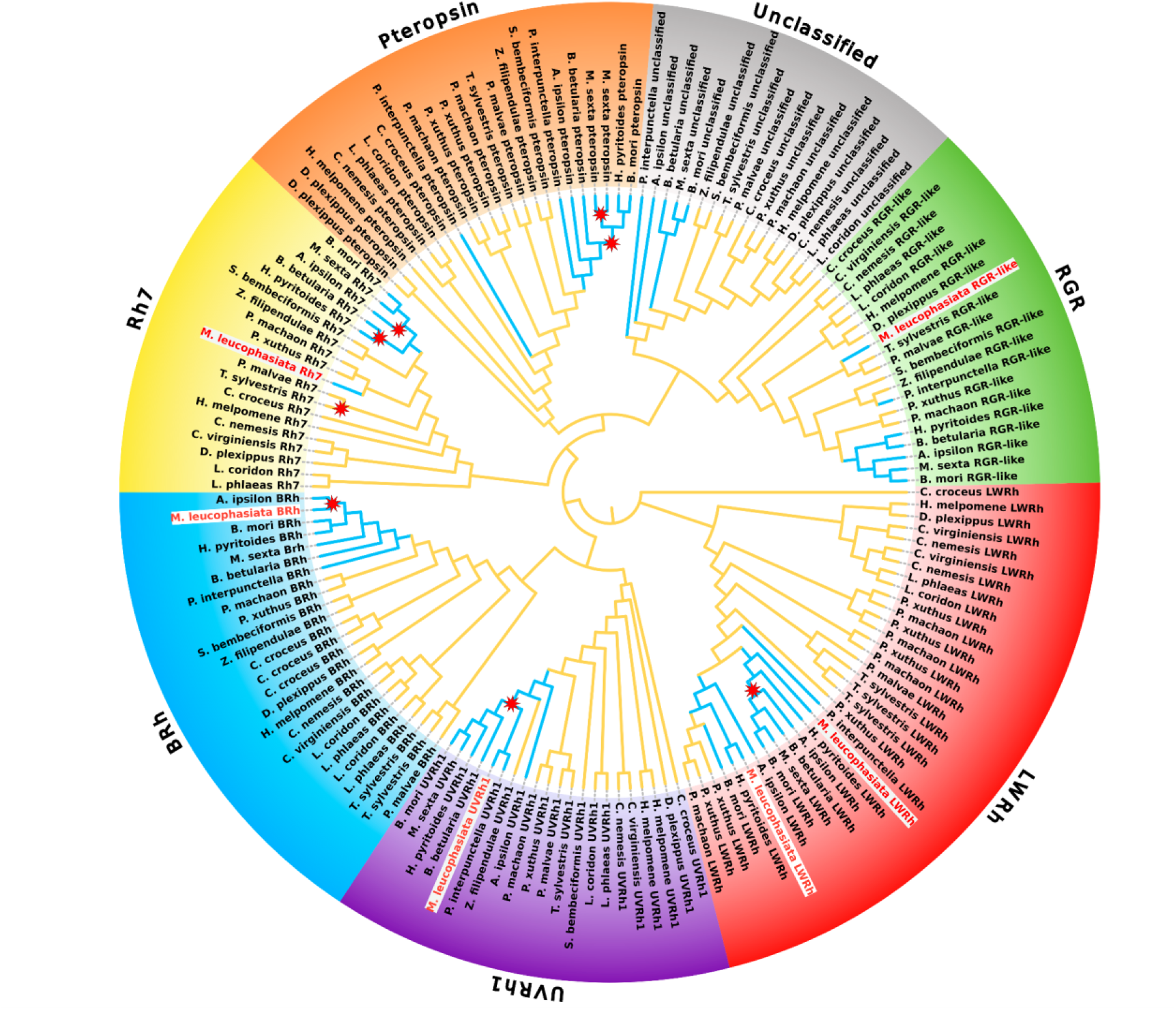
Convergence of visual opsins in *Macrosoma leucophasiata* with nocturnal moth species highlighting branches under positive selection. Opsin gene family tree showing the seven visual opsins in the studied species. Colored branches represent nocturnal (blue) and diurnal (orange) species. Stars indicate branches or nodes detected to be under positive selection by the aBSREL test for episodic diversification.

### Selection of opsin genes

Our selection analyses on opsins uncovered evidence of positive selection acting on multiple branches and nodes within the gene trees. Specifically, we tested the hypothesis that opsin genes in nocturnal and diurnal taxa vary in their rates of selection (dN/dS ratio) owing to adaptive (positive) selection. We used branch-site (aBSREL^43^) and site-substitution (MEME^44^) models to detect branches and identify the individual sites under selection. *BRh* showed significant evidence of positive selection on the branch leading to *A. ipsilon* (LRT 11.01, p= 0.018). In *UVRh1*, the clade representing all nocturnal taxa, including *M. leucophasiata*, was detected to have undergone diversifying selection (LRT 9.89, p= 0.037). The inferred *ω* rate classes among the branches were evenly split (*ω_1_* = 49%, *ω_2_*= 51%) in the *UVRh1* phylogeny with no *ω_3_* classes identified. Similarly, *LWRh* opsins indicated signal for positive selection with inferred fluctuating selection pressure on the nodes, representing four nocturnal taxa with LW duplications (LRT 13.65, p= 0.007) (**Figure 2**). In *Pteropsin*, we identified sites on two branches of the nocturnal clade classified as being under positive selection (LRT 9.41, 40.96; p= 0.047, <0.001) including a disproportionately higher distribution of *ω_2_* rate class (62% of branches, 92% of tree length). We recovered the *M. sexta* duplication of *pteropsin* as shown in Macias-Muñoz et al.^42^, and detected a high proportion of sites with dN/dS > 1 on that branch. No significant signal of selection was observed for the *RGR-like* and Unclassified (*UnRh*) opsins. Branches under positive selection are annotated on the opsin gene family tree in **Figure 2** and detailed selection statistics are provided in **Supplementary Table 2**.

Furthermore, employing the site-substitution model, MEME, within the HyPhy suite, we conducted a comprehensive examination of branches identified as being under positive selection by aBSREL. We examined specific sites within these genes and compared them, following the methods of Murrell et al.^44^. Based on the likelihood ratio test, we found that episodic diversifying selection has acted on multiple sites in the opsin sequences (**Supplementary Data 12**). The sites detected to be under diversifying selection were examined using the interactive plots on ObservableHQ. We used the Empirical Bayes’ Factor evidence ratio as an exploratory tool to assess the support for positive selection at the reported sites. Notably, multiple sites in the *BRh*, *UVRh1, Pteropsin* and *LWRh* sequences were observed with strong support for positive selection in *M. leucophasiata* (**Supplementary Figure 3**).

## Discussion

In this study, we generated the first, high-quality genome of Hedylidae to understand the genetic components underlying transitions to nocturnality. While visual opsin genes were a major focus, we also examined a range of phototransduction genes that displayed a conserved pattern of gene copy numbers across all species. To achieve this, we employed a comparative genomics approach, analyzing multiple high-quality genomes of butterflies and moths to gain evolutionary insights into their visual systems. We further examined the discordant gene tree topologies and signatures of diversifying selection; potentially tied to the emergence of nocturnality in this unusual butterfly lineage.

### Novel genome assembly for Hedylidae

New genome assemblies are now being generated at a rapid pace, and they have created the opportunity to study molecular signatures of adaptation across diverse lineages. Genomic studies of non-model species have uncovered a wealth of new knowledge on diverse topics, such as polyphagy-linked gene family expansions^45^, long-distance moth migration^46^, genome size evolution^47^ and opsin evolution across Lepidoptera^22^. However, this influx of genomic data varies in taxonomic coverage and quality; often obscuring the genetic underpinnings of such non-model species^48,49^. Within Lepidoptera, Hedylidae and their sister group Hesperiidae (Skippers) demonstrate this disparity in genomic resources. In Hesperiidae, ∼12% (n= 420) of the total species have publicly available reference genomes^48,50^ that have been widely utilized in evolutionary studies^51,52^, while Hedylidae have none. Previous studies have highlighted the need to address the paucity of hedylid genomes^29,53,54^.

### Phototransduction gene evolution

Gains and losses of phototransduction genes are generally thought to be infrequent in insects despite a wide range of photic niches covered by different species^55,56^. This trend was also supported by our orthology analysis (**Figure 1**) with some exceptions in specific taxa and gene families including duplications of opsin genes in many butterfly species. Notable duplications are also found in individual species, such as the *DAGLβ* gene in *M. leucophasiata* and *H. pyritoides*, *Ddc* gene in *M. sexta* and *B. betularia*, *Pis* gene in *M. sexta* and *Pid* gene in *D. plexippus*.

Many opsin genes have undergone numerous gene duplication events in dragonflies and damselflies (Odonata)^57^. Parallel gene losses of phototransduction genes also have been reported in subterranean water beetles (Coleoptera: Dytiscidae)^58^. In general, opsin is the most well explored among the phototransduction gene families for its duplications and losses, where *LWRh* shows highest variability among the opsin genes^59^. Indeed, our opsin gene family tree shows more gene duplications in the *LWRh* clade than any others (**Figure 2**). The accumulation of *LWRh* in butterflies has been found to modify or expand spectral sensitivity^59,60^. Nevertheless, our analysis shows that no *LWRh* gene duplication is detected on the branch leading to *M. leucophasiata*, suggesting that the adaptation of night vision of this nocturnal butterfly was unlikely achieved through *LWRh* gene duplication.

Despite various approaches that we applied to detect phototransduction genes across genome assemblies, the absence of orthologous genes occurred in some species **(Supplementary Data 7)**. Considering that the quality of the genomes are generally high, the absence of orthologous genes is likely to be the result of gene loss or substantial divergence of the sequence. Interestingly, more than half (4/6) of *innexin* genes were absent in *M. leucophasiata* (only *inx3* and *inx9* were detected). Innexin proteins were found forming gap junction channels and playing important roles in nervous system development during embryogenesis in *Drosophila*^61^, as developmental defects are observed when depleting or down-regulating *innexin* genes in *Drosophila*^62,63^. In addition, *inx2* and *inx3* play significant roles in vision, including eye disk development and possibly phototransduction processes in different insect species^64–67^. *Macrosoma* possesses adaptive characteristics for nocturnal vision, including larger relative eye size and more corneal nipples on the facet surface of compound eyes^28^. Experimental approaches testing the relationship between adaptive eye morphology and the gain and loss of *innexin* genes may improve our understanding about the underlying genetic basis of nocturnal butterfly vision evolution. Finally, an unexpected *DAGLβ* gene duplication in *M. leucophasiata* was detected. *DAGLβ* was single-copy in most of the species studied and is thought to be involved in the phototransduction cascade and neural development^55^. The rare duplication of the *DAGLβ* gene in *M. leucophasiata* implies a potential role in the species’ adaptation to night vision.

### Adaptive molecular evolution and signatures of selection in opsin genes

Visual opsins play a key role in initiating the phototransduction cascade and are known to be considerably variable in their repertoire size across the animal kingdom. They represent major targets of adaptive evolution with direct, well-characterized connections established between opsin gene sequences and resulting wavelength sensitivities. While non-adaptive mechanisms can also affect the evolution of opsin sequences, adaptive forces such as changing light environments are more likely to cause consistent diversification^21^. Intense selective pressure imposed by changing light environments on insect visual systems has led to changes in the diversity of opsin gene repertoire to adapt to altered sensory demands. Insects, due to their varying lifestyles, often exhibit distinct patterns of opsin gene expression. Notable differences in opsin expression between diurnal and nocturnal Lepidoptera were reported by Macias-Muñoz et al.^42^, contrasting with Akiyama et al.^68^ who found no significant differences. This suggests that while light environments can influence opsin expression and loss, other factors also play a role in shaping these patterns.

Molecular adaptations of visual opsins are hypothesized to accompany diel niche transitions in Lepidoptera, conferring enhanced sensitivity under dim conditions^69^. Our finding that diversifying selection is targeting opsins in moth and moth-butterfly lineages, support this premise. Intriguingly, we observed a unique clustering of hedylid opsins with moths in our dataset, which runs counter to species relationships. While other factors could be involved, grouping with nocturnal taxa hints at possible convergent sequence evolution stemming from shared selective pressures. Previous studies have used selection analyses to further examine such discordance and characterize genes that evolve more rapidly than expected under neutral evolution^70–72^. We applied rigorous branch-site (aBSREL) and site-specific (MEME) models in HyPhy to identify specific lineages experiencing diversifying selection and reported signatures of episodic diversifying selection across multiple opsin sequences **(Supplementary Data 12)**. Positively selected sites and evidence of diversifying selection on the branches of blue (BRh) and UV (UVRh1) opsin sequences leading to *M. leucophasiata* hints at an adaptation to a crepuscular niche. Sensitivity to UV signals is important for night-active insects as it assists integral functions such as detecting foraging cues in night-blooming flowers^73^. Alternatively, the long-wavelength (LWRh) opsin expressing photoreceptors play an even more critical role in maintaining sensitivity in dim-light environments and are most numerous in both diurnal and nocturnal species. A clear clustering of LWRh opsin sequences was observed but no evidence for positive selection was observed on the hedylid branch **(Figure 2)**. It is possible that *M. leucophasiata* has retained the extended spectral sensitivity to longer wavelengths; generally prevalent and recurrently evolved in butterflies^74,75^. While multiple sites in the LWRh sequences were identified to be under diversifying selection, our analyses are limited by the lack of a proper characterization of the spectral tuning sites, which would enable us to understand whether these positively selected sites affect the visual range of LW opsins in Hedylidae. While selection analyses on opsins yielded insights, our study was limited by the lack of RNA sequencing data, which could have refined our gene models and helped validate exon-intron boundaries.

## Conclusions

Our genome assembly of *Macrosoma leucophasiata* provides the first reference genome for Hedylidae, which was of high quality **(Table 1)**. Addressing this knowledge gap collectively reinforces the significance of genomics in broader ecological and evolutionary contexts. Beyond vision genes, these assemblies could elucidate genetic underpinnings of other aspects of hedylid biology such as the evolution of tympanal ears, circadian rhythm and hostplant associations. The findings from selection analyses reported here can likely be examined further with transcriptomic profiling across diel categories in Hedylidae. Our findings showcase the power of leveraging genomic resources across lineages occupying diverse ecological niches. Future efforts to improve the assembly could also focus on generating chromosomal-scale data with synteny mapping.

## Materials and methods

### Sample preparation and sequencing

Two adult individuals of *Macrosoma leucophasiata* were collected by AYK in August, 2016 at the Wildsumaco Biological Station, Napo, Ecuador (0°40’17.2"S, 77°35’55.1"W, 1400m a.s.l). The butterfly wing vouchering and tissue storage methods followed Cho et al. (2016). Wing vouchers (LEP36876, LEP36882) and a tissue sample (MGCL_1059464) are stored at the McGuire Center for Lepidoptera and Biodiversity (MGCL), Florida Museum of Natural History (Gainesville, USA).

We extracted high molecular weight DNA from muscle tissue obtained from the abdomen and thorax using a Qiagen Genomic-tip DNA extraction kit. Following DNA extraction, DNA was visualized using pulse-field gel electrophoresis. DNA was sheared to ∼20 kbp using a Diagenode Megaruptor 3. We size-selected the DNA for fragments greater than 10 kbp using a Sage Science BluePippin for library preparation. We prepared a PacBio HiFi library with the SMRTbell Express Template Prep Kit 2.0 (PacBio, Menlo Park, CA, USA). The library was sequenced on a PacBio Sequel II instrument on an 8M SMRT cell in CCS mode with a 30-hour movie time. The library preparation and the sequencing of the DNA were conducted at the DNA Sequencing Center at Brigham Young University (Provo, Utah, USA).

### Genome profiling and coverage estimation

We performed quality control checks on the read quality of the raw high-fidelity (HiFi) sequence reads using FastQC v 0.11.7^76^. The k-mer density distribution was assessed from the HiFi reads using K-Mer Counter (KMC) v.3.2.1^77^ with the k-mer length being 21 nucleotides. We surveyed the genome characteristics based on the k-mer distribution analyses, using GenomeScope v2.0^78^ to predict the genome size, heterozygosity and assess genome quality. The GenomeScope 2.0 profile (**Supplementary Figure 2**) was generated using default settings for a diploid species. We used the resulting profile and other metrics to inform the choice of assembly parameters and allow stringency in error correction.

### Genome assembly and decontamination

To assemble the HiFi reads into contigs, we used Hifiasm v 0.16.1^79^ using default parameters with aggressive purging level 3 (-l 3) to filter erroneous and low-quality reads. We used QUAST v5.2.0 to assess the assembly statistics and generate a summary table for assessing contiguity and GC content^80^ (**Supplementary Data 1**). We followed the *Purge Haplotigs* pipeline, purge_haplotigs v1.1.2^81^ to sequentially identify and remove haplotigs and re-assign allelic contig pairings to generate a deduplicated assembly. The pipeline uses a haplotype-resolved assembly to generate mapped read coverages and aligns the contigs using *Minimap2* v. 2.21^82^ which are flagged as suspect and junk contigs, for haplotig removal.

We screened the purged assembly for contamination, in the form of non-target sequences, by examining GC-coverage plots using BlobTools v1.0^33^. To assess the read coverage, we aligned the HiFi reads to the assembly using Minimap2^82^ and sorted the aligned BAM file using *samtools sort*^83,84^. We then used BLASTn^85^ with an E-value of 1e-25 against the NCBI nucleotide database to facilitate the taxonomic assignment of these contigs. The resulting mapped files, coverage results, and BLAST output were used to generate BlobPlots^33^ (**Supplementary Figure 4**). We identified and removed the non-target sequences from the assembly after inspecting the sequence coverage, proportion, and variation in the GC content. Final assembly completeness was determined by repeating the BUSCO v5.3 calculation^86,87^, and the QUAST reports were compared to evaluate changes in assembly statistics after contamination removal (**Supplementary Data 2**).

### Mitogenome assembly

We assembled the mitochondrial genome for *Macrosoma leucophasiata* using MitoHiFi v3.2^88^. The raw PacBio HiFi reads and contigs from the unpurged Hifiasm assembly were used in two separate runs. We used the mitogenome of *Macrosoma conifera* with accession number NC_050856^34^ as a reference mitogenome. MitoFinder v1.4.0^89^ was used to annotate the final mitogenome (**Supplementary Figure 5**).

### Repeat region annotation

For the genome annotation pipeline, we first generated a *de novo* library of repeat sequences in the assembly using RepeatModeler2^35^. We soft-masked the repeat regions in the assembly with repetitive elements from various lines of evidence that were incorporated. This included the integration of simple and short repeats and the initially identified elements from RepeatModeler2. Additionally, repeats sourced from the lepidopteran entries in Repbase^90–92^ were utilized. We used the RMBLAST search within the RepeatMasker v4.1.1 software to complete the soft-masking steps^36^.

### Gene prediction and functional annotation

We used the fully automatic BRAKER3 pipeline for gene structure prediction of protein-coding genes in the soft-masked assembly^40,93–105^. We ran BRAKER using the arthropod protein sequences from OrthoDB v11 protein database^38^, following the GeneMark-EP+ pipeline for a final gene model predicted by AUGUSTUS and GeneMark-EP combined^97,98^. By relying solely on orthologous protein evidence, we aimed to overcome artifacts encountered during the transcript-mapping step. We assessed the completeness of the predicted gene models by comparing the BUSCO scores from the *lepidoptera_odb10* database for each BRAKER3 run. gFACs v1.1.2 was used to analyze the summary statistics and report the gene model profiles^106^ (**Supplementary Table 3**).

Functional annotation of the predicted genes from the transcript sequences was performed using the eggNOG-Mapper v2.1.9 web server^39^. We used stringent parameters for the annotation by restricting the taxonomic scope to ‘Arthropoda’ and including annotations with ‘experimental evidence only’. Additionally, a separate annotation file was also generated using the recommended default settings. Gene ontology (GO) terms were filtered and extracted from the eggNOG database annotations (**Supplementary Data 4**).

### Species tree and orthology inference

We constructed a butterfly-focused lepidopteran phylogeny with genomes from 20 species, including *M. leucophasiata*, 11 species from the other six butterfly families, and eight moth species including one each of Sesiidae and Zygaenidae, using BUSCO single-copy orthologs. Specifically, we downloaded genome assemblies of these species from GenBank, RefSeq, and Darwin Tree of Life to run BUSCO using the same settings as those that were applied for the *M. leucophasiata* genome (**Supplementary Figure 6**). The 3,376 BUSCO single-copy orthologs (amino acid sequences) were aligned using MAFFT v7.490 (the more accurate alignment option “mafft-linsi” was used) and concatenated into supermatrix alignment^107^. The maximum likelihood (ML) tree was constructed using IQ-tree v.2.0.3 with ’’Q.insect+FO+G4’’ substitution model^108,109^. Branch supports were assessed using 1,000 replicates SH-aLRT support and ultrafast bootstrap with the ’’bnni’’ option to avoid overestimating ultrafast bootstrap support^110,111^. The ML tree was converted to an ultrametric tree using treePL, and this tree served as the basis for subsequent gene family evolution analyses^112^. We re-annotated the 19 soft-masked publicly available genome assemblies using the BRAKER3 pipeline and the Augustus models generated were selected for downstream analyses. For these gene models, together with the model of *M. leucophasiata*, we used the longest transcript of each gene to perform orthology analysis using OrthoFinder v2.5.2, and the output phylogenetic hierarchical orthogroups (HOGs) were used to represent the orthologs^113^. The gene count matrix and the ultrametric tree were jointly used to detect the rapid repertoire size changes for each HOGs using CAFE v5.0.0^41,114^. We report our results on the rapidly evolving genes for the *M. leucophasiata* branch with a more stringent significance level of *p*-value= 0.01.

### Vision gene evolution

Since *M. leucophasiata* and many other hedylid species are well known for their distinctive nocturnal circadian rhythm compared to other butterfly species, we investigated the gene evolution of the putative vision-related genes. We used three lepidopteran species (*Danaus plexippus*, *Heliconius melpomene,* and *Manduca sexta*) that have been closely examined with their phototransduction-related genes in recent published work as benchmark species to identify the vision genes^55^. We downloaded phototransduction amino acid sequences for each of the three species and used BLASTp to identify these genes from the BRAKER gene models with a strict E-value threshold of 1E−50, as high sequence similarly was expected for intraspecific orthologous gene blast^115^. For the other 17 species, genes from the same HOGs with the blast-identified phototransduction genes were considered orthologs. In addition, we manually extracted the putative phototransduction-related genes by searching keywords from the eggNOG annotations of all species (**Supplementary Data 7**). Keywords were selected based on the eggNOG annotations of the BLASTp-determined phototransduction-related genes of the three benchmark species. To ensure the accuracy of the orthology of these annotations, we further blast the sequences against the NCBI non-redundant (nr) protein database using DIAMOND v2.1.8 with an E-value threshold of 1E−50^40,116^. Genes without phototransduction-related results in the top hits (max 50 hits) were removed from the vision gene model (**Supplementary Data 8**).

Considering that the recruited genome assemblies are not all chromosomal or near-chromosomal level, and not all the gene predictions have training transcriptomic data involved (**Supplementary Table 4**), the false gene duplication (an ortholog of a species that was divided into multiple small fragments due to the fragmented genome assemblies or incorrect gene prediction, and each was predicted as an independent gene and annotated with same function thus being identified as gene duplication) were further corrected. Specifically, we applied three criteria to determine the false gene duplication. First, the sum of duplicated gene sequence lengths is within the range of mean ± standard deviation of other single-copy orthologs. Second, the sequence similarities of the duplicated genes are less than mean pairwise sequence similarity of the ortholog. Third, genes were annotated in close tandem, in this case, we defined the distance as a span of five predicted genes. Sequence similarities (percent identities) were calculated using Clustal Omega v1.2.0 and the falsely duplicated genes were assembled using emboss v6.6.0 with other orthologous sequences as reference^117,118^. The falsely duplicated genes were removed from the gene count matrix which was jointly visualized using R package “phyloheatmap”^119^ (**Figure 1** and **Supplementary Data 9**). Finally, we clustered the genes of vision related HOGs and aligned sequences of each family using MAFFT v7.505 (mafft-linsi) to construct gene family trees using IQ-tree v.2.0.3^107,108^. We applied the best substitution model selected by ModelFinder and the remaining settings were the same as those applied to reconstruct the species phylogeny^120^.

### Selection analyses

Since some phototransduction-related genes, especially in the opsin gene family, showed discordant topology with the species tree (see results), we further tested for positive selection on the three opsin genes (i.e., *BRh*, *UVRh*, and *LWRh*). Testing for positive selection could provide additional clues in determining whether the discordant gene tree topologies are driven by selection or methodological limitations. We first built the individual gene trees using the same approach we used for gene family trees and viewed them on FigTree v1.43 (**Supplementary Data 11**) (http://tree.bio.ed.ac.uk/software/figtree/). The gene trees were then reconciled with the rooted species tree (**Figure 1**) using GeneRax^121^. We employed the undated DTL reconciliation model and SPR correction strategy in GeneRax to identify duplications, losses, and transfers. The reconciled gene trees were assessed using the ThirdKind reconciliation viewer^122^ (**Supplementary Data 13**). Codon alignments (DNA sequences) were generated using aligned protein sequences as references by PAL2NAL v14^123^. These codon alignments, along with the reconciled gene trees, were utilized for detecting positive selection. We employed the adaptive branch-site random effects model (aBSREL^43^) available in the HyPhy^124^ suite via the Datamonkey 2.0 webserver^125^. The foreground branches were defined by the branches of nocturnal species (**Figure 1**). The mixed-effects model of evolution (MEME^44^) was employed to detect episodic positive selection on individual sites, allowing for variations in selection pressures across the different branches and sites in the gene trees. Additionally, aBSREL was utilized to identify positive selection on specific branches, employing a random effects framework to model variable selection pressures. The branches detected to be under positive selection were identified using the likelihood ratio (LRT) test statistic and Bonferroni-Holm corrected p-values. We marked the branches and nodes with signatures of positive selection on to the gene family tree (**Figure 2** and **Supplementary Table 2**). We visualized and annotated the opsin gene family tree using the Interactive tree of life (iTOL) v. 6.5.7 tool^126^.

## Supporting information

Supplementary Data 1: Hifiasm assembly QUAST report

Supplementary Data 2: Final soft-masked assembly QUAST report

Supplementary Data 3: M. leucophasiata Augustus gene model

Supplementary Data 4: M. leucophasiata EggNOG annotation

Supplementary Data 5: Hierarchical orthogroups (HOGs) list

Supplementary Data 6: Rapidly evolving genes in M. leucophasiata

Supplementary Data 7: Unfiltered phototransduction-related orthologs

Supplementary Data 8: Filtered phototransduction-related orthologs

Supplementary Data 9: Corrected phototransduction-related orthologs

Supplementary Data 10: All (32) vision gene family trees

Supplementary Data 11: Opsin gene trees

Supplementary Data 12: MEME HyPhy Opsins summary

Supplementary Data 13: Reconciled gene trees ThirdKind

## Data availability

The sequence data including PacBio HiFi sequence reads and genome assembly are deposited in the NCBI database under bioproject PRJNA1033655 (sample ID: SAMN38040146). The raw HiFi reads are in the SRA database with accession number SRR26586523 and the assembly is available on genbank. The annotated mitochondrial genome is also available on NCBI with accession number OR825356.

## Author contributions

**RPS:** Conceptualization, Data curation, Formal analysis, Visualization, Writing- Original draft preparation, Reviewing and editing. **YMW:** Conceptualization, Data curation, Formal analysis, Visualization, Writing- Original draft preparation, Reviewing and editing. **YS:** Conceptualization, Validation, Writing- Reviewing and editing. **DP:** Resource acquisition, Writing- Reviewing and editing **PBF:** Resource acquisition, Project administration **AYK:** Conceptualization, Investigation, Validation, Writing- Reviewing and editing.

## Acknowledgements

We would like to thank Edward Wilcox at BYU for assisting molecular work, library preparation and sequencing, Taylor Pierson for helping with the wet lab resources, Joe Martinez for helping with accessing the museum specimens and photographing the specimens. Funding support for this project was through the National Science Foundation (NSF) grant DEB-1541500 to AYK. We also acknowledge University of Florida Research Computing (http://researchcomputing.ufl.edu) for providing computational resources and support that has contributed to the research results reported in this publication.

## Supplementary materials

Supplementary information: Supplementary tables and figures

Supplementary Data 1: Hifiasm assembly QUAST report

Supplementary Data 2: Final soft-masked assembly QUAST report

Supplementary Data 3: *M. leucophasiata* Augustus gene model

Supplementary Data 4: *M. leucophasiata* EggNOG annotation

Supplementary Data 5: Hierarchical orthogroups (HOGs) list

Supplementary Data 6: Rapidly evolving genes in *M. leucophasiata*

Supplementary Data 7: Unfiltered phototransduction-related orthologs

Supplementary Data 8: Filtered phototransduction-related orthologs

Supplementary Data 9: Corrected phototransduction-related orthologs

Supplementary Data 10: All (32) vision gene family trees

Supplementary Data 11: Opsin gene trees

Supplementary Data 12: MEME HyPhy Opsins summary

Supplementary Data 13: Reconciled gene trees ThirdKind

