## supplementary_information for "New genome reveals molecular signatures of adaptation to nocturnality in moth-like butterflies (Hedylidae)"

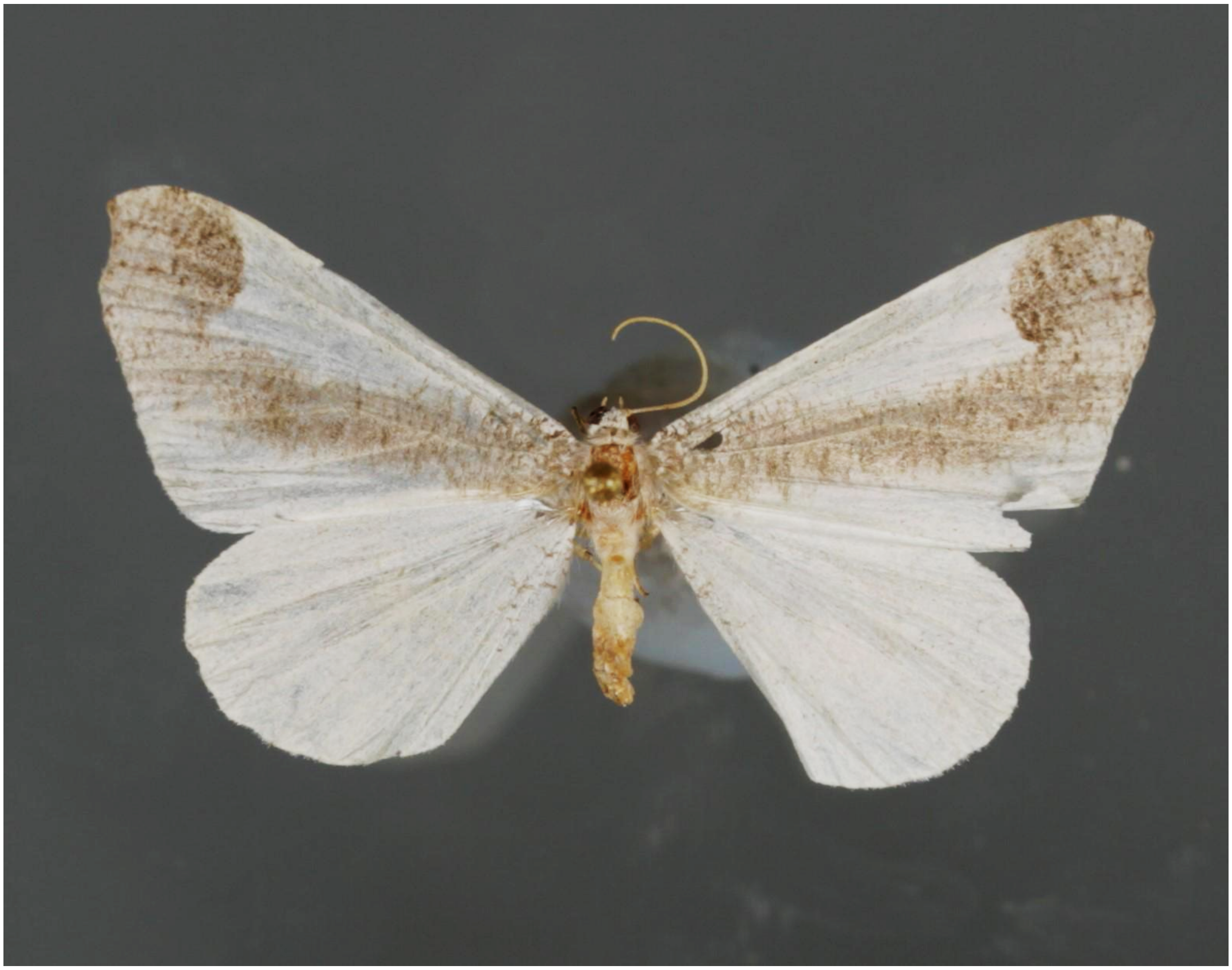

**Supplementary Figure 1.** Exemplar pinned specimen of *Macrosoma leucophasiata* deposited in the collections of McGuire Center for Lepidoptera and Biodiversity, Florida Museum of Natural History, Gainesville, FL USA.

**Supplementary Table 1.** Species involved in the gene family evolution analyses.

| Species | Family | Genome Source | Diel Niche | References |
|---|---|---|---|---|
| *Agrotis ipsilon* | Noctuidae | Darwin Tree of Life | Nocturnal | 1 |
| *Biston betularia* | Geometridae | Darwin Tree of Life | Nocturnal | 2 |
| *Bombyx mori* | Bombycidae | GenBank | Nocturnal | 3 |
| *Calephelis nemesis* | Riodinidae | GenBank | Diurnal | 4 |
| *Calephelis virginiensis* | Riodinidae | GenBank | Diurnal | 4 |
| *Colias croceus* | Pieridae | Darwin Tree of Life | Diurnal | 5 |
| *Danaus plexippus* | Nymphalidae | RefSeq | Diurnal | 5 |
| *Habrosyne pyritoides* | Drepanidae | Darwin Tree of Life | Nocturnal | 6 |
| *Heliconius melpomene* | Nymphalidae | GenBank | Diurnal | 5 |
| *Lycaena phlaeas* | Lycaenidae | Darwin Tree of Life | Diurnal | 7 |
| *Lysandra coridon* | Lycaenidae | Darwin Tree of Life | Diurnal | 7 |
| *Macrosoma leucophasiata* | Hedylidae | this study | Nocturnal | 8 |
| *Manduca sexta* | Sphingidae | RefSeq | Nocturnal | 3 |
| *Papilio machaon* | Papilionidae | Darwin Tree of Life | Diurnal | 9 |
| *Papilio xuthus* | Papilionidae | RefSeq | Diurnal | 9 |
| *Plodia interpunctella* | Pyralidae | RefSeq | Nocturnal | 10 |
| *Pyrgus malvae* | Hesperiidae | Darwin Tree of Life | Diurnal | 11 |
| *Sesia bembeciformis* | Sesiidae | Darwin Tree of Life | Diurnal | 9,12 |
| *Thymelicus sylvestris* | Hesperiidae | Darwin Tree of Life | Diurnal | 13 |
| *Zygaena filipendulae* | Zygaenidae | Darwin Tree of Life | Diurnal | 14 |

**Supplementary Table 2.** Statistics of branch-site model testing positive selection for opsin genes using aBSREL.

| Gene tree | Total branches chosen | Exploratory/ targeted | Branches detected | p-value (corr) | LRT statistic | Branch ID |
|---|---|---|---|---|---|---|
| LWRh | 64 | All | 5 | 0.001, 0.001, 0.005, 0.022, 0.048 | 28.8, 23.2, 16.5, 13.6, 12.1 | Tsyl-12674, N11, N3, N50, N6 |
| LWRh | 19 | Foreground | 1 | 0.007 | 13.6591 | N50 |
| BRh | 47 | All | 2 | 0.03, 0.02 | 12.1499, 12.6127 | N8, N14 |
| BRh | 13 | Foreground | 1 | 0.018 | 11.001 | Aips & Mleu branch |
| unclassified | 31 | All | - | - | - | - |
| unclassified | 8 | Foreground | - | - | - | - |
| UVRh1 | 39 | All | - | - | - | - |
| UVRh1 | 15 | Foreground | 1 | 0.037 | 9.8874 | Node 25 |
| RGR-like | 37 | All | - | - | - | - |
| RGR-like | 13 | Foreground | - | - | - | - |
| Pteropsin | 43 | All | | | | |
| Pteropsin | 17 | Foreground | 3 | 0.001, 0.003, 0.04 | 40.9609, 14.942, 9.4199 | Msex-g8710, N10, N12 |
| Rh7 | 35 | All | 3 | 0.001, 0.003, 0.007 | 24.64, 16.15, 14.69 | Cnem, N34, N17 |
| Rh7 | 14 | Foreground | 3 | 0.001, 0.045, 0.047 | 16.153, 9.033, 9.133 | Aips-Bbet-Hpyr node34, Aips branch, Tsyl branch |

**Supplementary Table 3.** gFACs statistics of the Augustus gene prediction model.

| | |
|---|---|
| Number of genes | 19929 |
| Number of monoexonic genes | 4751 |
| Number of multiexonic genes | 15159 |
| Number of positive strand genes | 9874 |
| Monoexonic | 2294 |
| Multiexonic | 7561 |
| Number of negative strand genes | 10055 |
| Monoexonic | 2457 |
| Multiexonic | 7598 |
| Average overall gene size | 8963.3 |
| Median overall gene size | 4580 |
| Average overall CDS size | 1421.389 |
| Median overall CDS size | 996 |
| Average overall exon size | 242.253 |
| Median overall exon size | 158 |
| Average size of monoexonic genes | 878.856 |
| Median size of monoexonic genes | 654 |
| Largest monoexonic gene | 11226 |
| Smallest monoexonic gene | 93 |
| Average size of multiexonic genes | 11508.18 |
| Median size of multiexonic genes | 7614 |
| Largest multiexonic gene | 166836 |
| Smallest multiexonic gene | 172 |
| Average size of multiexonic CDS | 1593.207 |
| Median size of multiexonic CDS | 1140 |
| Largest multiexonic CDS | 58680 |
| Smallest multiexonic CDS | 21 |
| Average size of multiexonic exons | 215.292 |
| Median size of multiexonic exons | 154 |
| Average size of multiexonic introns | 1548.328 |
| Median size of multiexonic introns | 883 |
| Average number of exons per multiexonic gene | 7.4 |
| Median number of exons per multiexonic gene | 5 |
| Largest multiexonic exon | 14804 |
| Smallest multiexonic exon | 3 |
| Most exons in one gene | 150 |
| Average number of introns per multiexonic gene | 6.4 |
| Median number of introns per multiexonic gene | 4 |
| Largest intron | 31832 |
| Smallest intron | 33 |
| The following columns do not involve codons | |
| Number of complete models | 18801 |
| Number of 5' only incomplete models | 762 |
| Number of 3' only incomplete models | 311 |
| Number of 5' and 3' incomplete models | 36 |

**Supplementary Table 4.** Data used in training the BRAKER3 Augustus gene prediction model.

| Species | Genome_accession | RNAseq accession | Protein data type | Gene_model_BUSCO (lepidoptera_odb10 database, n=5286) | Braker3 pipeline |
|---|---|---|---|---|---|
| *Agrotis ipsilon* | GCA_004193855.1 | SRR15020091 | OrthoDB11 | C:96.9%[S:80.8%,D:16.1%],F:1.0%,M:2.1% | ETP |
| *Biston betularia* | GCA_905404145.2 | ERR10123640 | OrthoDB11+DToL predicted protein | C:98.7%[S:84.2%,D:14.5%],F:0.4%,M:0.9% | ETP |
| *Bombyx mori* | GCA_000151625.1 | SRR24819236 | OrthoDB11+Refeq predicted protein | C:94.7%[S:81.1%,D:13.6%],F:2.6%,M:2.7% | ETP |
| *Calephelis nemesis* | GCA_002245505.1 | NA | OrthoDB11 | C:93.0%[S:86.9%,D:6.1%],F:3.4%,M:3.6% | EP |
| *Calephelis virginiensis* | GCA_002245475.1 | NA | OrthoDB11 | C:89.7%[S:83.5%,D:6.2%],F:4.7%,M:5.6% | EP |
| *Colias croceus* | GCA_009982905.1 | ERR6054399 | OrthoDB11+DToL predicted protein | C:99.2%[S:83.3%,D:15.9%],F:0.2%,M:0.6% | ETP |
| *Danaus plexippus* | GCA_000235995.2 | SRR585568 | OrthoDB11 | C:96.9%[S:78.5%,D:18.4%],F:1.7%,M:1.4% | ETP |
| *Habrosyne pyritoides* | GCA_907165245.1 | ERR9434975 | OrthoDB11+DToL predicted protein | C:98.6%[S:82.8%,D:15.8%],F:0.3%,M:1.1% | ETP |
| *Heliconius melpomene* | GCA_000313835.2 | ERR2206056 | OrthoDB11 | C:94.1%[S:81.9%,D:12.2%],F:1.9%,M:4.0% | ETP |
| *Lycaena phlaeas* | GCA_905333005.2 | ERR3169698 | OrthoDB11+DToL predicted protein | C:98.9%[S:85.9%,D:13.0%],F:0.2%,M:0.9% | ETP |
| *Lysandra coridon* | GCA_905220515.1 | ERR6286713 | OrthoDB11 | C:97.6%[S:84.3%,D:13.3%],F:0.4%,M:2.0% | ETP |
| *Macrosoma leucophasiata* | NA | NA | OrthoDB11 | C:92.9%[S:83.3%,D:9.6%],F:1.1%,M:6.0% | EP |
| *Manduca sexta* | GCA_000262585.1 | SRR1577001, SRR1577000, SRR1576999 | OrthoDB11 | C:96.6%[S:79.7%,D:16.9%],F:1.6%,M:1.8% | ETP |
| *Papilio machaon* | GCA_001298355.1 | NA | OrthoDB11+DToL predicted protein | C:98.4%[S:87.0%,D:11.4%],F:0.2%,M:1.4% | EP |
| *Papilio xuthus* | GCA_000836235.2 | NA | OrthoDB11+Refeq predicted protein | C:96.9%[S:82.6%,D:14.3%],F:1.3%,M:1.8% | ETP |
| *Plodia interpunctella* | GCA_001368715.1 | SRR6002827 | OrthoDB11+DToL predicted protein | C:96.3%[S:80.9%,D:15.4%],F:1.4%,M:2.3% | ETP |
| *Pyrgus malvae* | GCA_911387765.1 | ERR6363277 | OrthoDB11+DToL predicted protein | C:98.8%[S:83.6%,D:15.2%],F:0.4%,M:0.8% | ETP |
| *Sesia bembeciformis* | GCA_943735995.1 | ERR10123699 | OrthoDB11+DToL predicted protein | C:98.7%[S:84.2%,D:14.5%],F:0.6%,M:0.7% | ETP |
| *Thymelicus sylvestris* | GCA_911387775.1 | ERR6363314 | OrthoDB11+DToL predicted protein | C:98.9%[S:83.7%,D:15.2%],F:0.5%,M:0.6% | ETP |
| *Zygaena filipendulae* | GCA_907165275.2 | ERR9434973 | OrthoDB11+DToL predicted protein | C:97.6%[S:82.9%,D:14.7%],F:0.5%,M:1.9% | ETP |

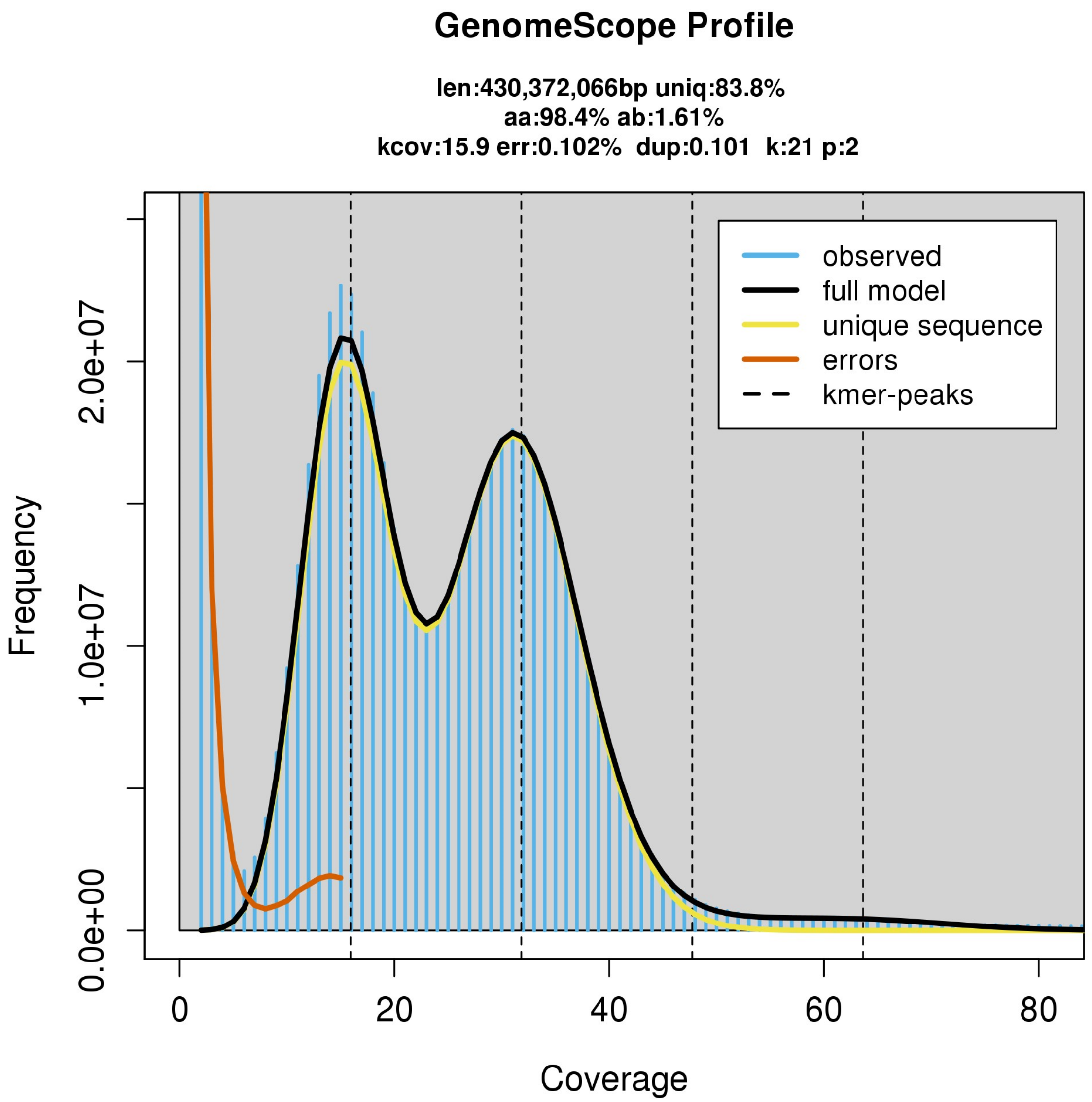


**Supplementary Figure 2.** K-mer distribution calculated using KMC with k-mer size of 21 bps, the fitting model (black line) is created using GenomeScope2.

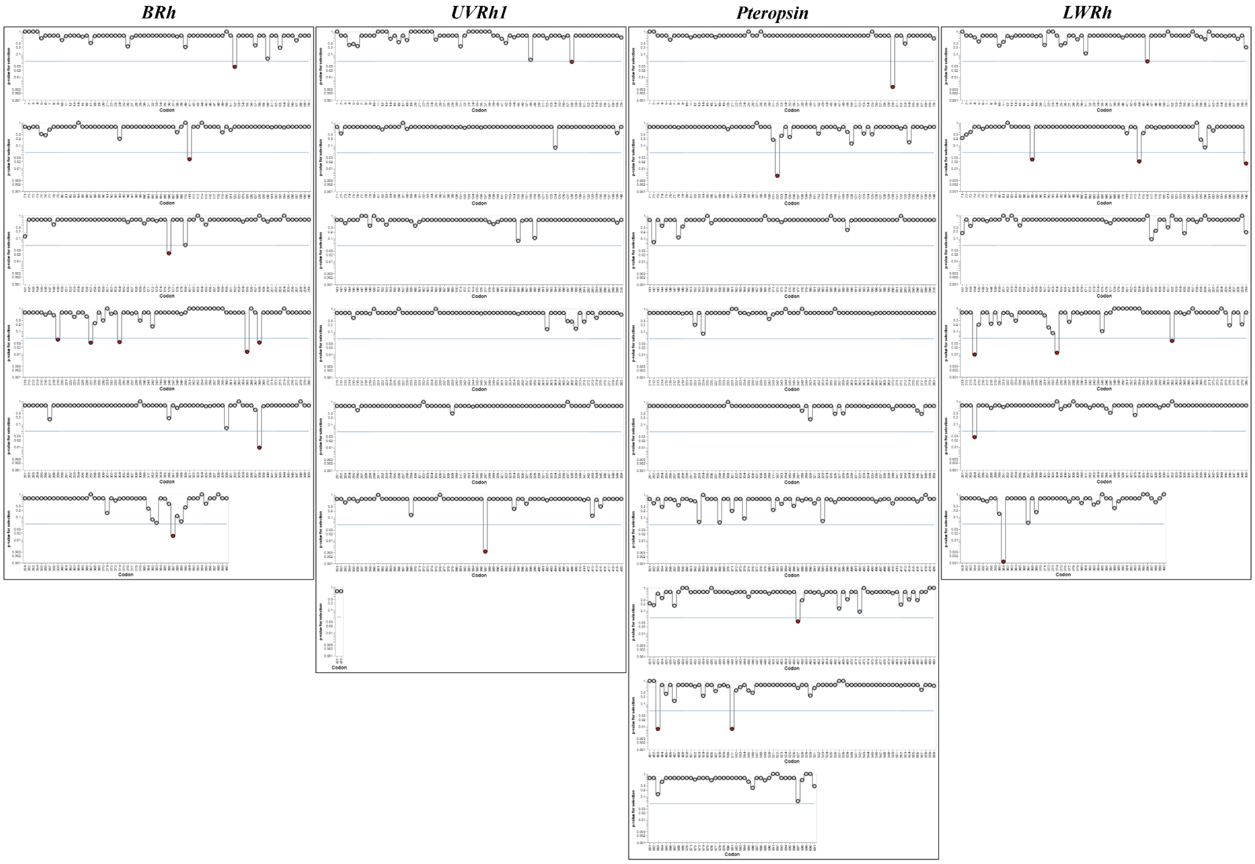


**Supplementary Figure 3.** Episodic diversifying selection of the four codons at the *M. leucophasiata* branch based on the result of selection analysis of site-substitution model, MEME. Detailed statistics are provided in **Supplementary Data 12**.

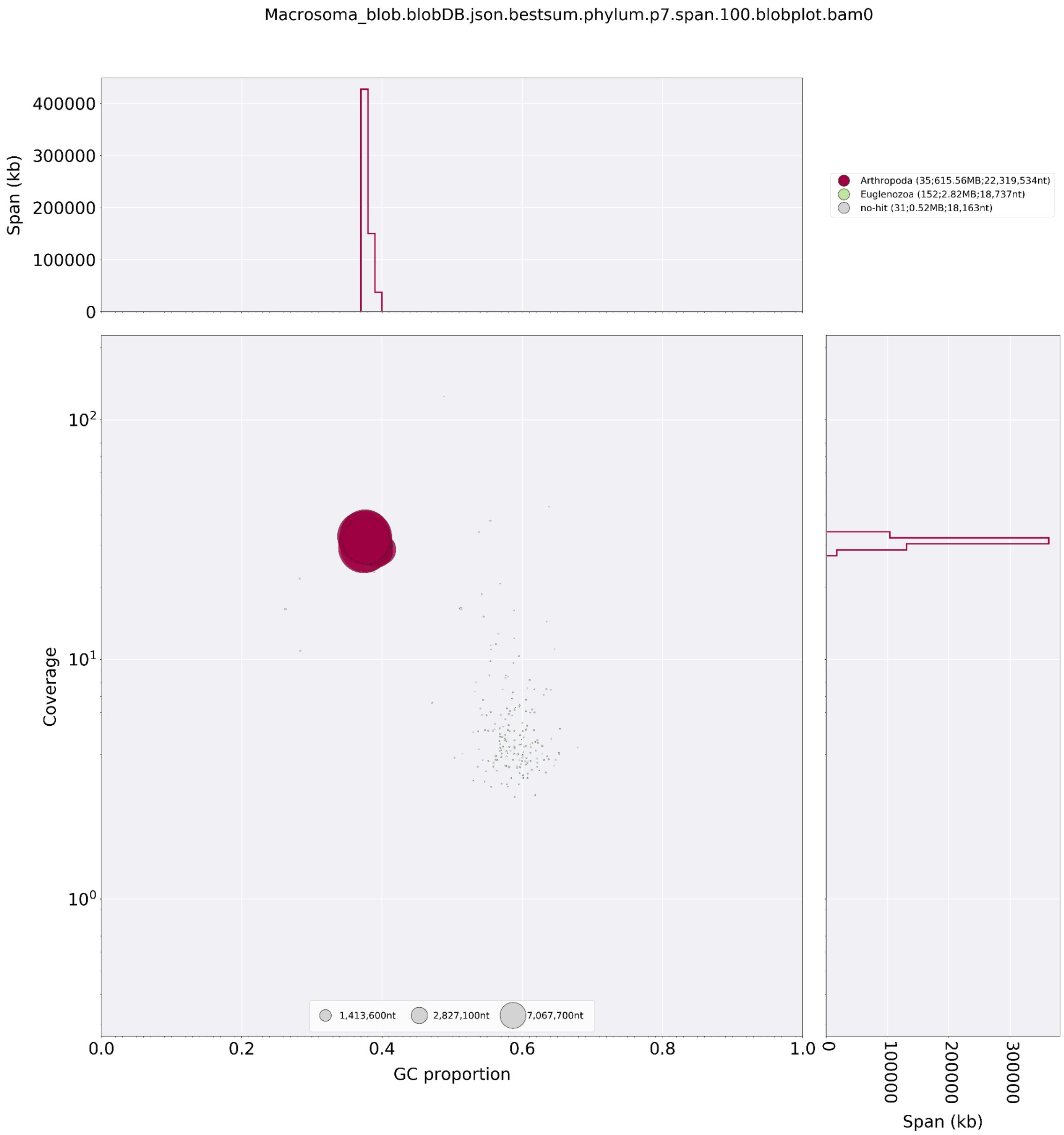


**Supplementary Figure 4.** BlobPlot of the *M. leucophasiata* purged genome assembly. Red dots show contigs with best blast hits to Arthropoda, the 152 small green dots were blasted to Euglenozoa, and 31 gray dots had no hits. We removed 152 contigs that blasted to Euglenozoa (2.82 Mb in total) from the assembly.

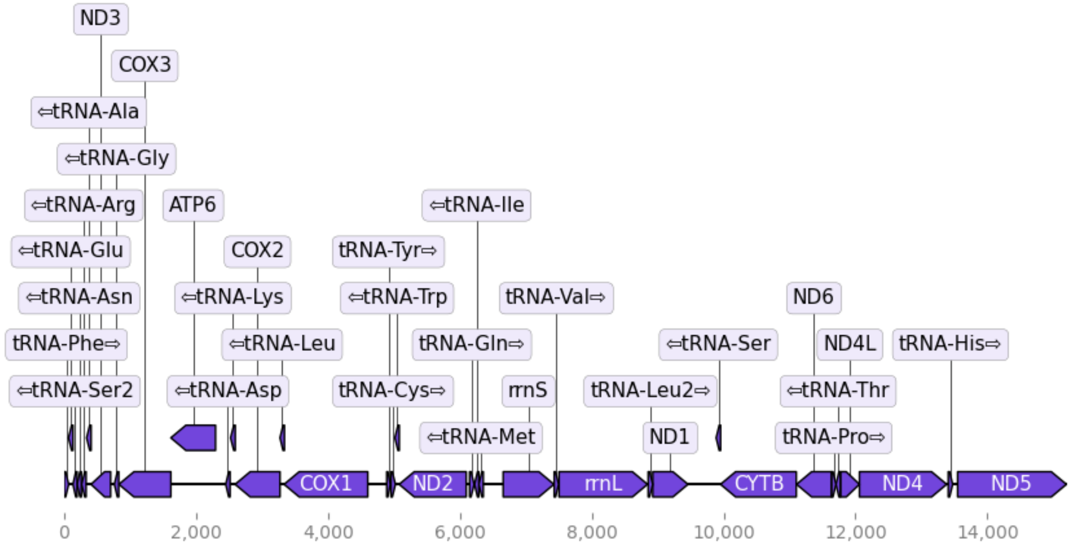


**Supplementary Figure 5.** Graphical annotation results of the *M. leucophasiata* mitochondrial genome.

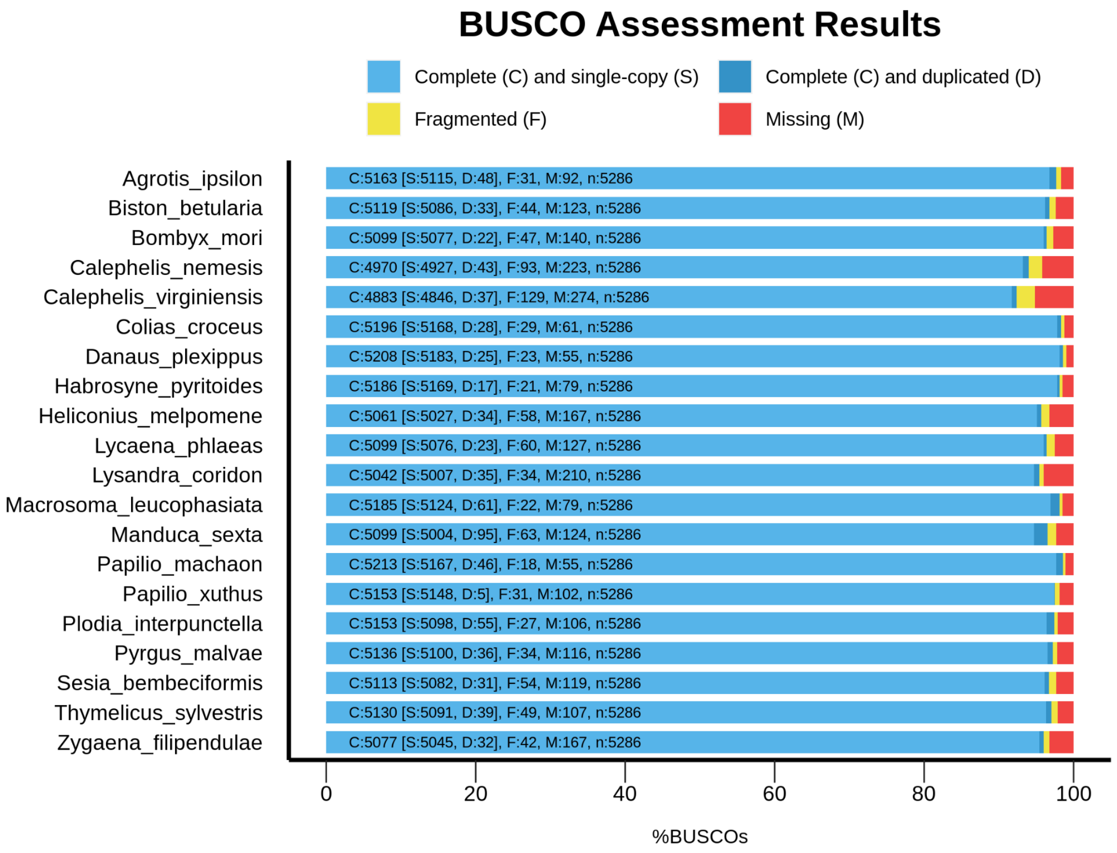


**Supplementary Figure 6.** BUSCO scores of selected species using the lepidoptera_odb10 database.
