## Supplementary Data 1: Hifiasm assembly QUAST report for "New genome reveals molecular signatures of adaptation to nocturnality in moth-like butterflies (Hedylidae)"

| | Macrosoma_default.asm |
|---|---|
| # contigs (>= 0 bp) | 397 |
| # contigs (>= 1000 bp) | 397 |
| # contigs (>= 5000 bp) | 393 |
| # contigs (>= 10000 bp) | 363 |
| # contigs (>= 25000 bp) | 77 |
| # contigs (>= 50000 bp) | 43 |
| Total length (>= 0 bp) | 622866406 |
| Total length (>= 1000 bp) | 622866406 |
| Total length (>= 5000 bp) | 622849610 |
| Total length (>= 10000 bp) | 622619556 |
| Total length (>= 25000 bp) | 617820304 |
| Total length (>= 50000 bp) | 616765004 |
| # contigs | 397 |
| Largest contig | 28270568 |
| Total length | 622866406 |
| GC (%) | 38.11 |
| N50 | 22319534 |
| N90 | 12667486 |
| auN | 20754773.5 |
| L50 | 13 |
| L90 | 27 |
| # N's per 100 kbp | 0.00 |

All statistics are based on contigs of size >= 500 bp, unless otherwise noted (e.g., "# contigs (>= 0 bp)" and "Total length (>= 0 bp)" include all contigs).

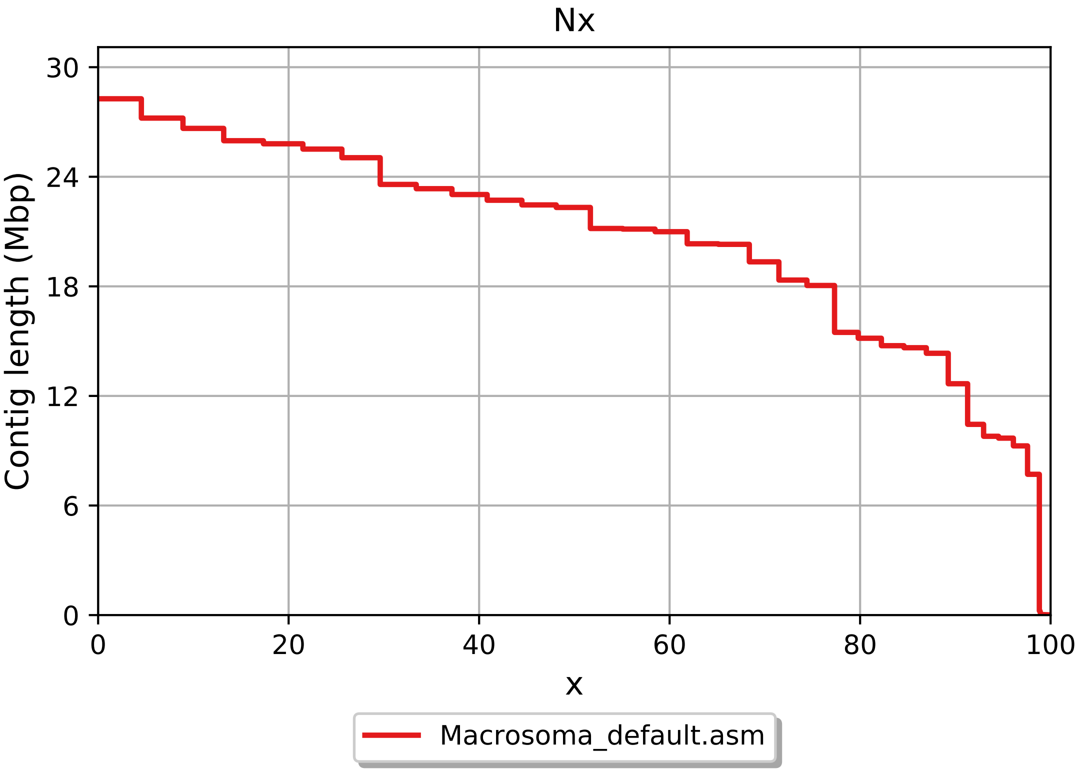
Nx
Contig length (Mbp)
0
6
12
18
24
30
0
20
40
60
80
100
x
Macrosoma_default.asm

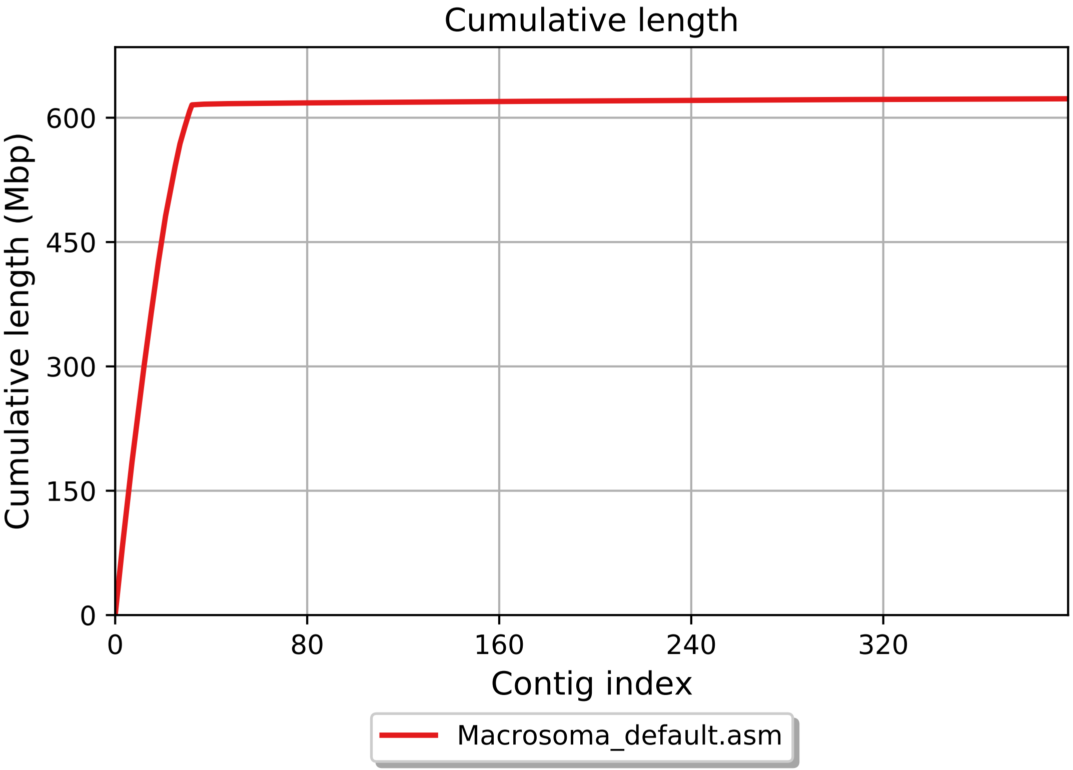

Cumulative length
Cumulative length (Mbp)
0
150
300
450
600
0
80
160
240
320
Contig index
Macrosoma_default.asm

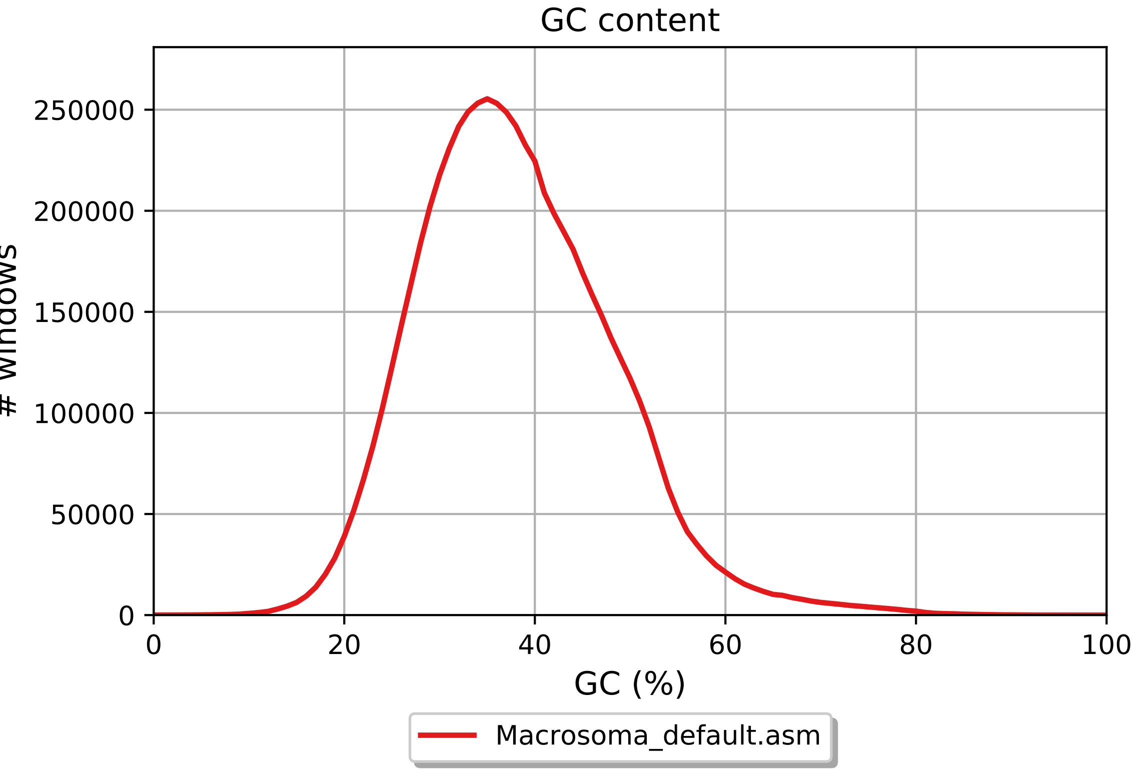
GC content
250000
200000
150000
100000
50000
0
### windows
0
20
40
60
80
100
GC (%)
Macrosoma_default.asm

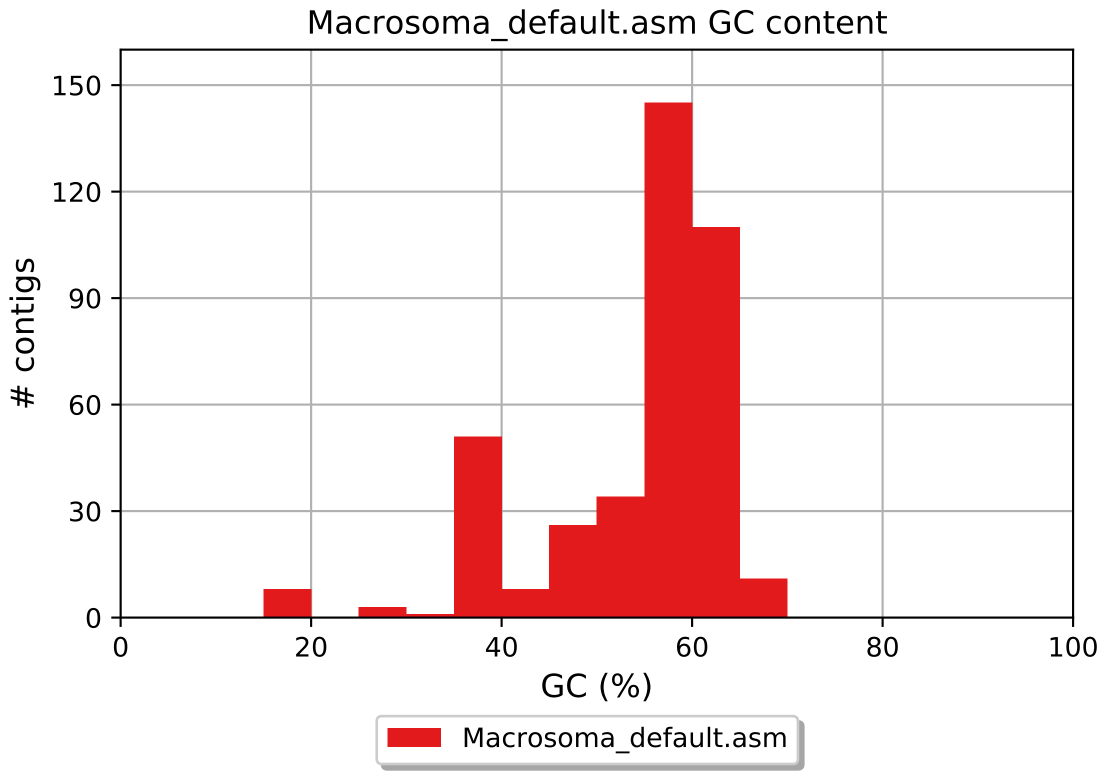
Macrosoma_default.asm GC content
150
120
90
60
30
0
### contigs
0
20
40
60
80
100
GC (%)
Macrosoma_default.asm
