## Supplementary Data 2: Final soft-masked assembly QUAST report for "New genome reveals molecular signatures of adaptation to nocturnality in moth-like butterflies (Hedylidae)"

| | Macrosoma_final_assembly |
|---|---|
| # contigs (>= 0 bp) | 66 |
| # contigs (>= 1000 bp) | 66 |
| # contigs (>= 5000 bp) | 65 |
| # contigs (>= 10000 bp) | 65 |
| # contigs (>= 25000 bp) | 34 |
| # contigs (>= 50000 bp) | 32 |
| Total length (>= 0 bp) | 616085051 |
| Total length (>= 1000 bp) | 616085051 |
| Total length (>= 5000 bp) | 616081851 |
| Total length (>= 10000 bp) | 616081851 |
| Total length (>= 25000 bp) | 615561551 |
| Total length (>= 50000 bp) | 615506505 |
| # contigs | 66 |
| Largest contig | 28270568 |
| Total length | 616085051 |
| GC (%) | 37.94 |
| N50 | 22319534 |
| N90 | 14335051 |
| auN | 20982743.1 |
| L50 | 13 |
| L90 | 26 |
| # N's per 100 kbp | 0.00 |

All statistics are based on contigs of size >= 500 bp, unless otherwise noted (e.g., "# contigs (>= 0 bp)" and "Total length (>= 0 bp)" include all contigs).

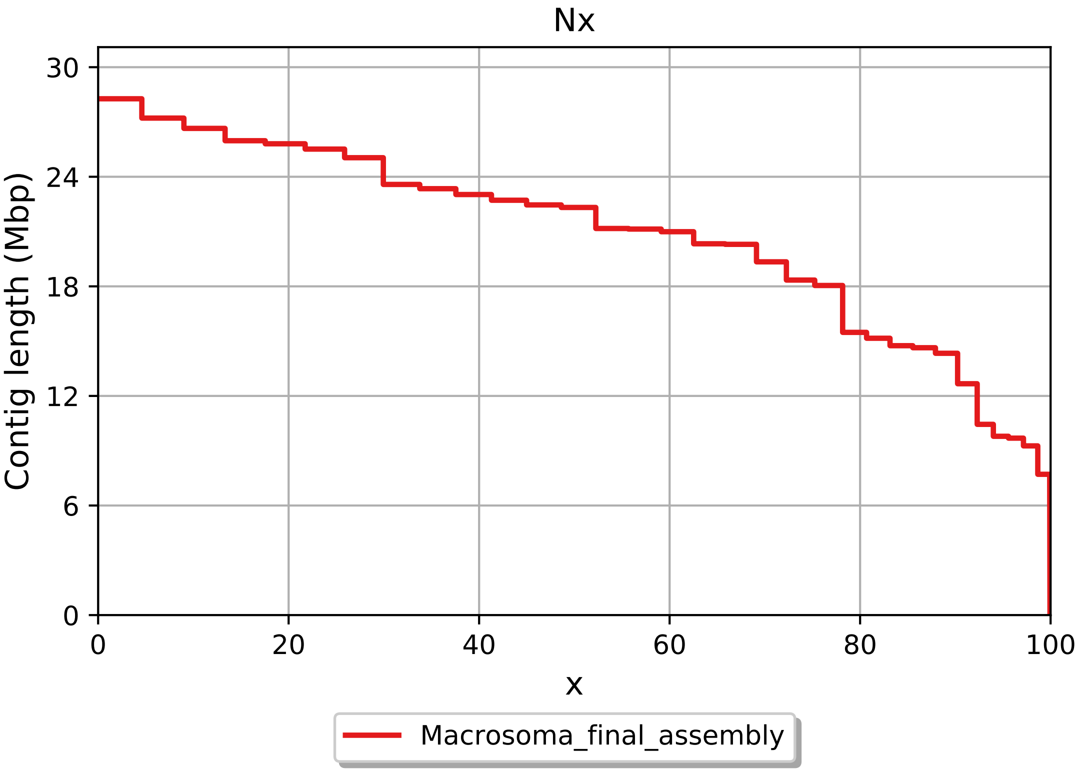
Nx
30
24
18
12
6
0
0
20
40
60
80
100
x
Contig length (Mbp)
Macrosoma_final_assembly

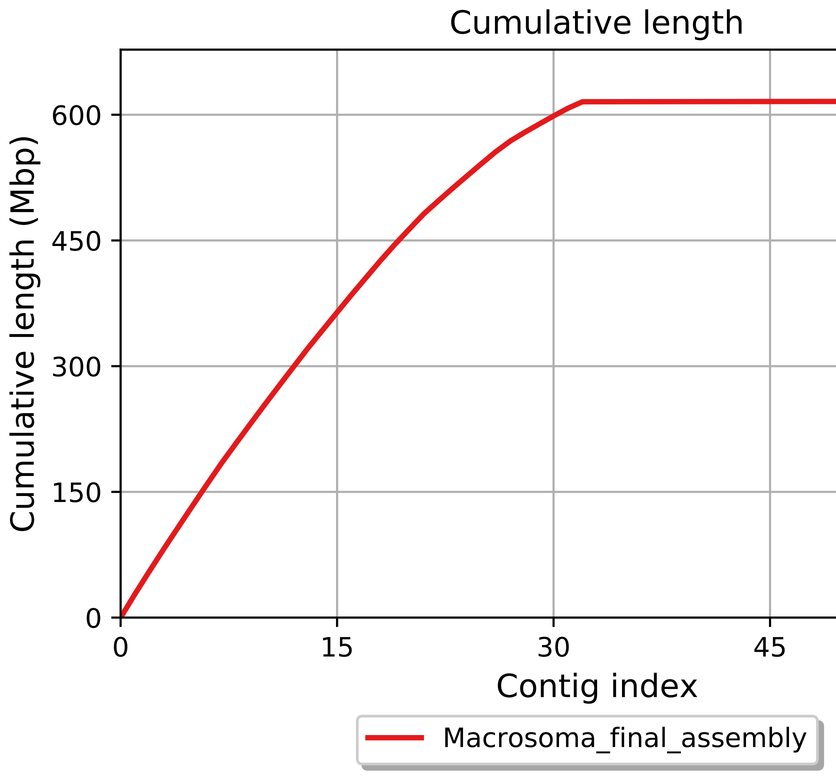

Cumulative length
Cumulative length (Mbp)
0
150
300
450
600
0
15
30
45
60
Contig index
Macrosoma_final_assembly

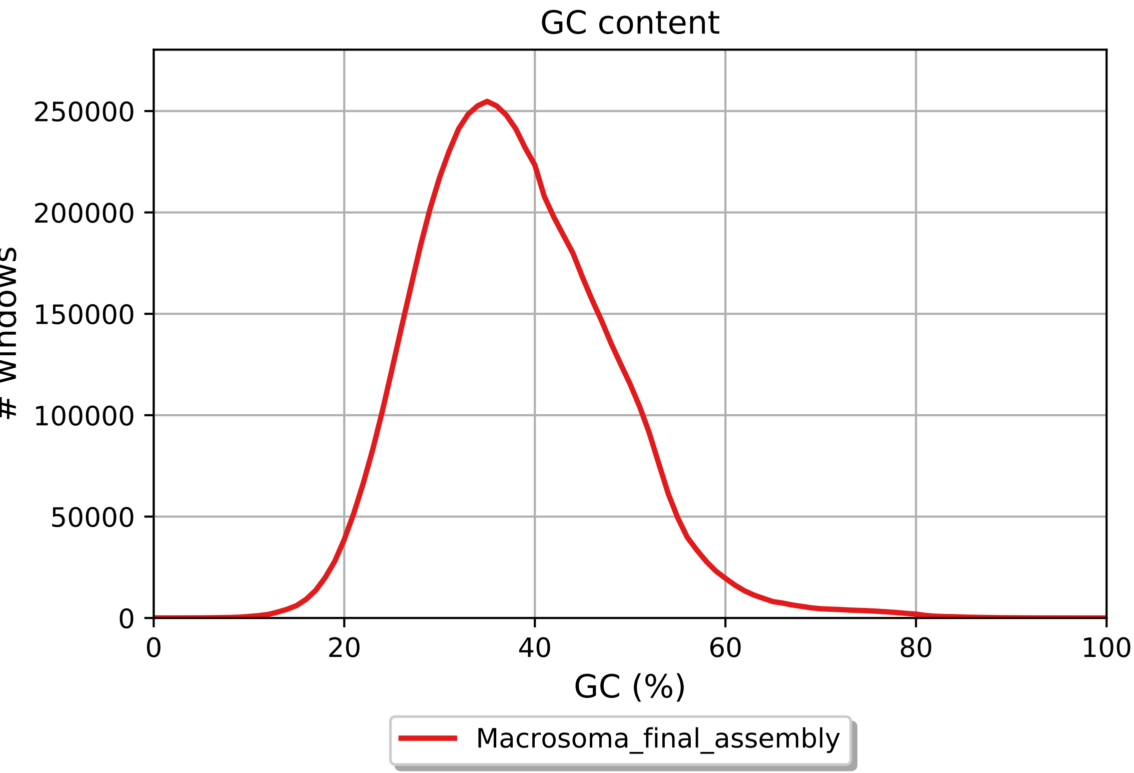
GC content
250000
200000
150000
100000
50000
0
### windows
0
20
40
60
80
100
GC (%)
Macrosoma_final_assembly

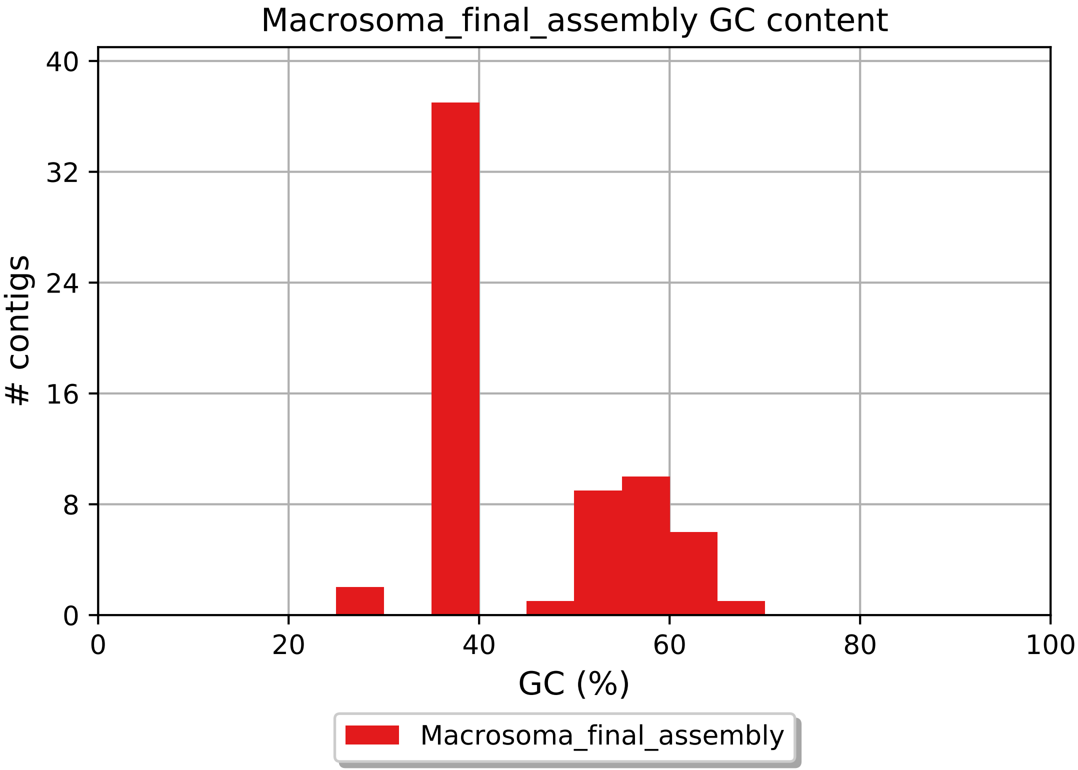

Macrosoma_final_assembly GC content
### contigs
40
32
24
16
8
0
0
20
40
60
80
100
GC (%)
Macrosoma_final_assembly
