## Supplementary Data 10: All (32) vision gene family trees for "New genome reveals molecular signatures of adaptation to nocturnality in moth-like butterflies (Hedylidae)"

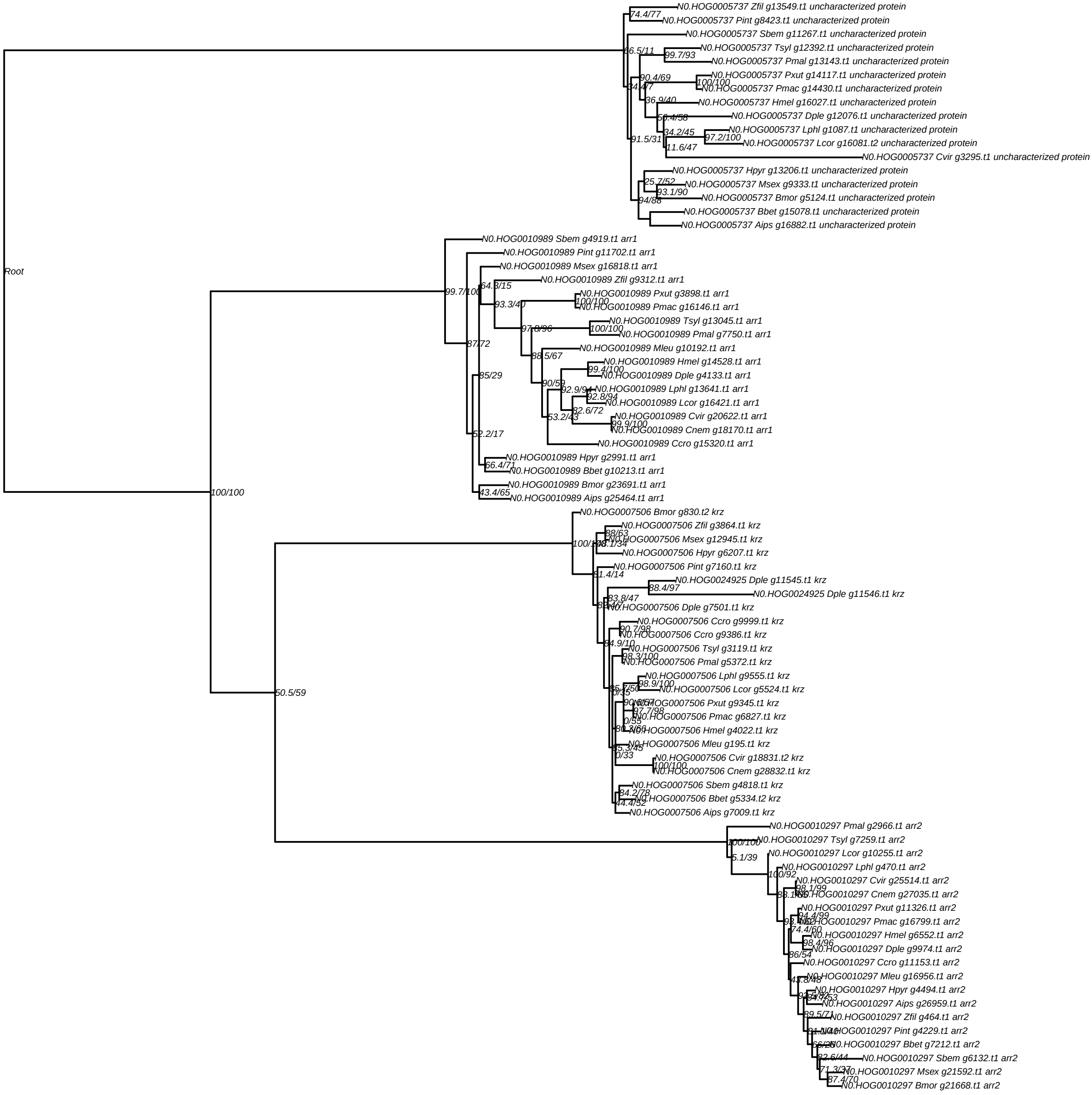
Root
100/100
50.5/59
N0.HOG0005737 Zfil g13549.t1 uncharacterized protein
74.4/77
N0.HOG0005737 Pint g8423.t1 uncharacterized protein
N0.HOG0005737 Sbem g11267.t1 uncharacterized protein
66.5/11
N0.HOG0005737 Tsyl g12392.t1 uncharacterized protein
99.7/93
N0.HOG0005737 Pmal g13143.t1 uncharacterized protein
90.4/69
N0.HOG0005737 Pxut g14117.t1 uncharacterized protein
100/100
N0.HOG0005737 Pmac g14430.t1 uncharacterized protein
36.9/40
N0.HOG0005737 Hmel g16027.t1 uncharacterized protein
N0.HOG0005737 Dple g12076.t1 uncharacterized protein
34.2/45
N0.HOG0005737 Lphl g1087.t1 uncharacterized protein
97.2/100
N0.HOG0005737 Lcor g16081.t2 uncharacterized protein
11.6/47
91.5/31
N0.HOG0005737 Cvir g3295.t1 uncharacterized protein
N0.HOG0005737 Hpyr g13206.t1 uncharacterized protein
25.7/52
N0.HOG0005737 Msex g9333.t1 uncharacterized protein
93.1/90
N0.HOG0005737 Bmor g5124.t1 uncharacterized protein
94/88
N0.HOG0005737 Bbet g15078.t1 uncharacterized protein
N0.HOG0005737 Aips g16882.t1 uncharacterized protein
N0.HOG0010989 Sbem g4919.t1 arr1
N0.HOG0010989 Pint g11702.t1 arr1
N0.HOG0010989 Msex g16818.t1 arr1
99.7/100
N0.HOG0010989 Zfil g9312.t1 arr1
N0.HOG0010989 Pxut g3898.t1 arr1
100/100
93.3/40
N0.HOG0010989 Pmac g16146.t1 arr1
N0.HOG0010989 Tsyl g13045.t1 arr1
97.8/96
100/100
N0.HOG0010989 Pmal g7750.t1 arr1
87/72
N0.HOG0010989 Mleu g10192.t1 arr1
88.5/67
N0.HOG0010989 Hmel g14528.t1 arr1
99.4/100
N0.HOG0010989 Dple g4133.t1 arr1
85/29
90/59
N0.HOG0010989 Lphl g13641.t1 arr1
92.9/94
92.8/94
N0.HOG0010989 Lcor g16421.t1 arr1
82.6/72
N0.HOG0010989 Cvir g20622.t1 arr1
53.2/43
99.9/100
N0.HOG0010989 Cnem g18170.t1 arr1
N0.HOG0010989 Ccro g15320.t1 arr1
52.2/17
N0.HOG0010989 Hpyr g2991.t1 arr1
66.4/71
N0.HOG0010989 Bbet g10213.t1 arr1
N0.HOG0010989 Bmor g23691.t1 arr1
43.4/65
N0.HOG0010989 Aips g25464.t1 arr1
N0.HOG0007506 Bmor g830.t2 krz
N0.HOG0007506 Zfil g3864.t1 krz
88/63
N0.HOG0007506 Msex g12945.t1 krz
N0.HOG0007506 Hpyr g6207.t1 krz
N0.HOG0007506 Pint g7160.t1 krz
81.4/14
N0.HOG0024925 Dple g11545.t1 krz
88.4/97
N0.HOG0024925 Dple g11546.t1 krz
83.8/47
N0.HOG0007506 Dple g7501.t1 krz
N0.HOG0007506 Ccro g9999.t1 krz
90.7/98
N0.HOG0007506 Ccro g9386.t1 krz
84.9/10
N0.HOG0007506 Tsyl g3119.t1 krz
98.3/100
N0.HOG0007506 Pmal g5372.t1 krz
N0.HOG0007506 Lphl g9555.t1 krz
98.9/100
N0.HOG0007506 Lcor g5524.t1 krz
0/35
N0.HOG0007506 Pxut g9345.t1 krz
97.7/98
N0.HOG0007506 Pmac g6827.t1 krz
0/55
N0.HOG0007506 Hmel g4022.t1 krz
N0.HOG0007506 Mleu g195.t1 krz
0/33
N0.HOG0007506 Cvir g18831.t2 krz
100/100
N0.HOG0007506 Cnem g28832.t1 krz
N0.HOG0007506 Sbem g4818.t1 krz
84.2/78
N0.HOG0007506 Bbet g5334.t2 krz
44.4/52
N0.HOG0007506 Aips g7009.t1 krz
N0.HOG0010297 Pmal g2966.t1 arr2
100/100
N0.HOG0010297 Tsyl g7259.t1 arr2
5.1/39
N0.HOG0010297 Lcor g10255.t1 arr2
100/92
N0.HOG0010297 Lphl g470.t1 arr2
N0.HOG0010297 Cvir g25514.t1 arr2
98.1/99
N0.HOG0010297 Cnem g27035.t1 arr2
N0.HOG0010297 Pxut g11326.t1 arr2
94.4/99
N0.HOG0010297 Pmac g16799.t1 arr2
74.4/60
N0.HOG0010297 Hmel g6552.t1 arr2
98.4/96
N0.HOG0010297 Dple g9974.t1 arr2
86/54
N0.HOG0010297 Ccro g11153.t1 arr2
N0.HOG0010297 Mleu g16956.t1 arr2
43.8/48
N0.HOG0010297 Hpyr g4494.t1 arr2
N0.HOG0010297 Aips g26959.t1 arr2
89.5/71
N0.HOG0010297 Zfil g464.t1 arr2
N0.HOG0010297 Pint g4229.t1 arr2
N0.HOG0010297 Bbet g7212.t1 arr2
82.6/44
N0.HOG0010297 Sbem g6132.t1 arr2
71.3/37
N0.HOG0010297 Msex g21592.t1 arr2
87.4/70
N0.HOG0010297 Bmor g21668.t1 arr2

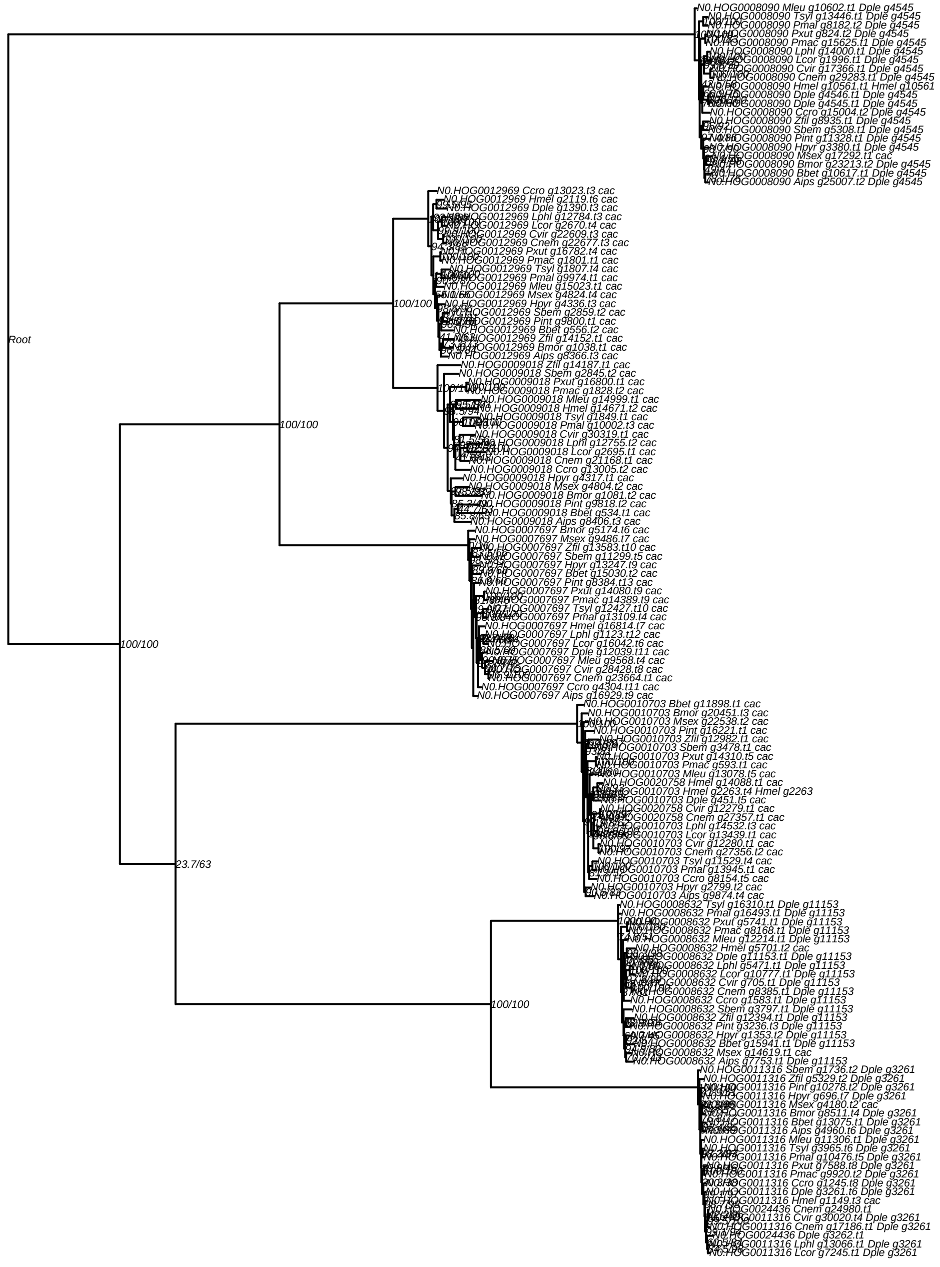

Root
100/100
100/100
100/100
100/100
23.7/63
100/100
N0.HOG0008090 Mleu g10602.t1 Dple g4545
N0.HOG0008090 Tsyl g13446.t1 Dple g4545
N0.HOG0008090 Pmal g8182.t2 Dple g4545
N0.HOG0008090 Pxut g824.t2 Dple g4545
N0.HOG0008090 Pmac g15625.t1 Dple g4545
N0.HOG0008090 Lphl g14000.t1 Dple g4545
N0.HOG0008090 Lcor g1996.t1 Dple g4545
N0.HOG0008090 Cvir g17366.t1 Dple g4545
N0.HOG0008090 Cnem g29283.t1 Dple g4545
N0.HOG0008090 Hmel g10561.t1 Hmel g10561
N0.HOG0008090 Dple g4546.t1 Dple g4545
N0.HOG0008090 Dple g4545.t1 Dple g4545
N0.HOG0008090 Ccro g15004.t2 Dple g4545
N0.HOG0008090 Zfil g8935.t1 Dple g4545
N0.HOG0008090 Sbem g5308.t1 Dple g4545
N0.HOG0008090 Pint g11328.t1 Dple g4545
N0.HOG0008090 Hpyr g3380.t1 Dple g4545
N0.HOG0008090 Msex g17292.t1 cac
N0.HOG0008090 Bmor g23213.t2 Dple g4545
N0.HOG0008090 Bbet g10617.t1 Dple g4545
N0.HOG0008090 Aips g25007.t2 Dple g4545
N0.HOG0012969 Ccro g13023.t3 cac
N0.HOG0012969 Hmel g2119.t6 cac
N0.HOG0012969 Dple g1390.t3 cac
N0.HOG0012969 Lphl g12784.t3 cac
N0.HOG0012969 Lcor g2670.t4 cac
N0.HOG0012969 Cvir g22609.t3 cac
N0.HOG0012969 Cnem g22677.t3 cac
N0.HOG0012969 Pxut g16782.t4 cac
N0.HOG0012969 Pmac g1801.t1 cac
N0.HOG0012969 Tsyl g1807.t4 cac
N0.HOG0012969 Pmal g9974.t1 cac
N0.HOG0012969 Mleu g15023.t1 cac
N0.HOG0012969 Msex g4824.t4 cac
N0.HOG0012969 Hpyr g4336.t3 cac
N0.HOG0012969 Sbem g2859.t2 cac
N0.HOG0012969 Pint g9800.t1 cac
N0.HOG0012969 Bbet g556.t2 cac
N0.HOG0012969 Zfil g14152.t1 cac
N0.HOG0012969 Bmor g1038.t1 cac
N0.HOG0012969 Aips g8366.t3 cac
N0.HOG0009018 Zfil g14187.t1 cac
N0.HOG0009018 Sbem g2845.t2 cac
N0.HOG0009018 Pxut g16800.t1 cac
N0.HOG0009018 Pmac g1828.t2 cac
N0.HOG0009018 Mleu g14999.t1 cac
N0.HOG0009018 Hmel g14671.t2 cac
N0.HOG0009018 Tsyl g1849.t1 cac
N0.HOG0009018 Pmal g10002.t3 cac
N0.HOG0009018 Cvir g30319.t1 cac
N0.HOG0009018 Lphl g12755.t2 cac
N0.HOG0009018 Lcor g2695.t1 cac
N0.HOG0009018 Cnem g21168.t1 cac
N0.HOG0009018 Ccro g13005.t2 cac
N0.HOG0009018 Hpyr g4317.t1 cac
N0.HOG0009018 Msex g4804.t2 cac
N0.HOG0009018 Bmor g1081.t2 cac
N0.HOG0009018 Pint g9818.t2 cac
N0.HOG0009018 Bbet g534.t1 cac
N0.HOG0009018 Aips g8406.t3 cac
N0.HOG0007697 Bmor g5174.t6 cac
N0.HOG0007697 Msex g9486.t7 cac
N0.HOG0007697 Zfil g13583.t10 cac
N0.HOG0007697 Sbem g11299.t5 cac
N0.HOG0007697 Hpyr g13247.t9 cac
N0.HOG0007697 Bbet g15030.t2 cac
N0.HOG0007697 Pint g8384.t13 cac
N0.HOG0007697 Pxut g14080.t9 cac
N0.HOG0007697 Pmac g14389.t9 cac
N0.HOG0007697 Tsyl g12427.t10 cac
N0.HOG0007697 Pmal g13109.t4 cac
N0.HOG0007697 Hmel g16814.t7 cac
N0.HOG0007697 Lphl g1123.t12 cac
N0.HOG0007697 Lcor g16042.t6 cac
N0.HOG0007697 Dple g12039.t11 cac
N0.HOG0007697 Mleu g9568.t4 cac
N0.HOG0007697 Cvir g28428.t8 cac
N0.HOG0007697 Cnem g23664.t1 cac
N0.HOG0007697 Ccro g4304.t11 cac
N0.HOG0007697 Aips g16929.t9 cac
N0.HOG0010703 Bbet g11898.t1 cac
N0.HOG0010703 Bmor g20451.t3 cac
N0.HOG0010703 Msex g22538.t2 cac
N0.HOG0010703 Pint g16221.t1 cac
N0.HOG0010703 Zfil g12982.t1 cac
N0.HOG0010703 Sbem g3478.t1 cac
N0.HOG0010703 Pxut g14310.t5 cac
N0.HOG0010703 Pmac g593.t1 cac
N0.HOG0010703 Mleu g13078.t5 cac
N0.HOG0020758 Hmel g14088.t1 cac
N0.HOG0010703 Hmel g2263.t4 Hmel g2263
N0.HOG0010703 Dple g451.t5 cac
N0.HOG0020758 Cvir g12279.t1 cac
N0.HOG0020758 Cnem g27357.t1 cac
N0.HOG0010703 Lphl g14532.t3 cac
N0.HOG0010703 Lcor g13439.t1 cac
N0.HOG0010703 Cvir g12280.t1 cac
N0.HOG0010703 Cnem g27356.t2 cac
N0.HOG0010703 Tsyl g11529.t4 cac
N0.HOG0010703 Pmal g13945.t1 cac
N0.HOG0010703 Ccro g8154.t5 cac
N0.HOG0010703 Hpyr g2799.t2 cac
N0.HOG0010703 Aips g9874.t4 cac
N0.HOG0008632 Tsyl g16310.t1 Dple g11153
N0.HOG0008632 Pmal g16493.t1 Dple g11153
N0.HOG0008632 Pxut g5741.t1 Dple g11153
N0.HOG0008632 Pmac g8168.t1 Dple g11153
N0.HOG0008632 Mleu g12214.t1 Dple g11153
N0.HOG0008632 Hmel g5701.t2 cac
N0.HOG0008632 Dple g11153.t1 Dple g11153
N0.HOG0008632 Lphl g5471.t1 Dple g11153
N0.HOG0008632 Lcor g10777.t1 Dple g11153
N0.HOG0008632 Cvir g705.t1 Dple g11153
N0.HOG0008632 Cnem g8385.t1 Dple g11153
N0.HOG0008632 Ccro g1583.t1 Dple g11153
N0.HOG0008632 Sbem g3797.t1 Dple g11153
N0.HOG0008632 Zfil g12394.t1 Dple g11153
N0.HOG0008632 Pint g3236.t3 Dple g11153
N0.HOG0008632 Hpyr g1353.t2 Dple g11153
N0.HOG0008632 Bbet g15941.t1 Dple g11153
N0.HOG0008632 Msex g14619.t1 cac
N0.HOG0008632 Aips g7753.t1 Dple g11153
N0.HOG0011316 Sbem g1736.t2 Dple g3261
N0.HOG0011316 Zfil g5329.t2 Dple g3261
N0.HOG0011316 Pint g10278.t2 Dple g3261
N0.HOG0011316 Hpyr g696.t7 Dple g3261
N0.HOG0011316 Msex g4180.t2 cac
N0.HOG0011316 Bmor g8511.t4 Dple g3261
N0.HOG0011316 Bbet g13075.t1 Dple g3261
N0.HOG0011316 Aips g4960.t6 Dple g3261
N0.HOG0011316 Mleu g11306.t1 Dple g3261
N0.HOG0011316 Tsyl g3965.t6 Dple g3261
N0.HOG0011316 Pmal g10476.t5 Dple g3261
N0.HOG0011316 Pxut g7588.t8 Dple g3261
N0.HOG0011316 Pmac g9920.t2 Dple g3261
N0.HOG0011316 Ccro g1245.t8 Dple g3261
N0.HOG0011316 Dple g3261.t6 Dple g3261
N0.HOG0011316 Hmel g1149.t3 cac
N0.HOG0024436 Cnem g24980.t1
N0.HOG0011316 Cvir g30020.t4 Dple g3261
N0.HOG0011316 Cnem g17186.t1 Dple g3261
N0.HOG0024436 Dple g3262.t1
N0.HOG0011316 Lphl g13066.t1 Dple g3261
N0.HOG0011316 Lcor g7245.t1 Dple g3261

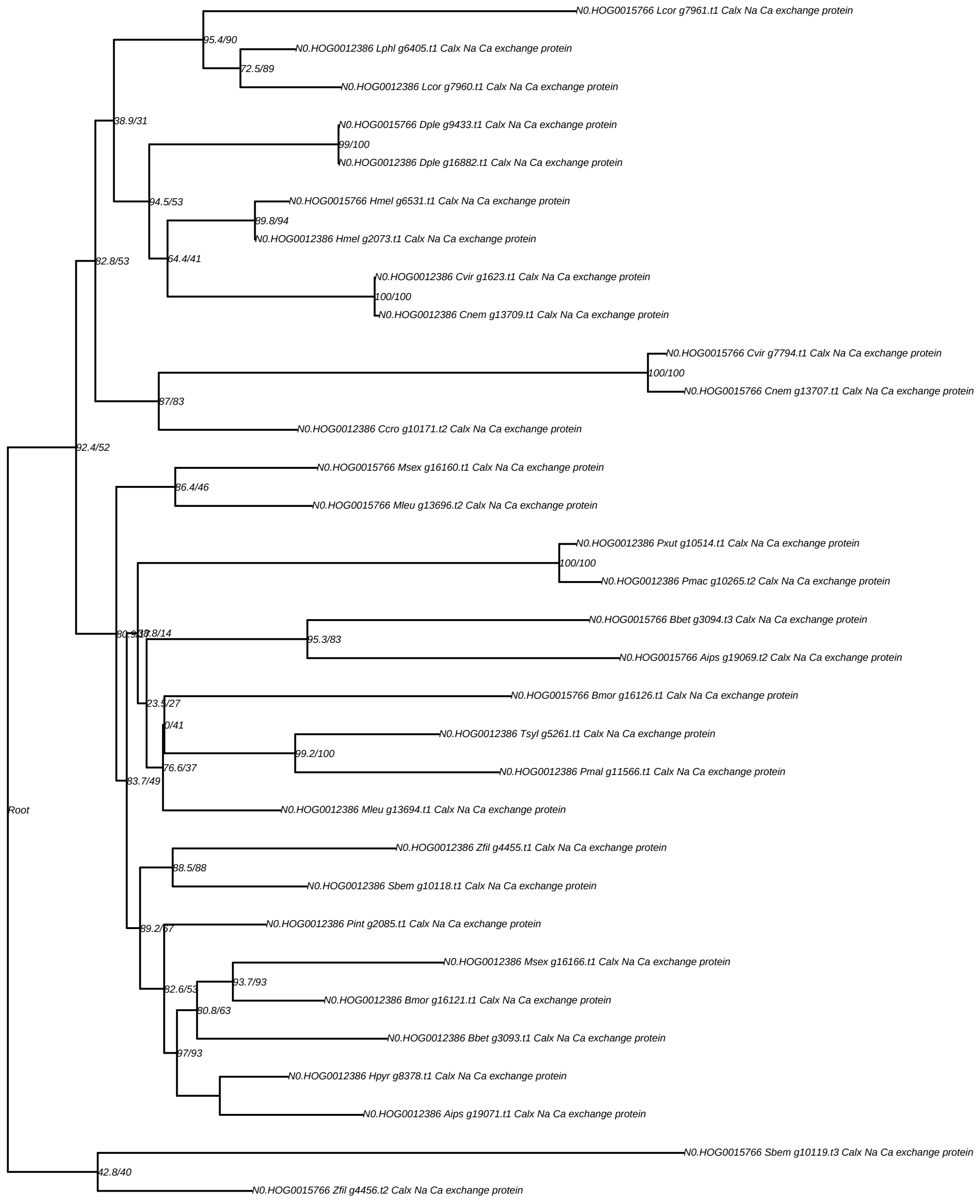
N0.HOG0015766 Lcor g7961.t1 Calx Na Ca exchange protein
95.4/90
N0.HOG0012386 Lphl g6405.t1 Calx Na Ca exchange protein
72.5/89
N0.HOG0012386 Lcor g7960.t1 Calx Na Ca exchange protein
38.9/31
N0.HOG0015766 Dple g9433.t1 Calx Na Ca exchange protein
99/100
N0.HOG0012386 Dple g16882.t1 Calx Na Ca exchange protein
94.5/53
N0.HOG0015766 Hmel g6531.t1 Calx Na Ca exchange protein
89.8/94
N0.HOG0012386 Hmel g2073.t1 Calx Na Ca exchange protein
82.8/53
64.4/41
N0.HOG0012386 Cvir g1623.t1 Calx Na Ca exchange protein
100/100
N0.HOG0012386 Cnem g13709.t1 Calx Na Ca exchange protein
N0.HOG0015766 Cvir g7794.t1 Calx Na Ca exchange protein
100/100
N0.HOG0015766 Cnem g13707.t1 Calx Na Ca exchange protein
87/83
N0.HOG0012386 Ccro g10171.t2 Calx Na Ca exchange protein
92.4/52
N0.HOG0015766 Msex g16160.t1 Calx Na Ca exchange protein
86.4/46
N0.HOG0015766 Mleu g13696.t2 Calx Na Ca exchange protein
N0.HOG0012386 Pxut g10514.t1 Calx Na Ca exchange protein
100/100
N0.HOG0012386 Pmac g10265.t2 Calx Na Ca exchange protein
N0.HOG0015766 Bbet g3094.t3 Calx Na Ca exchange protein
95.3/83
N0.HOG0015766 Aips g19069.t2 Calx Na Ca exchange protein
N0.HOG0015766 Bmor g16126.t1 Calx Na Ca exchange protein
23.5/27
0/41
N0.HOG0012386 Tsyl g5261.t1 Calx Na Ca exchange protein
99.2/100
76.6/37
N0.HOG0012386 Pmal g11566.t1 Calx Na Ca exchange protein
83.7/49
N0.HOG0012386 Mleu g13694.t1 Calx Na Ca exchange protein
Root
N0.HOG0012386 Zfil g4455.t1 Calx Na Ca exchange protein
88.5/88
N0.HOG0012386 Sbem g10118.t1 Calx Na Ca exchange protein
N0.HOG0012386 Pint g2085.t1 Calx Na Ca exchange protein
89.2/67
N0.HOG0012386 Msex g16166.t1 Calx Na Ca exchange protein
93.7/93
82.6/53
N0.HOG0012386 Bmor g16121.t1 Calx Na Ca exchange protein
80.8/63
N0.HOG0012386 Bbet g3093.t1 Calx Na Ca exchange protein
97/93
N0.HOG0012386 Hpyr g8378.t1 Calx Na Ca exchange protein
N0.HOG0012386 Aips g19071.t1 Calx Na Ca exchange protein
N0.HOG0015766 Sbem g10119.t3 Calx Na Ca exchange protein
42.8/40
N0.HOG0015766 Zfil g4456.t2 Calx Na Ca exchange protein

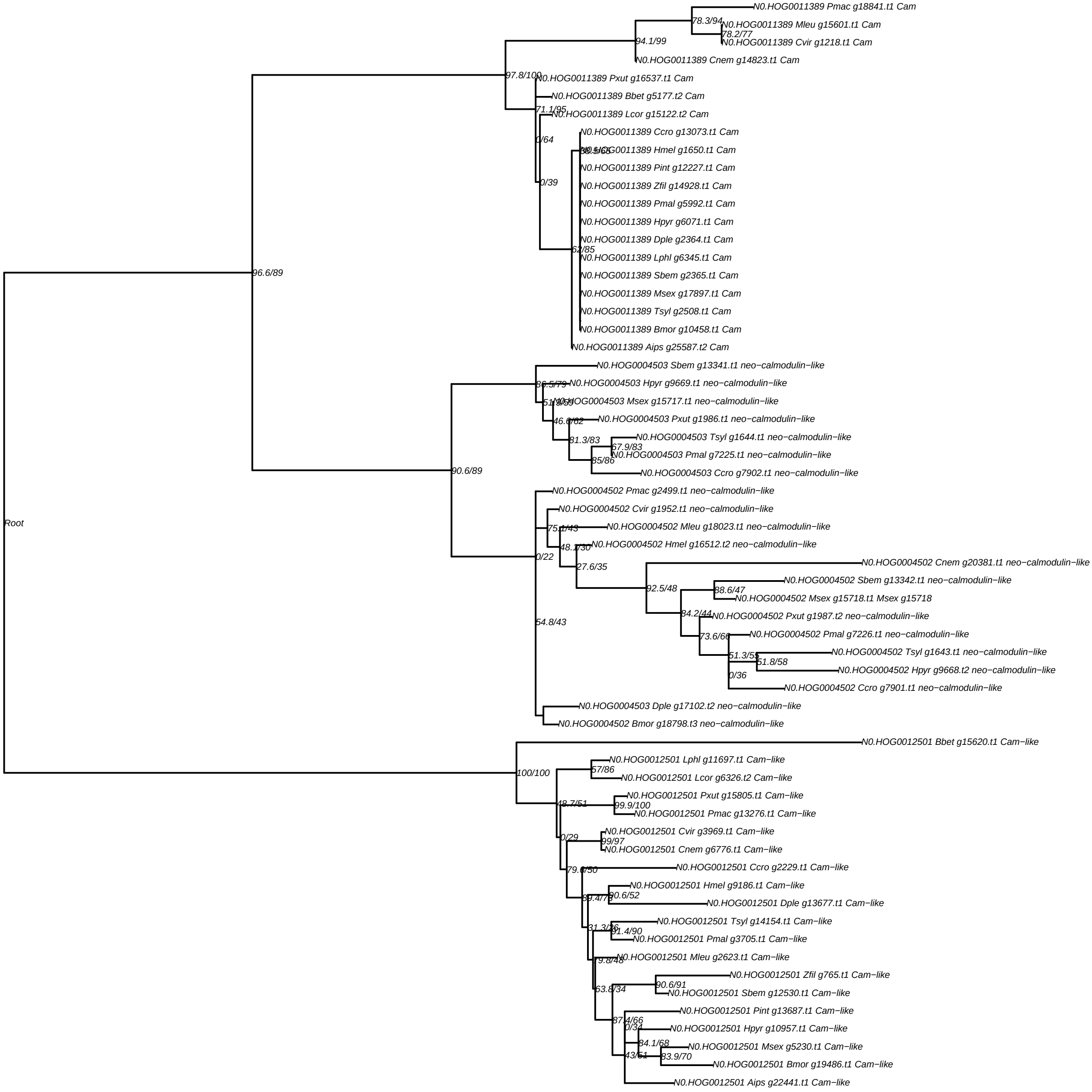

N0.HOG0011389 Pmac g18841.t1 Cam
78.3/94
N0.HOG0011389 Mleu g15601.t1 Cam
78.2/77
N0.HOG0011389 Cvir g1218.t1 Cam
94.1/99
N0.HOG0011389 Cnem g14823.t1 Cam
97.8/100
N0.HOG0011389 Pxut g16537.t1 Cam
N0.HOG0011389 Bbet g5177.t2 Cam
71.1/95
N0.HOG0011389 Lcor g15122.t2 Cam
N0.HOG0011389 Ccro g13073.t1 Cam
0/64
38.5/65
N0.HOG0011389 Hmel g1650.t1 Cam
N0.HOG0011389 Pint g12227.t1 Cam
0/39
N0.HOG0011389 Zfil g14928.t1 Cam
N0.HOG0011389 Pmal g5992.t1 Cam
N0.HOG0011389 Hpyr g6071.t1 Cam
N0.HOG0011389 Dple g2364.t1 Cam
62/85
N0.HOG0011389 Lphl g6345.t1 Cam
96.6/89
N0.HOG0011389 Sbem g2365.t1 Cam
N0.HOG0011389 Msex g17897.t1 Cam
N0.HOG0011389 Tsyl g2508.t1 Cam
N0.HOG0011389 Bmor g10458.t1 Cam
N0.HOG0011389 Aips g25587.t2 Cam
N0.HOG0004503 Sbem g13341.t1 neo−calmodulin−like
86.5/79
N0.HOG0004503 Hpyr g9669.t1 neo−calmodulin−like
51.9/59
N0.HOG0004503 Msex g15717.t1 neo−calmodulin−like
46.6/62
N0.HOG0004503 Pxut g1986.t1 neo−calmodulin−like
N0.HOG0004503 Tsyl g1644.t1 neo−calmodulin−like
81.3/83
67.9/83
N0.HOG0004503 Pmal g7225.t1 neo−calmodulin−like
85/86
90.6/89
N0.HOG0004503 Ccro g7902.t1 neo−calmodulin−like
N0.HOG0004502 Pmac g2499.t1 neo−calmodulin−like
N0.HOG0004502 Cvir g1952.t1 neo−calmodulin−like
Root
75.1/43
N0.HOG0004502 Mleu g18023.t1 neo−calmodulin−like
48.1/30
N0.HOG0004502 Hmel g16512.t2 neo−calmodulin−like
0/22
N0.HOG0004502 Cnem g20381.t1 neo−calmodulin−like
27.6/35
N0.HOG0004502 Sbem g13342.t1 neo−calmodulin−like
92.5/48
88.6/47
N0.HOG0004502 Msex g15718.t1 Msex g15718
84.2/44
N0.HOG0004502 Pxut g1987.t2 neo−calmodulin−like
54.8/43
73.6/66
N0.HOG0004502 Pmal g7226.t1 neo−calmodulin−like
N0.HOG0004502 Tsyl g1643.t1 neo−calmodulin−like
51.3/55
51.8/58
N0.HOG0004502 Hpyr g9668.t2 neo−calmodulin−like
0/36
N0.HOG0004502 Ccro g7901.t1 neo−calmodulin−like
N0.HOG0004503 Dple g17102.t2 neo−calmodulin−like
N0.HOG0004502 Bmor g18798.t3 neo−calmodulin−like
N0.HOG0012501 Bbet g15620.t1 Cam−like
N0.HOG0012501 Lphl g11697.t1 Cam−like
57/86
100/100
N0.HOG0012501 Lcor g6326.t2 Cam−like
N0.HOG0012501 Pxut g15805.t1 Cam−like
48.7/51
99.9/100
N0.HOG0012501 Pmac g13276.t1 Cam−like
N0.HOG0012501 Cvir g3969.t1 Cam−like
0/29
99/97
N0.HOG0012501 Cnem g6776.t1 Cam−like
N0.HOG0012501 Ccro g2229.t1 Cam−like
79.6/50
N0.HOG0012501 Hmel g9186.t1 Cam−like
90.6/52
89.4/78
N0.HOG0012501 Dple g13677.t1 Cam−like
N0.HOG0012501 Tsyl g14154.t1 Cam−like
31.3/26
91.4/90
N0.HOG0012501 Pmal g3705.t1 Cam−like
N0.HOG0012501 Mleu g2623.t1 Cam−like
79.8/48
N0.HOG0012501 Zfil g765.t1 Cam−like
63.8/34
90.6/91
N0.HOG0012501 Sbem g12530.t1 Cam−like
N0.HOG0012501 Pint g13687.t1 Cam−like
87.4/66
0/34
N0.HOG0012501 Hpyr g10957.t1 Cam−like
84.1/68
N0.HOG0012501 Msex g5230.t1 Cam−like
43/61
83.9/70
N0.HOG0012501 Bmor g19486.t1 Cam−like
N0.HOG0012501 Aips g22441.t1 Cam−like

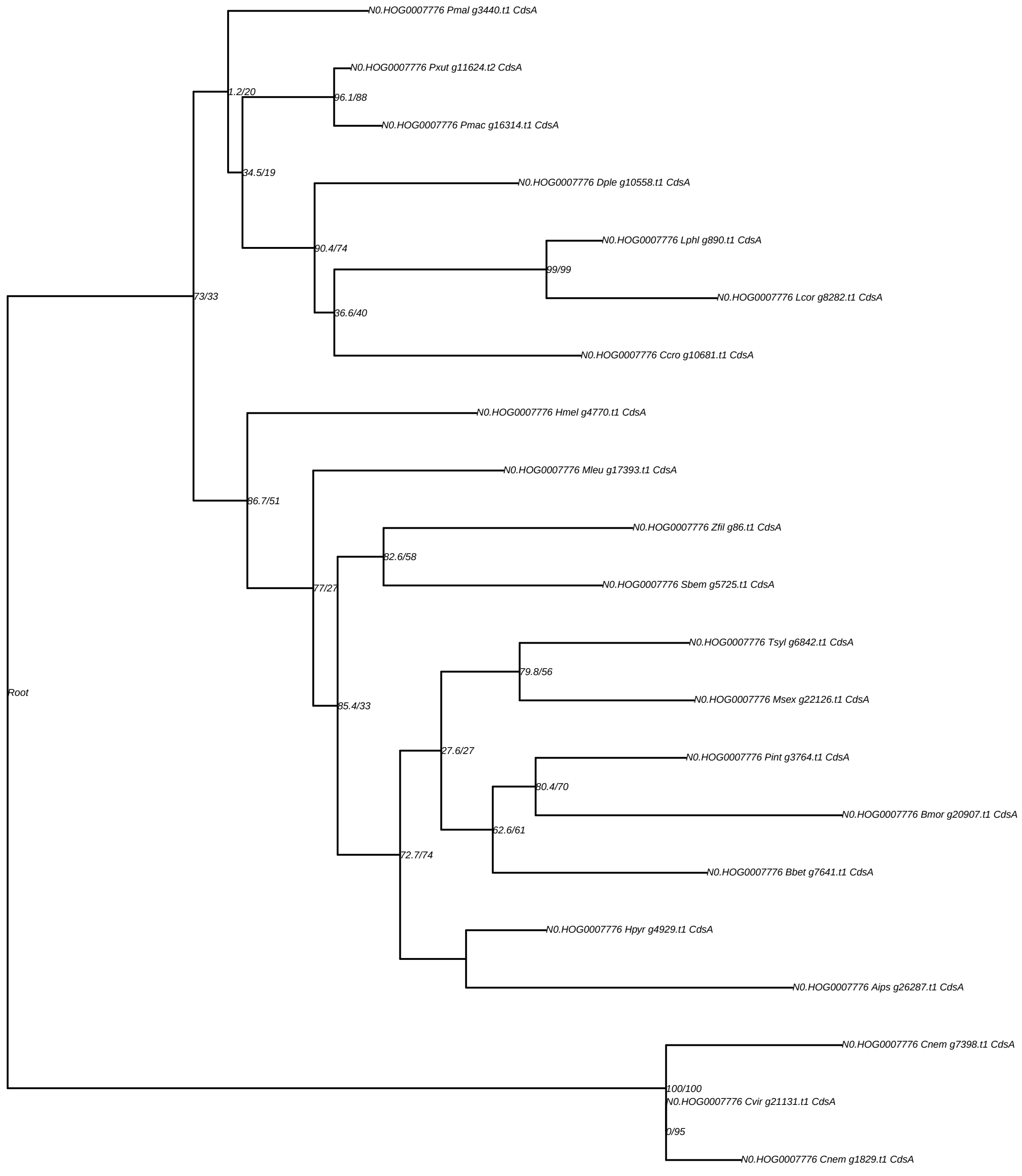
N0.HOG0007776 Pmal g3440.t1 CdsA
N0.HOG0007776 Pxut g11624.t2 CdsA
96.1/88
N0.HOG0007776 Pmac g16314.t1 CdsA
1.2/20
34.5/19
N0.HOG0007776 Dple g10558.t1 CdsA
90.4/74
N0.HOG0007776 Lphl g890.t1 CdsA
99/99
N0.HOG0007776 Lcor g8282.t1 CdsA
36.6/40
N0.HOG0007776 Ccro g10681.t1 CdsA
73/33
N0.HOG0007776 Hmel g4770.t1 CdsA
N0.HOG0007776 Mleu g17393.t1 CdsA
86.7/51
N0.HOG0007776 Zfil g86.t1 CdsA
82.6/58
N0.HOG0007776 Sbem g5725.t1 CdsA
77/27
N0.HOG0007776 Tsyl g6842.t1 CdsA
79.8/56
N0.HOG0007776 Msex g22126.t1 CdsA
85.4/33
27.6/27
N0.HOG0007776 Pint g3764.t1 CdsA
80.4/70
N0.HOG0007776 Bmor g20907.t1 CdsA
62.6/61
N0.HOG0007776 Bbet g7641.t1 CdsA
72.7/74
N0.HOG0007776 Hpyr g4929.t1 CdsA
N0.HOG0007776 Aips g26287.t1 CdsA
Root
N0.HOG0007776 Cnem g7398.t1 CdsA
100/100
N0.HOG0007776 Cvir g21131.t1 CdsA
0/95
N0.HOG0007776 Cnem g1829.t1 CdsA

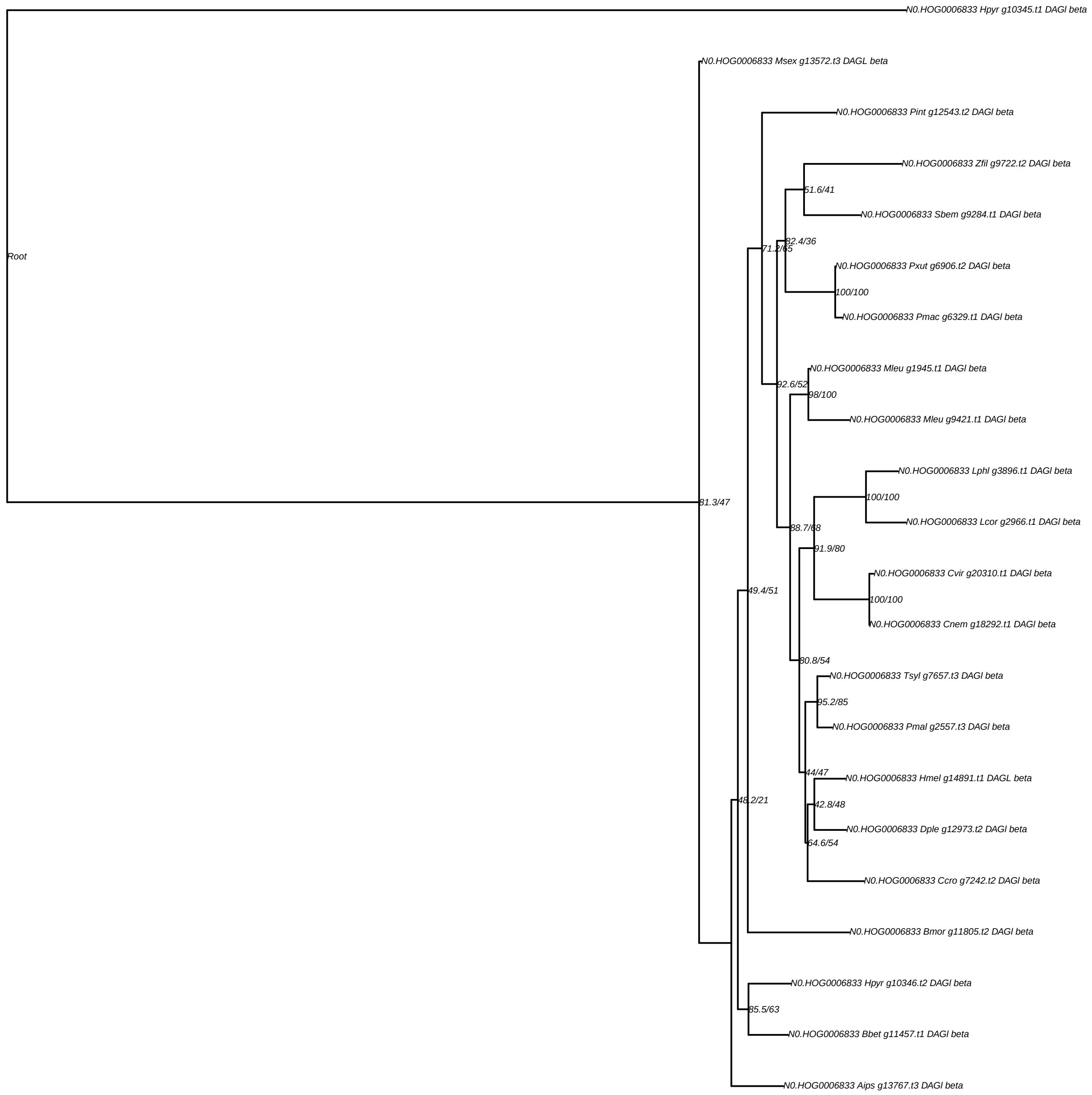

N0.HOG0006833 Hpyr g10345.t1 DAGl beta
N0.HOG0006833 Msex g13572.t3 DAGL beta
N0.HOG0006833 Pint g12543.t2 DAGl beta
N0.HOG0006833 Zfil g9722.t2 DAGl beta
51.6/41
N0.HOG0006833 Sbem g9284.t1 DAGl beta
82.4/36
Root
71.2/65
N0.HOG0006833 Pxut g6906.t2 DAGl beta
100/100
N0.HOG0006833 Pmac g6329.t1 DAGl beta
N0.HOG0006833 Mleu g1945.t1 DAGl beta
92.6/52
98/100
N0.HOG0006833 Mleu g9421.t1 DAGl beta
N0.HOG0006833 Lphl g3896.t1 DAGl beta
100/100
81.3/47
N0.HOG0006833 Lcor g2966.t1 DAGl beta
88.7/68
91.9/80
N0.HOG0006833 Cvir g20310.t1 DAGl beta
49.4/51
100/100
N0.HOG0006833 Cnem g18292.t1 DAGl beta
80.8/54
N0.HOG0006833 Tsyl g7657.t3 DAGl beta
95.2/85
N0.HOG0006833 Pmal g2557.t3 DAGl beta
44/47
N0.HOG0006833 Hmel g14891.t1 DAGL beta
48.2/21
42.8/48
N0.HOG0006833 Dple g12973.t2 DAGl beta
64.6/54
N0.HOG0006833 Ccro g7242.t2 DAGl beta
N0.HOG0006833 Bmor g11805.t2 DAGl beta
N0.HOG0006833 Hpyr g10346.t2 DAGl beta
85.5/63
N0.HOG0006833 Bbet g11457.t1 DAGl beta
N0.HOG0006833 Aips g13767.t3 DAGl beta

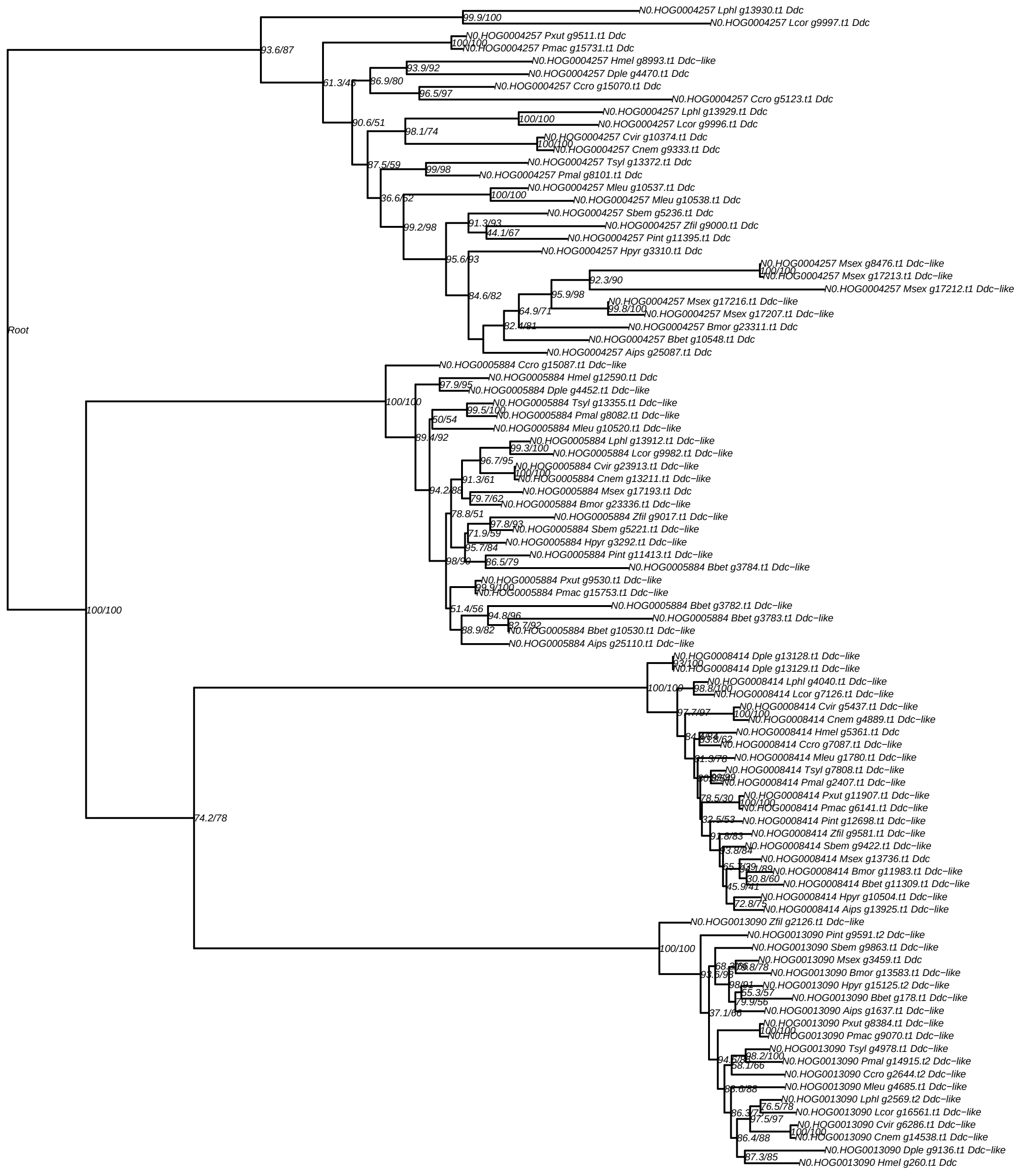

Root
N0.HOG0004257 Lphl g13930.t1 Ddc
N0.HOG0004257 Lcor g9997.t1 Ddc
N0.HOG0004257 Pxut g9511.t1 Ddc
N0.HOG0004257 Pmac g15731.t1 Ddc
N0.HOG0004257 Hmel g8993.t1 Ddc−like
N0.HOG0004257 Dple g4470.t1 Ddc
N0.HOG0004257 Ccro g15070.t1 Ddc
N0.HOG0004257 Ccro g5123.t1 Ddc
N0.HOG0004257 Lphl g13929.t1 Ddc
N0.HOG0004257 Lcor g9996.t1 Ddc
N0.HOG0004257 Cvir g10374.t1 Ddc
N0.HOG0004257 Cnem g9333.t1 Ddc
N0.HOG0004257 Tsyl g13372.t1 Ddc
N0.HOG0004257 Pmal g8101.t1 Ddc
N0.HOG0004257 Mleu g10537.t1 Ddc
N0.HOG0004257 Mleu g10538.t1 Ddc
N0.HOG0004257 Sbem g5236.t1 Ddc
N0.HOG0004257 Zfil g9000.t1 Ddc
N0.HOG0004257 Pint g11395.t1 Ddc
N0.HOG0004257 Hpyr g3310.t1 Ddc
N0.HOG0004257 Msex g8476.t1 Ddc−like
N0.HOG0004257 Msex g17213.t1 Ddc−like
N0.HOG0004257 Msex g17212.t1 Ddc−like
N0.HOG0004257 Msex g17216.t1 Ddc−like
N0.HOG0004257 Msex g17207.t1 Ddc−like
N0.HOG0004257 Bmor g23311.t1 Ddc
N0.HOG0004257 Bbet g10548.t1 Ddc
N0.HOG0004257 Aips g25087.t1 Ddc
N0.HOG0005884 Ccro g15087.t1 Ddc−like
N0.HOG0005884 Hmel g12590.t1 Ddc
N0.HOG0005884 Dple g4452.t1 Ddc−like
N0.HOG0005884 Tsyl g13355.t1 Ddc−like
N0.HOG0005884 Pmal g8082.t1 Ddc−like
N0.HOG0005884 Mleu g10520.t1 Ddc−like
N0.HOG0005884 Lphl g13912.t1 Ddc−like
N0.HOG0005884 Lcor g9982.t1 Ddc−like
N0.HOG0005884 Cvir g23913.t1 Ddc−like
N0.HOG0005884 Cnem g13211.t1 Ddc−like
N0.HOG0005884 Msex g17193.t1 Ddc
N0.HOG0005884 Bmor g23336.t1 Ddc−like
N0.HOG0005884 Zfil g9017.t1 Ddc−like
N0.HOG0005884 Sbem g5221.t1 Ddc−like
N0.HOG0005884 Hpyr g3292.t1 Ddc−like
N0.HOG0005884 Pint g11413.t1 Ddc−like
N0.HOG0005884 Bbet g3784.t1 Ddc−like
N0.HOG0005884 Pxut g9530.t1 Ddc−like
N0.HOG0005884 Pmac g15753.t1 Ddc−like
N0.HOG0005884 Bbet g3782.t1 Ddc−like
N0.HOG0005884 Bbet g3783.t1 Ddc−like
N0.HOG0005884 Bbet g10530.t1 Ddc−like
N0.HOG0005884 Aips g25110.t1 Ddc−like
N0.HOG0008414 Dple g13128.t1 Ddc−like
N0.HOG0008414 Dple g13129.t1 Ddc−like
N0.HOG0008414 Lphl g4040.t1 Ddc−like
N0.HOG0008414 Lcor g7126.t1 Ddc−like
N0.HOG0008414 Cvir g5437.t1 Ddc−like
N0.HOG0008414 Cnem g4889.t1 Ddc−like
N0.HOG0008414 Hmel g5361.t1 Ddc
N0.HOG0008414 Ccro g7087.t1 Ddc−like
N0.HOG0008414 Mleu g1780.t1 Ddc−like
N0.HOG0008414 Tsyl g7808.t1 Ddc−like
N0.HOG0008414 Pmal g2407.t1 Ddc−like
N0.HOG0008414 Pxut g11907.t1 Ddc−like
N0.HOG0008414 Pmac g6141.t1 Ddc−like
N0.HOG0008414 Pint g12698.t1 Ddc−like
N0.HOG0008414 Zfil g9581.t1 Ddc−like
N0.HOG0008414 Sbem g9422.t1 Ddc−like
N0.HOG0008414 Msex g13736.t1 Ddc
N0.HOG0008414 Bmor g11983.t1 Ddc−like
N0.HOG0008414 Bbet g11309.t1 Ddc−like
N0.HOG0008414 Hpyr g10504.t1 Ddc−like
N0.HOG0008414 Aips g13925.t1 Ddc−like
N0.HOG0013090 Zfil g2126.t1 Ddc−like
N0.HOG0013090 Pint g9591.t2 Ddc−like
N0.HOG0013090 Sbem g9863.t1 Ddc−like
N0.HOG0013090 Msex g3459.t1 Ddc
N0.HOG0013090 Bmor g13583.t1 Ddc−like
N0.HOG0013090 Hpyr g15125.t2 Ddc−like
N0.HOG0013090 Bbet g178.t1 Ddc−like
N0.HOG0013090 Aips g1637.t1 Ddc−like
N0.HOG0013090 Pxut g8384.t1 Ddc−like
N0.HOG0013090 Pmac g9070.t1 Ddc−like
N0.HOG0013090 Tsyl g4978.t1 Ddc−like
N0.HOG0013090 Pmal g14915.t2 Ddc−like
N0.HOG0013090 Ccro g2644.t2 Ddc−like
N0.HOG0013090 Mleu g4685.t1 Ddc−like
N0.HOG0013090 Lphl g2569.t2 Ddc−like
N0.HOG0013090 Lcor g16561.t1 Ddc−like
N0.HOG0013090 Cvir g6286.t1 Ddc−like
N0.HOG0013090 Cnem g14538.t1 Ddc−like
N0.HOG0013090 Dple g9136.t1 Ddc−like
N0.HOG0013090 Hmel g260.t1 Ddc

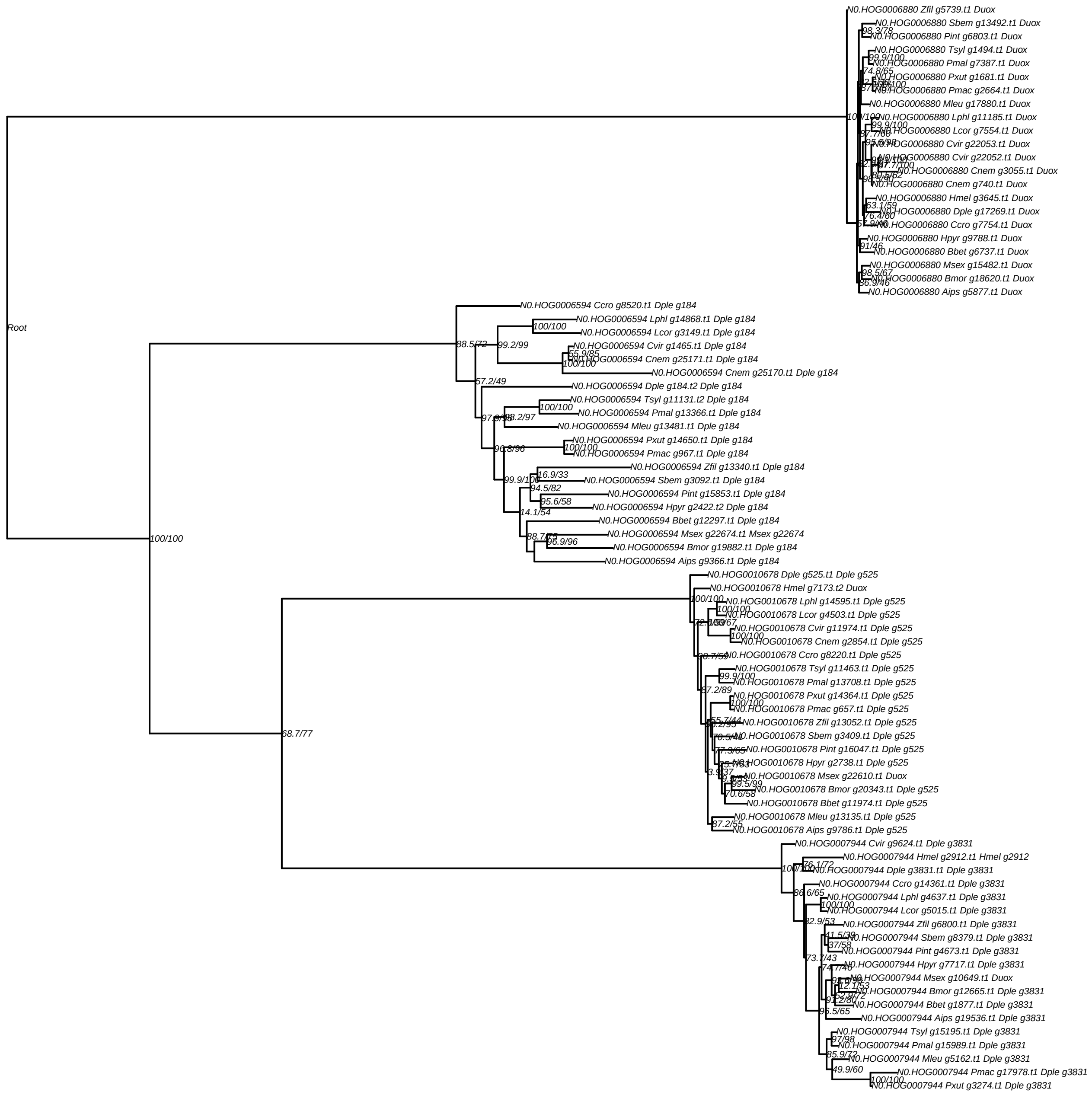
Root
N0.HOG0006880 Zfil g5739.t1 Duox
N0.HOG0006880 Sbem g13492.t1 Duox
N0.HOG0006880 Pint g6803.t1 Duox
N0.HOG0006880 Tsyl g1494.t1 Duox
N0.HOG0006880 Pmal g7387.t1 Duox
N0.HOG0006880 Pxut g1681.t1 Duox
N0.HOG0006880 Pmac g2664.t1 Duox
N0.HOG0006880 Mleu g17880.t1 Duox
N0.HOG0006880 Lphl g11185.t1 Duox
N0.HOG0006880 Lcor g7554.t1 Duox
N0.HOG0006880 Cvir g22053.t1 Duox
N0.HOG0006880 Cvir g22052.t1 Duox
N0.HOG0006880 Cnem g3055.t1 Duox
N0.HOG0006880 Cnem g740.t1 Duox
N0.HOG0006880 Hmel g3645.t1 Duox
N0.HOG0006880 Dple g17269.t1 Duox
N0.HOG0006880 Ccro g7754.t1 Duox
N0.HOG0006880 Hpyr g9788.t1 Duox
N0.HOG0006880 Bbet g6737.t1 Duox
N0.HOG0006880 Msex g15482.t1 Duox
N0.HOG0006880 Bmor g18620.t1 Duox
N0.HOG0006880 Aips g5877.t1 Duox
N0.HOG0006594 Ccro g8520.t1 Dple g184
N0.HOG0006594 Lphl g14868.t1 Dple g184
N0.HOG0006594 Lcor g3149.t1 Dple g184
N0.HOG0006594 Cvir g1465.t1 Dple g184
N0.HOG0006594 Cnem g25171.t1 Dple g184
N0.HOG0006594 Cnem g25170.t1 Dple g184
N0.HOG0006594 Dple g184.t2 Dple g184
N0.HOG0006594 Tsyl g11131.t2 Dple g184
N0.HOG0006594 Pmal g13366.t1 Dple g184
N0.HOG0006594 Mleu g13481.t1 Dple g184
N0.HOG0006594 Pxut g14650.t1 Dple g184
N0.HOG0006594 Pmac g967.t1 Dple g184
N0.HOG0006594 Zfil g13340.t1 Dple g184
N0.HOG0006594 Sbem g3092.t1 Dple g184
N0.HOG0006594 Pint g15853.t1 Dple g184
N0.HOG0006594 Hpyr g2422.t2 Dple g184
N0.HOG0006594 Bbet g12297.t1 Dple g184
N0.HOG0006594 Msex g22674.t1 Msex g22674
N0.HOG0006594 Bmor g19882.t1 Dple g184
N0.HOG0006594 Aips g9366.t1 Dple g184
N0.HOG0010678 Dple g525.t1 Dple g525
N0.HOG0010678 Hmel g7173.t2 Duox
N0.HOG0010678 Lphl g14595.t1 Dple g525
N0.HOG0010678 Lcor g4503.t1 Dple g525
N0.HOG0010678 Cvir g11974.t1 Dple g525
N0.HOG0010678 Cnem g2854.t1 Dple g525
N0.HOG0010678 Ccro g8220.t1 Dple g525
N0.HOG0010678 Tsyl g11463.t1 Dple g525
N0.HOG0010678 Pmal g13708.t1 Dple g525
N0.HOG0010678 Pxut g14364.t1 Dple g525
N0.HOG0010678 Pmac g657.t1 Dple g525
N0.HOG0010678 Zfil g13052.t1 Dple g525
N0.HOG0010678 Sbem g3409.t1 Dple g525
N0.HOG0010678 Pint g16047.t1 Dple g525
N0.HOG0010678 Hpyr g2738.t1 Dple g525
N0.HOG0010678 Msex g22610.t1 Duox
N0.HOG0010678 Bmor g20343.t1 Dple g525
N0.HOG0010678 Bbet g11974.t1 Dple g525
N0.HOG0010678 Mleu g13135.t1 Dple g525
N0.HOG0010678 Aips g9786.t1 Dple g525
N0.HOG0007944 Cvir g9624.t1 Dple g3831
N0.HOG0007944 Hmel g2912.t1 Hmel g2912
N0.HOG0007944 Dple g3831.t1 Dple g3831
N0.HOG0007944 Ccro g14361.t1 Dple g3831
N0.HOG0007944 Lphl g4637.t1 Dple g3831
N0.HOG0007944 Lcor g5015.t1 Dple g3831
N0.HOG0007944 Zfil g6800.t1 Dple g3831
N0.HOG0007944 Sbem g8379.t1 Dple g3831
N0.HOG0007944 Pint g4673.t1 Dple g3831
N0.HOG0007944 Hpyr g7717.t1 Dple g3831
N0.HOG0007944 Msex g10649.t1 Duox
N0.HOG0007944 Bmor g12665.t1 Dple g3831
N0.HOG0007944 Bbet g1877.t1 Dple g3831
N0.HOG0007944 Aips g19536.t1 Dple g3831
N0.HOG0007944 Tsyl g15195.t1 Dple g3831
N0.HOG0007944 Pmal g15989.t1 Dple g3831
N0.HOG0007944 Mleu g5162.t1 Dple g3831
N0.HOG0007944 Pmac g17978.t1 Dple g3831
N0.HOG0007944 Pxut g3274.t1 Dple g3831

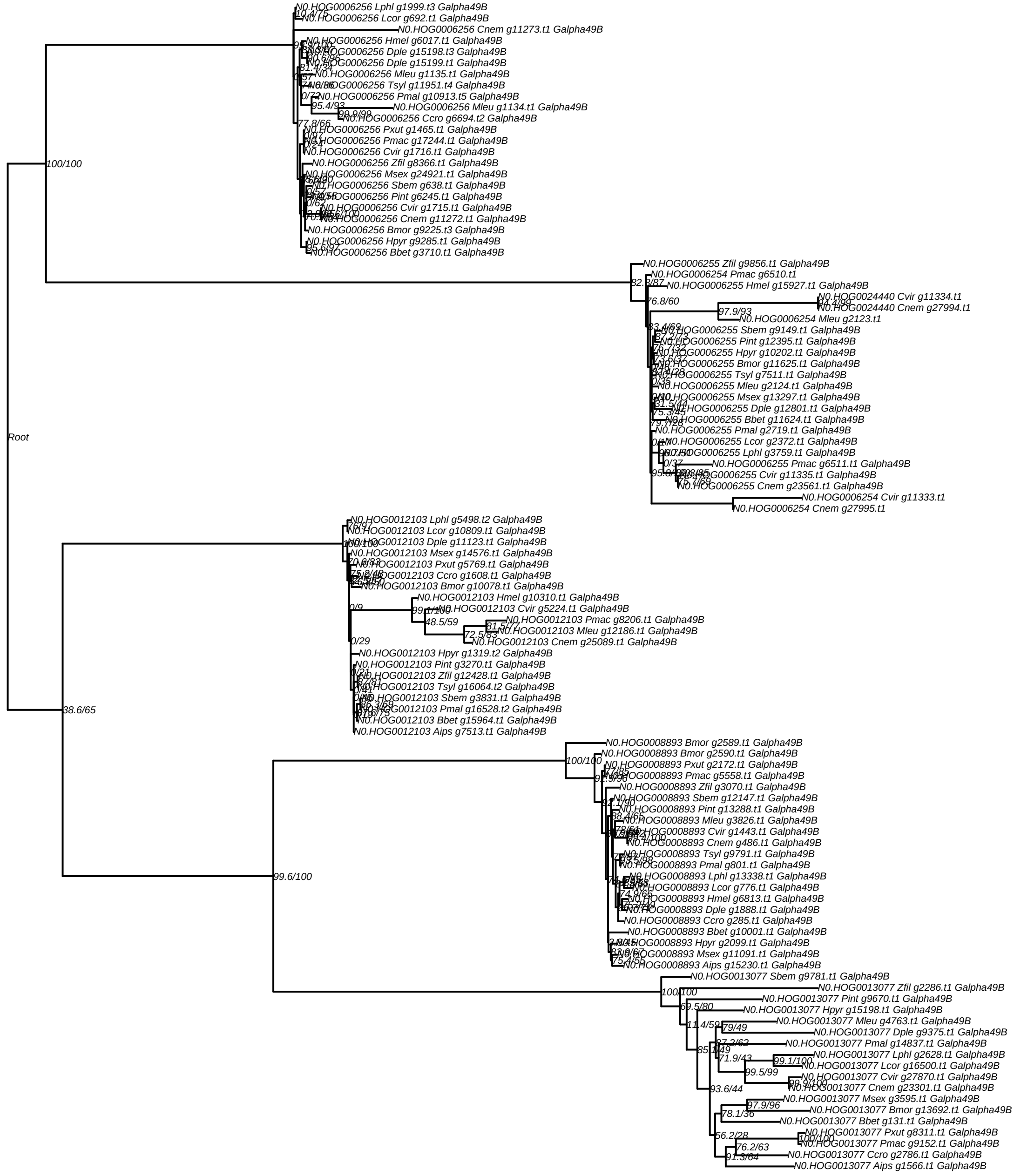

N0.HOG0006256 Lphl g1999.t3 Galpha49B
N0.HOG0006256 Lcor g692.t1 Galpha49B
N0.HOG0006256 Cnem g11273.t1 Galpha49B
N0.HOG0006256 Hmel g6017.t1 Galpha49B
N0.HOG0006256 Dple g15198.t3 Galpha49B
N0.HOG0006256 Dple g15199.t1 Galpha49B
N0.HOG0006256 Mleu g1135.t1 Galpha49B
N0.HOG0006256 Tsyl g11951.t4 Galpha49B
N0.HOG0006256 Pmal g10913.t5 Galpha49B
N0.HOG0006256 Mleu g1134.t1 Galpha49B
N0.HOG0006256 Ccro g6694.t2 Galpha49B
N0.HOG0006256 Pxut g1465.t1 Galpha49B
N0.HOG0006256 Pmac g17244.t1 Galpha49B
N0.HOG0006256 Cvir g1716.t1 Galpha49B
N0.HOG0006256 Zfil g8366.t1 Galpha49B
N0.HOG0006256 Msex g24921.t1 Galpha49B
N0.HOG0006256 Sbem g638.t1 Galpha49B
N0.HOG0006256 Pint g6245.t1 Galpha49B
N0.HOG0006256 Cvir g1715.t1 Galpha49B
N0.HOG0006256 Cnem g11272.t1 Galpha49B
N0.HOG0006256 Bmor g9225.t3 Galpha49B
N0.HOG0006256 Hpyr g9285.t1 Galpha49B
N0.HOG0006256 Bbet g3710.t1 Galpha49B
100/100
Root
N0.HOG0006255 Zfil g9856.t1 Galpha49B
N0.HOG0006254 Pmac g6510.t1
N0.HOG0006255 Hmel g15927.t1 Galpha49B
N0.HOG0024440 Cvir g11334.t1
N0.HOG0024440 Cnem g27994.t1
N0.HOG0006254 Mleu g2123.t1
N0.HOG0006255 Sbem g9149.t1 Galpha49B
N0.HOG0006255 Pint g12395.t1 Galpha49B
N0.HOG0006255 Hpyr g10202.t1 Galpha49B
N0.HOG0006255 Bmor g11625.t1 Galpha49B
N0.HOG0006255 Tsyl g7511.t1 Galpha49B
N0.HOG0006255 Mleu g2124.t1 Galpha49B
N0.HOG0006255 Msex g13297.t1 Galpha49B
N0.HOG0006255 Dple g12801.t1 Galpha49B
N0.HOG0006255 Bbet g11624.t1 Galpha49B
N0.HOG0006255 Pmal g2719.t1 Galpha49B
N0.HOG0006255 Lcor g2372.t1 Galpha49B
N0.HOG0006255 Lphl g3759.t1 Galpha49B
N0.HOG0006255 Pmac g6511.t1 Galpha49B
N0.HOG0006255 Cvir g11335.t1 Galpha49B
N0.HOG0006255 Cnem g23561.t1 Galpha49B
N0.HOG0006254 Cvir g11333.t1
N0.HOG0006254 Cnem g27995.t1
N0.HOG0012103 Lphl g5498.t2 Galpha49B
N0.HOG0012103 Lcor g10809.t1 Galpha49B
N0.HOG0012103 Dple g11123.t1 Galpha49B
N0.HOG0012103 Msex g14576.t1 Galpha49B
N0.HOG0012103 Pxut g5769.t1 Galpha49B
N0.HOG0012103 Ccro g1608.t1 Galpha49B
N0.HOG0012103 Bmor g10078.t1 Galpha49B
N0.HOG0012103 Hmel g10310.t1 Galpha49B
N0.HOG0012103 Cvir g5224.t1 Galpha49B
N0.HOG0012103 Pmac g8206.t1 Galpha49B
N0.HOG0012103 Mleu g12186.t1 Galpha49B
N0.HOG0012103 Cnem g25089.t1 Galpha49B
N0.HOG0012103 Hpyr g1319.t2 Galpha49B
N0.HOG0012103 Pint g3270.t1 Galpha49B
N0.HOG0012103 Zfil g12428.t1 Galpha49B
N0.HOG0012103 Tsyl g16064.t2 Galpha49B
N0.HOG0012103 Sbem g3831.t1 Galpha49B
N0.HOG0012103 Pmal g16528.t2 Galpha49B
N0.HOG0012103 Bbet g15964.t1 Galpha49B
N0.HOG0012103 Aips g7513.t1 Galpha49B
38.6/65
N0.HOG0008893 Bmor g2589.t1 Galpha49B
N0.HOG0008893 Bmor g2590.t1 Galpha49B
N0.HOG0008893 Pxut g2172.t1 Galpha49B
N0.HOG0008893 Pmac g5558.t1 Galpha49B
N0.HOG0008893 Zfil g3070.t1 Galpha49B
N0.HOG0008893 Sbem g12147.t1 Galpha49B
N0.HOG0008893 Pint g13288.t1 Galpha49B
N0.HOG0008893 Mleu g3826.t1 Galpha49B
N0.HOG0008893 Cvir g1443.t1 Galpha49B
N0.HOG0008893 Cnem g486.t1 Galpha49B
N0.HOG0008893 Tsyl g9791.t1 Galpha49B
N0.HOG0008893 Pmal g801.t1 Galpha49B
N0.HOG0008893 Lphl g13338.t1 Galpha49B
N0.HOG0008893 Lcor g776.t1 Galpha49B
N0.HOG0008893 Hmel g6813.t1 Galpha49B
N0.HOG0008893 Dple g1888.t1 Galpha49B
N0.HOG0008893 Ccro g285.t1 Galpha49B
N0.HOG0008893 Bbet g10001.t1 Galpha49B
N0.HOG0008893 Hpyr g2099.t1 Galpha49B
N0.HOG0008893 Msex g11091.t1 Galpha49B
N0.HOG0008893 Aips g15230.t1 Galpha49B
99.6/100
N0.HOG0013077 Sbem g9781.t1 Galpha49B
N0.HOG0013077 Zfil g2286.t1 Galpha49B
N0.HOG0013077 Pint g9670.t1 Galpha49B
N0.HOG0013077 Hpyr g15198.t1 Galpha49B
N0.HOG0013077 Mleu g4763.t1 Galpha49B
N0.HOG0013077 Dple g9375.t1 Galpha49B
N0.HOG0013077 Pmal g14837.t1 Galpha49B
N0.HOG0013077 Lphl g2628.t1 Galpha49B
N0.HOG0013077 Lcor g16500.t1 Galpha49B
N0.HOG0013077 Cvir g27870.t1 Galpha49B
N0.HOG0013077 Cnem g23301.t1 Galpha49B
N0.HOG0013077 Msex g3595.t1 Galpha49B
N0.HOG0013077 Bmor g13692.t1 Galpha49B
N0.HOG0013077 Bbet g131.t1 Galpha49B
N0.HOG0013077 Pxut g8311.t1 Galpha49B
N0.HOG0013077 Pmac g9152.t1 Galpha49B
N0.HOG0013077 Ccro g2786.t1 Galpha49B
N0.HOG0013077 Aips g1566.t1 Galpha49B

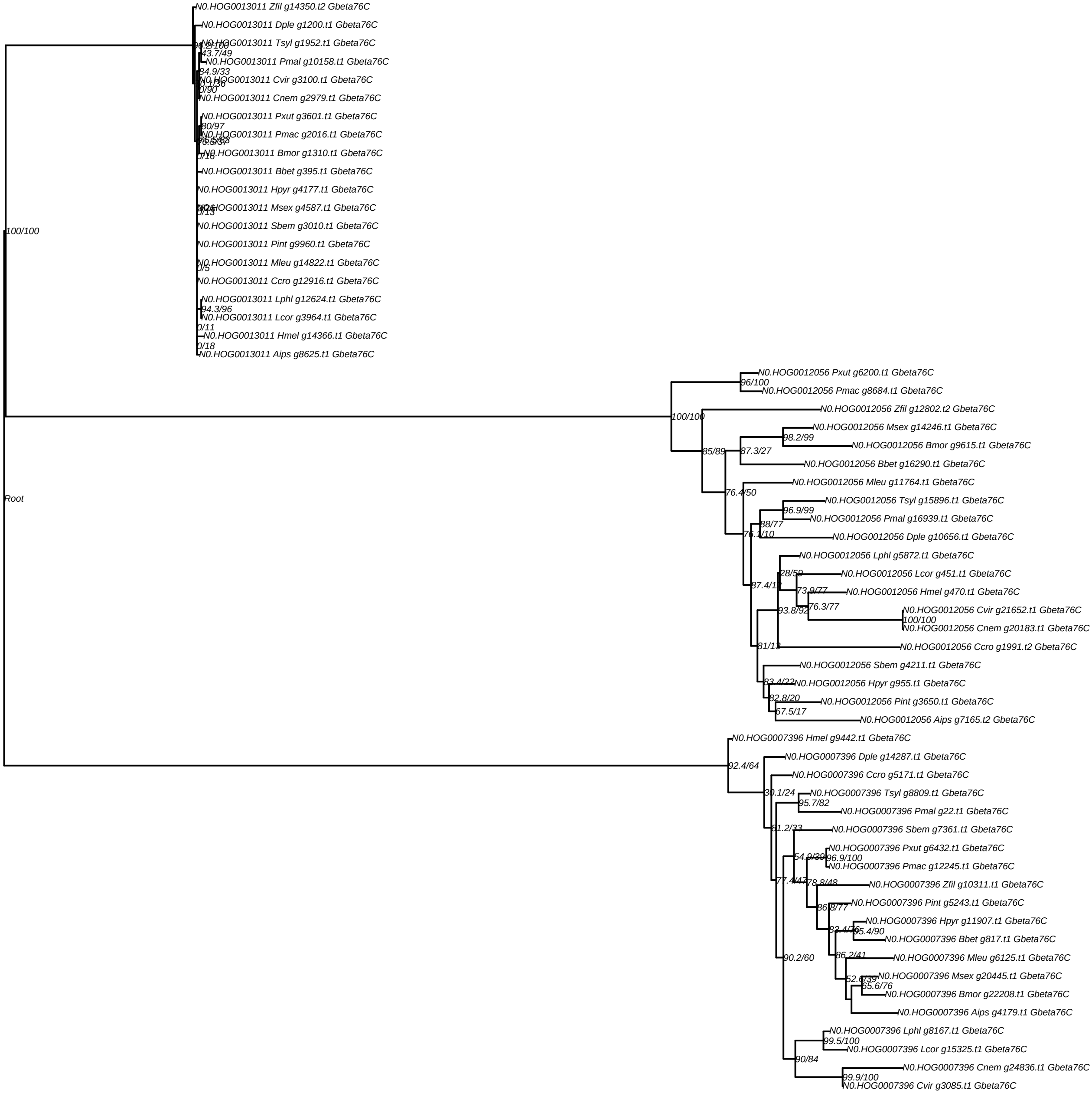
N0.HOG0013011 Zfil g14350.t2 Gbeta76C
N0.HOG0013011 Dple g1200.t1 Gbeta76C
N0.HOG0013011 Tsyl g1952.t1 Gbeta76C
43.7/49
N0.HOG0013011 Pmal g10158.t1 Gbeta76C
84.9/33
N0.HOG0013011 Cvir g3100.t1 Gbeta76C
0/90
N0.HOG0013011 Cnem g2979.t1 Gbeta76C
N0.HOG0013011 Pxut g3601.t1 Gbeta76C
80/97
N0.HOG0013011 Pmac g2016.t1 Gbeta76C
N0.HOG0013011 Bmor g1310.t1 Gbeta76C
0/16
N0.HOG0013011 Bbet g395.t1 Gbeta76C
N0.HOG0013011 Hpyr g4177.t1 Gbeta76C
N0.HOG0013011 Msex g4587.t1 Gbeta76C
0/13
N0.HOG0013011 Sbem g3010.t1 Gbeta76C
N0.HOG0013011 Pint g9960.t1 Gbeta76C
N0.HOG0013011 Mleu g14822.t1 Gbeta76C
0/5
N0.HOG0013011 Ccro g12916.t1 Gbeta76C
N0.HOG0013011 Lphl g12624.t1 Gbeta76C
94.3/96
N0.HOG0013011 Lcor g3964.t1 Gbeta76C
0/11
N0.HOG0013011 Hmel g14366.t1 Gbeta76C
0/18
N0.HOG0013011 Aips g8625.t1 Gbeta76C
100/100
Root
N0.HOG0012056 Pxut g6200.t1 Gbeta76C
96/100
N0.HOG0012056 Pmac g8684.t1 Gbeta76C
N0.HOG0012056 Zfil g12802.t2 Gbeta76C
100/100
N0.HOG0012056 Msex g14246.t1 Gbeta76C
98.2/99
N0.HOG0012056 Bmor g9615.t1 Gbeta76C
85/89
87.3/27
N0.HOG0012056 Bbet g16290.t1 Gbeta76C
N0.HOG0012056 Mleu g11764.t1 Gbeta76C
76.4/50
N0.HOG0012056 Tsyl g15896.t1 Gbeta76C
96.9/99
N0.HOG0012056 Pmal g16939.t1 Gbeta76C
88/77
N0.HOG0012056 Dple g10656.t1 Gbeta76C
N0.HOG0012056 Lphl g5872.t1 Gbeta76C
28/59
N0.HOG0012056 Lcor g451.t1 Gbeta76C
N0.HOG0012056 Hmel g470.t1 Gbeta76C
73.9/77
76.3/77
N0.HOG0012056 Cvir g21652.t1 Gbeta76C
93.8/92
100/100
N0.HOG0012056 Cnem g20183.t1 Gbeta76C
N0.HOG0012056 Ccro g1991.t2 Gbeta76C
N0.HOG0012056 Sbem g4211.t1 Gbeta76C
N0.HOG0012056 Hpyr g955.t1 Gbeta76C
82.8/20
N0.HOG0012056 Pint g3650.t1 Gbeta76C
67.5/17
N0.HOG0012056 Aips g7165.t2 Gbeta76C
N0.HOG0007396 Hmel g9442.t1 Gbeta76C
N0.HOG0007396 Dple g14287.t1 Gbeta76C
92.4/64
N0.HOG0007396 Ccro g5171.t1 Gbeta76C
N0.HOG0007396 Tsyl g8809.t1 Gbeta76C
95.7/82
N0.HOG0007396 Pmal g22.t1 Gbeta76C
81.2/33
N0.HOG0007396 Sbem g7361.t1 Gbeta76C
N0.HOG0007396 Pxut g6432.t1 Gbeta76C
96.9/100
N0.HOG0007396 Pmac g12245.t1 Gbeta76C
78.8/48
N0.HOG0007396 Zfil g10311.t1 Gbeta76C
N0.HOG0007396 Pint g5243.t1 Gbeta76C
N0.HOG0007396 Hpyr g11907.t1 Gbeta76C
N0.HOG0007396 Bbet g817.t1 Gbeta76C
86.2/41
N0.HOG0007396 Mleu g6125.t1 Gbeta76C
90.2/60
N0.HOG0007396 Msex g20445.t1 Gbeta76C
65.6/76
N0.HOG0007396 Bmor g22208.t1 Gbeta76C
N0.HOG0007396 Aips g4179.t1 Gbeta76C
N0.HOG0007396 Lphl g8167.t1 Gbeta76C
99.5/100
N0.HOG0007396 Lcor g15325.t1 Gbeta76C
90/84
N0.HOG0007396 Cnem g24836.t1 Gbeta76C
99.9/100
N0.HOG0007396 Cvir g3085.t1 Gbeta76C

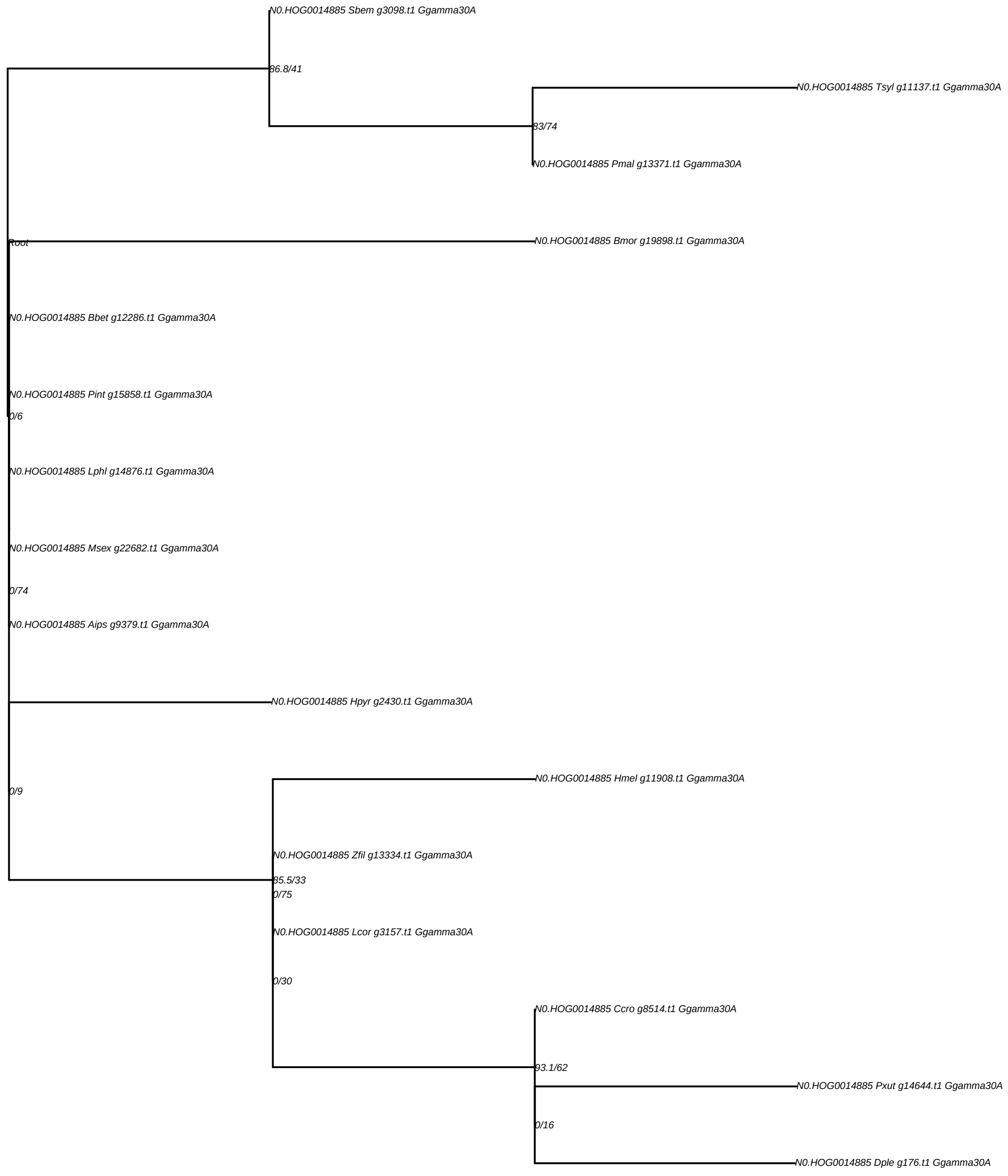
N0.HOG0014885 Sbem g3098.t1 Ggamma30A
86.8/41
N0.HOG0014885 Tsyl g11137.t1 Ggamma30A
83/74
N0.HOG0014885 Pmal g13371.t1 Ggamma30A
Root
N0.HOG0014885 Bmor g19898.t1 Ggamma30A
N0.HOG0014885 Bbet g12286.t1 Ggamma30A
N0.HOG0014885 Pint g15858.t1 Ggamma30A
0/6
N0.HOG0014885 Lphl g14876.t1 Ggamma30A
N0.HOG0014885 Msex g22682.t1 Ggamma30A
0/74
N0.HOG0014885 Aips g9379.t1 Ggamma30A
N0.HOG0014885 Hpyr g2430.t1 Ggamma30A
N0.HOG0014885 Hmel g11908.t1 Ggamma30A
0/9
N0.HOG0014885 Zfil g13334.t1 Ggamma30A
85.5/33
0/75
N0.HOG0014885 Lcor g3157.t1 Ggamma30A
0/30
N0.HOG0014885 Ccro g8514.t1 Ggamma30A
93.1/62
N0.HOG0014885 Pxut g14644.t1 Ggamma30A
0/16
N0.HOG0014885 Dple g176.t1 Ggamma30A

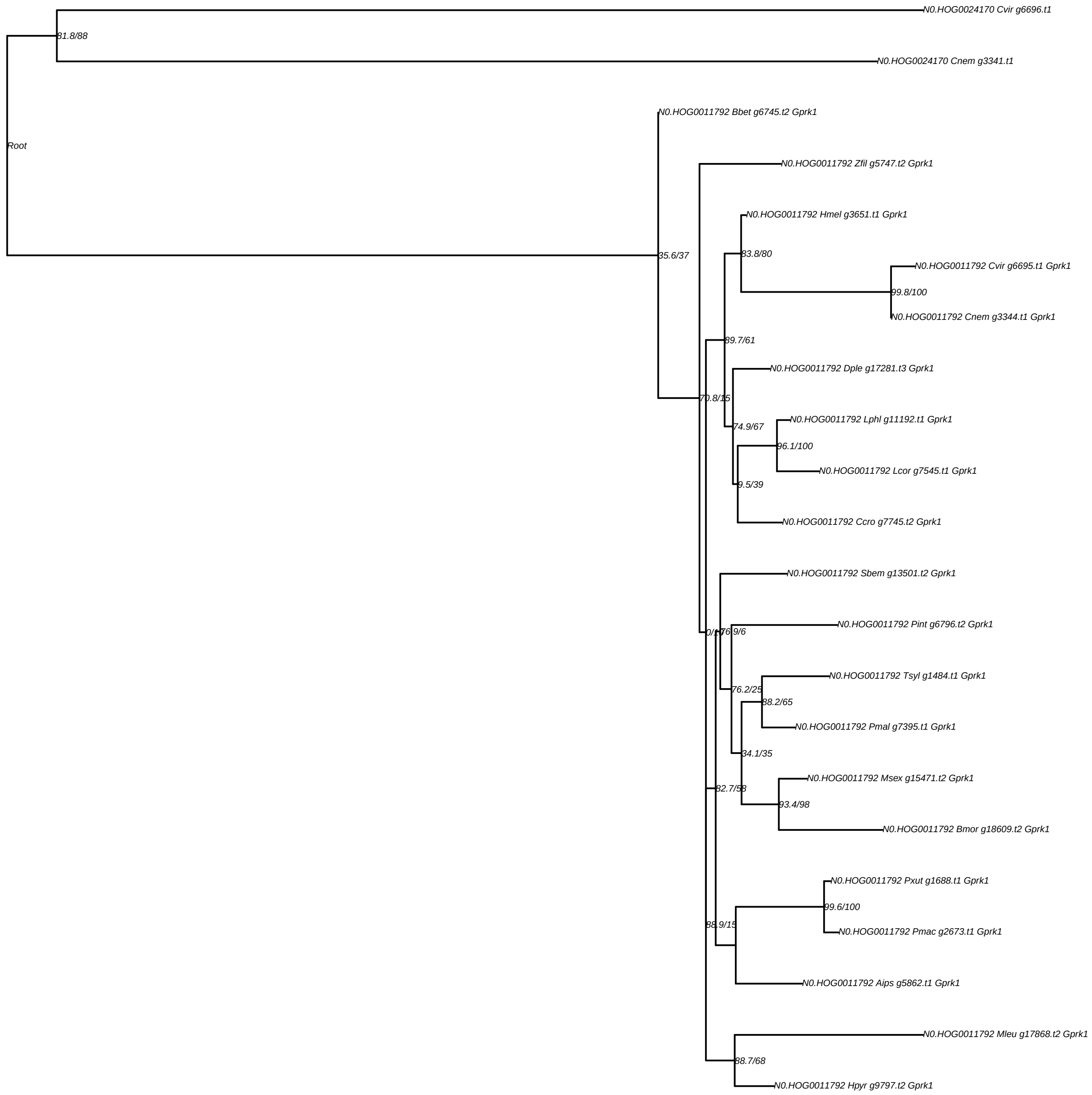

N0.HOG0024170 Cvir g6696.t1
81.8/88
N0.HOG0024170 Cnem g3341.t1
N0.HOG0011792 Bbet g6745.t2 Gprk1
Root
N0.HOG0011792 Zfil g5747.t2 Gprk1
N0.HOG0011792 Hmel g3651.t1 Gprk1
83.8/80
35.6/37
N0.HOG0011792 Cvir g6695.t1 Gprk1
99.8/100
N0.HOG0011792 Cnem g3344.t1 Gprk1
89.7/61
N0.HOG0011792 Dple g17281.t3 Gprk1
70.8/15
74.9/67
N0.HOG0011792 Lphl g11192.t1 Gprk1
96.1/100
N0.HOG0011792 Lcor g7545.t1 Gprk1
9.5/39
N0.HOG0011792 Ccro g7745.t2 Gprk1
N0.HOG0011792 Sbem g13501.t2 Gprk1
N0.HOG0011792 Pint g6796.t2 Gprk1
0/17
76.9/6
N0.HOG0011792 Tsyl g1484.t1 Gprk1
76.2/25
88.2/65
N0.HOG0011792 Pmal g7395.t1 Gprk1
34.1/35
N0.HOG0011792 Msex g15471.t2 Gprk1
82.7/58
93.4/98
N0.HOG0011792 Bmor g18609.t2 Gprk1
N0.HOG0011792 Pxut g1688.t1 Gprk1
99.6/100
88.9/15
N0.HOG0011792 Pmac g2673.t1 Gprk1
N0.HOG0011792 Aips g5862.t1 Gprk1
N0.HOG0011792 Mleu g17868.t2 Gprk1
88.7/68
N0.HOG0011792 Hpyr g9797.t2 Gprk1

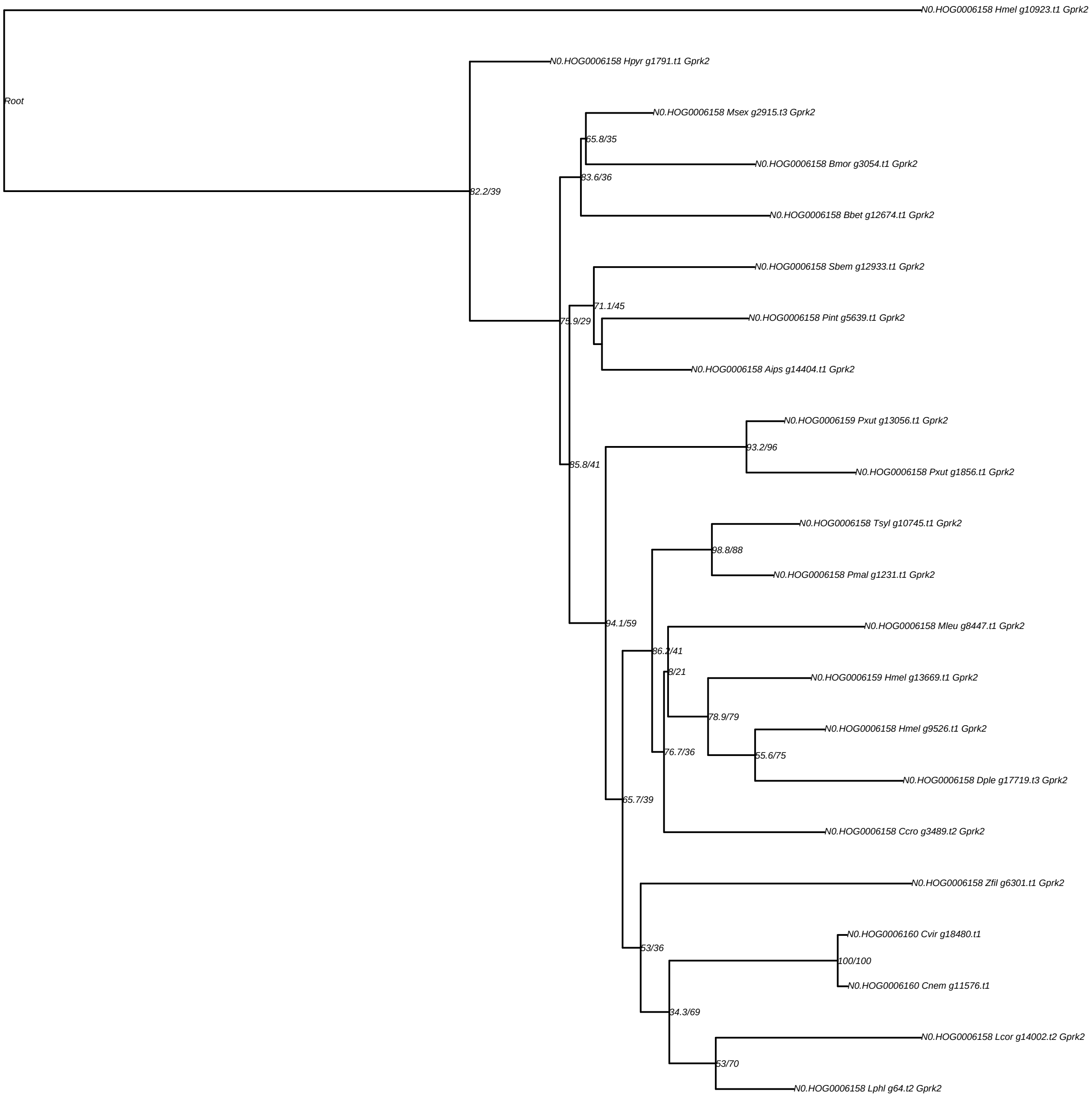

Root
N0.HOG0006158 Hmel g10923.t1 Gprk2
N0.HOG0006158 Hpyr g1791.t1 Gprk2
82.2/39
N0.HOG0006158 Msex g2915.t3 Gprk2
65.8/35
N0.HOG0006158 Bmor g3054.t1 Gprk2
83.6/36
N0.HOG0006158 Bbet g12674.t1 Gprk2
N0.HOG0006158 Sbem g12933.t1 Gprk2
71.1/45
N0.HOG0006158 Pint g5639.t1 Gprk2
75.9/29
N0.HOG0006158 Aips g14404.t1 Gprk2
N0.HOG0006159 Pxut g13056.t1 Gprk2
93.2/96
85.8/41
N0.HOG0006158 Pxut g1856.t1 Gprk2
N0.HOG0006158 Tsyl g10745.t1 Gprk2
98.8/88
N0.HOG0006158 Pmal g1231.t1 Gprk2
94.1/59
N0.HOG0006158 Mleu g8447.t1 Gprk2
86.2/41
8/21
N0.HOG0006159 Hmel g13669.t1 Gprk2
78.9/79
N0.HOG0006158 Hmel g9526.t1 Gprk2
76.7/36
55.6/75
N0.HOG0006158 Dple g17719.t3 Gprk2
65.7/39
N0.HOG0006158 Ccro g3489.t2 Gprk2
N0.HOG0006158 Zfil g6301.t1 Gprk2
N0.HOG0006160 Cvir g18480.t1
53/36
100/100
N0.HOG0006160 Cnem g11576.t1
34.3/69
N0.HOG0006158 Lcor g14002.t2 Gprk2
53/70
N0.HOG0006158 Lphl g64.t2 Gprk2

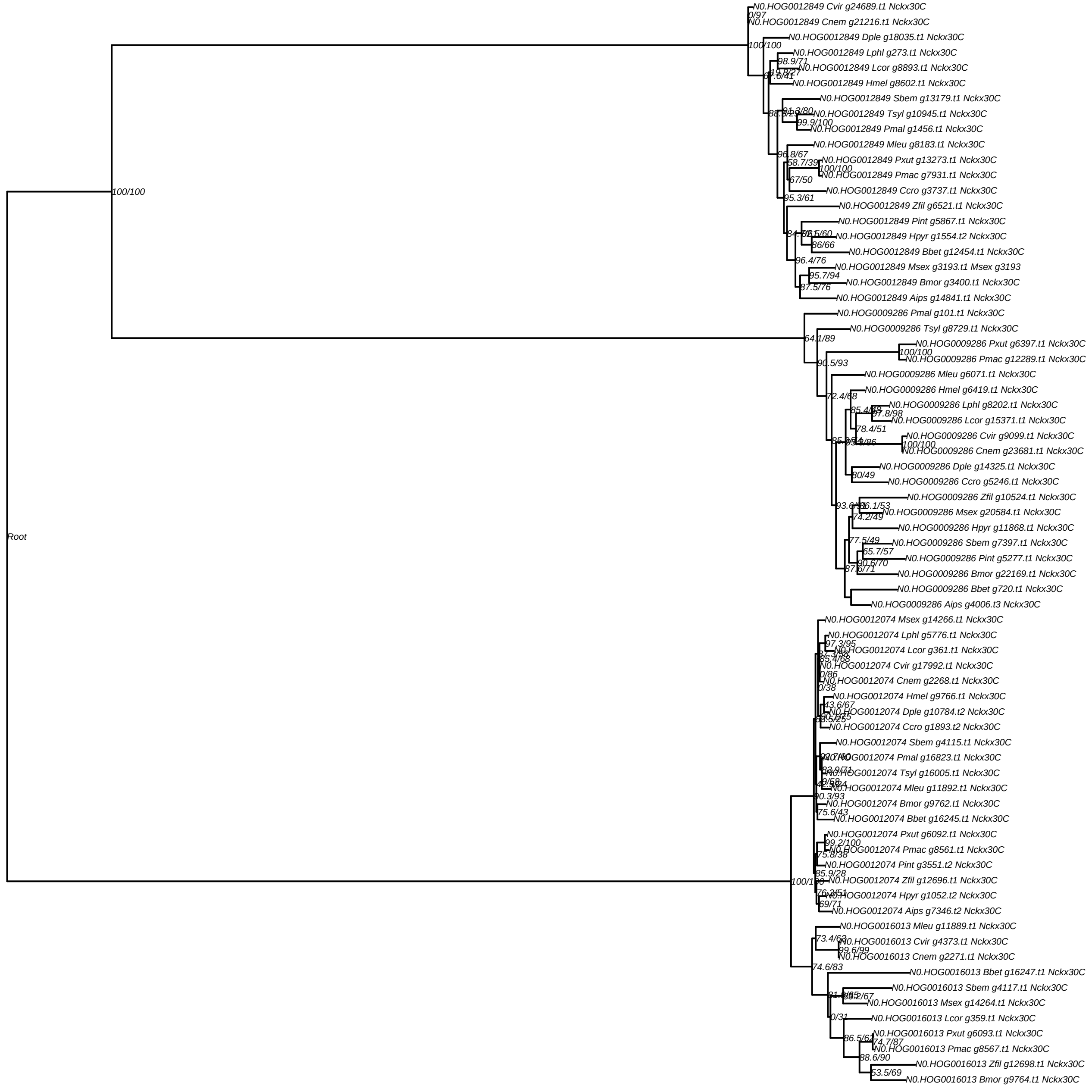
Root
100/100
N0.HOG0012849 Cvir g24689.t1 Nckx30C
0/97
N0.HOG0012849 Cnem g21216.t1 Nckx30C
100/100
N0.HOG0012849 Dple g18035.t1 Nckx30C
N0.HOG0012849 Lphl g273.t1 Nckx30C
98.9/71
N0.HOG0012849 Lcor g8893.t1 Nckx30C
N0.HOG0012849 Hmel g8602.t1 Nckx30C
N0.HOG0012849 Sbem g13179.t1 Nckx30C
N0.HOG0012849 Tsyl g10945.t1 Nckx30C
99.9/100
N0.HOG0012849 Pmal g1456.t1 Nckx30C
N0.HOG0012849 Mleu g8183.t1 Nckx30C
96.8/67
58.7/39
N0.HOG0012849 Pxut g13273.t1 Nckx30C
100/100
N0.HOG0012849 Pmac g7931.t1 Nckx30C
67/50
N0.HOG0012849 Ccro g3737.t1 Nckx30C
95.3/61
N0.HOG0012849 Zfil g6521.t1 Nckx30C
N0.HOG0012849 Pint g5867.t1 Nckx30C
N0.HOG0012849 Hpyr g1554.t2 Nckx30C
86/66
N0.HOG0012849 Bbet g12454.t1 Nckx30C
96.4/76
N0.HOG0012849 Msex g3193.t1 Msex g3193
95.7/94
N0.HOG0012849 Bmor g3400.t1 Nckx30C
87.5/76
N0.HOG0012849 Aips g14841.t1 Nckx30C
N0.HOG0009286 Pmal g101.t1 Nckx30C
64.1/89
N0.HOG0009286 Tsyl g8729.t1 Nckx30C
N0.HOG0009286 Pxut g6397.t1 Nckx30C
100/100
N0.HOG0009286 Pmac g12289.t1 Nckx30C
90.5/93
N0.HOG0009286 Mleu g6071.t1 Nckx30C
N0.HOG0009286 Hmel g6419.t1 Nckx30C
72.4/68
N0.HOG0009286 Lphl g8202.t1 Nckx30C
97.8/98
N0.HOG0009286 Lcor g15371.t1 Nckx30C
78.4/51
N0.HOG0009286 Cvir g9099.t1 Nckx30C
100/100
N0.HOG0009286 Cnem g23681.t1 Nckx30C
N0.HOG0009286 Dple g14325.t1 Nckx30C
80/49
N0.HOG0009286 Ccro g5246.t1 Nckx30C
N0.HOG0009286 Zfil g10524.t1 Nckx30C
N0.HOG0009286 Msex g20584.t1 Nckx30C
74.2/49
N0.HOG0009286 Hpyr g11868.t1 Nckx30C
77.5/49
N0.HOG0009286 Sbem g7397.t1 Nckx30C
65.7/57
N0.HOG0009286 Pint g5277.t1 Nckx30C
90.6/70
N0.HOG0009286 Bmor g22169.t1 Nckx30C
N0.HOG0009286 Bbet g720.t1 Nckx30C
N0.HOG0009286 Aips g4006.t3 Nckx30C
N0.HOG0012074 Msex g14266.t1 Nckx30C
N0.HOG0012074 Lphl g5776.t1 Nckx30C
97.3/95
N0.HOG0012074 Lcor g361.t1 Nckx30C
85.4/68
N0.HOG0012074 Cvir g17992.t1 Nckx30C
0/86
N0.HOG0012074 Cnem g2268.t1 Nckx30C
0/38
N0.HOG0012074 Hmel g9766.t1 Nckx30C
43.6/67
N0.HOG0012074 Dple g10784.t2 Nckx30C
N0.HOG0012074 Ccro g1893.t2 Nckx30C
N0.HOG0012074 Sbem g4115.t1 Nckx30C
N0.HOG0012074 Pmal g16823.t1 Nckx30C
N0.HOG0012074 Tsyl g16005.t1 Nckx30C
N0.HOG0012074 Mleu g11892.t1 Nckx30C
90.3/93
N0.HOG0012074 Bmor g9762.t1 Nckx30C
75.6/43
N0.HOG0012074 Bbet g16245.t1 Nckx30C
N0.HOG0012074 Pxut g6092.t1 Nckx30C
99.2/100
N0.HOG0012074 Pmac g8561.t1 Nckx30C
75.8/38
N0.HOG0012074 Pint g3551.t2 Nckx30C
85.9/28
100/100
N0.HOG0012074 Zfil g12696.t1 Nckx30C
N0.HOG0012074 Hpyr g1052.t2 Nckx30C
69/71
N0.HOG0012074 Aips g7346.t2 Nckx30C
N0.HOG0016013 Mleu g11889.t1 Nckx30C
73.4/63
N0.HOG0016013 Cvir g4373.t1 Nckx30C
99.6/99
N0.HOG0016013 Cnem g2271.t1 Nckx30C
74.6/83
N0.HOG0016013 Bbet g16247.t1 Nckx30C
N0.HOG0016013 Sbem g4117.t1 Nckx30C
N0.HOG0016013 Msex g14264.t1 Nckx30C
0/31
N0.HOG0016013 Lcor g359.t1 Nckx30C
86.5/61
N0.HOG0016013 Pxut g6093.t1 Nckx30C
74.7/87
N0.HOG0016013 Pmac g8567.t1 Nckx30C
88.6/90
N0.HOG0016013 Zfil g12698.t1 Nckx30C
53.5/69
N0.HOG0016013 Bmor g9764.t1 Nckx30C

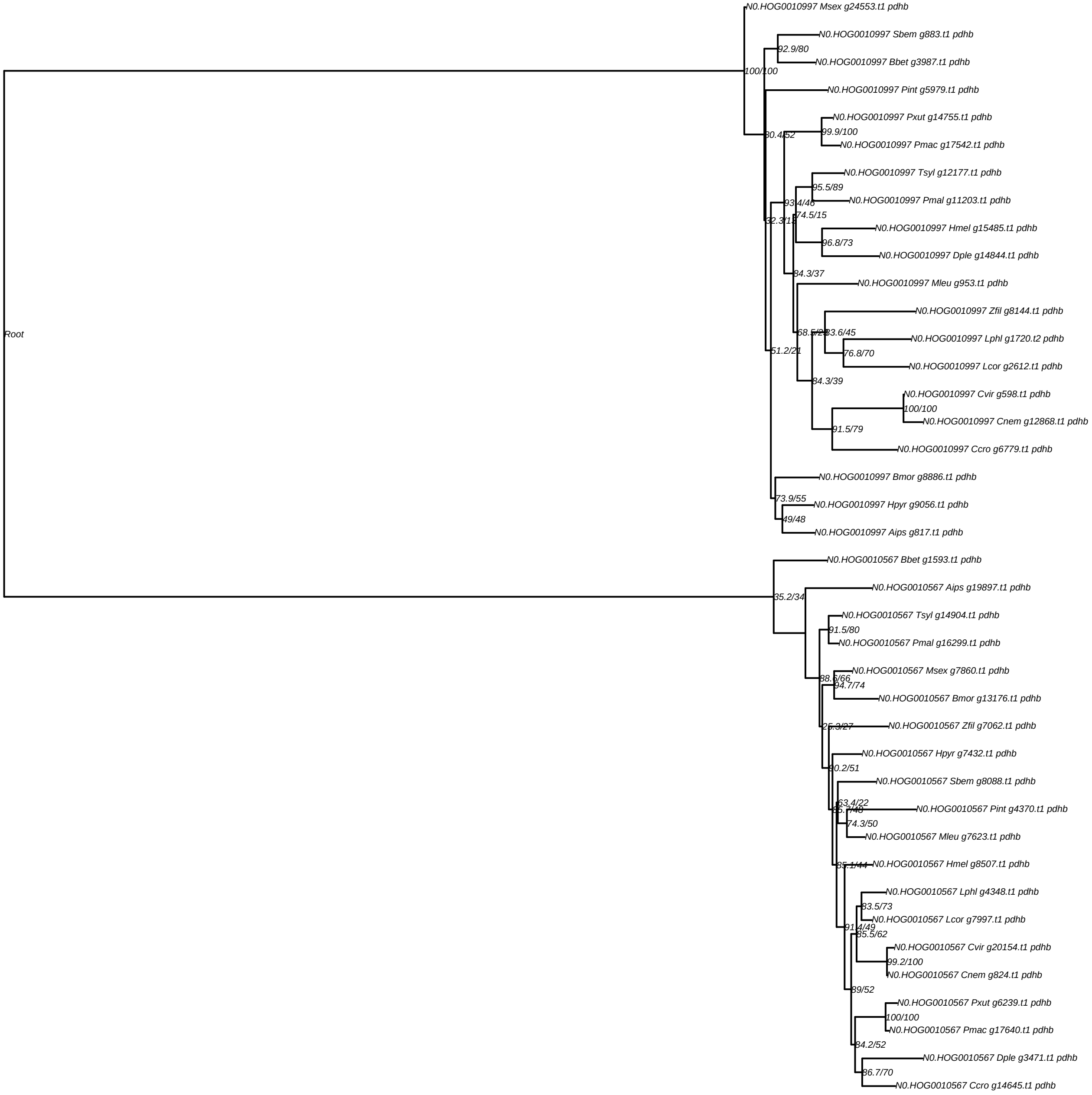
N0.HOG0010997 Msex g24553.t1 pdhb
N0.HOG0010997 Sbem g883.t1 pdhb
92.9/80
N0.HOG0010997 Bbet g3987.t1 pdhb
100/100
N0.HOG0010997 Pint g5979.t1 pdhb
N0.HOG0010997 Pxut g14755.t1 pdhb
99.9/100
80.4/52
N0.HOG0010997 Pmac g17542.t1 pdhb
N0.HOG0010997 Tsyl g12177.t1 pdhb
95.5/89
N0.HOG0010997 Pmal g11203.t1 pdhb
93.4/46
74.5/15
32.3/15
N0.HOG0010997 Hmel g15485.t1 pdhb
96.8/73
N0.HOG0010997 Dple g14844.t1 pdhb
84.3/37
N0.HOG0010997 Mleu g953.t1 pdhb
N0.HOG0010997 Zfil g8144.t1 pdhb
68.5/28
83.6/45
N0.HOG0010997 Lphl g1720.t2 pdhb
76.8/70
51.2/21
N0.HOG0010997 Lcor g2612.t1 pdhb
84.3/39
N0.HOG0010997 Cvir g598.t1 pdhb
100/100
N0.HOG0010997 Cnem g12868.t1 pdhb
91.5/79
N0.HOG0010997 Ccro g6779.t1 pdhb
N0.HOG0010997 Bmor g8886.t1 pdhb
73.9/55
N0.HOG0010997 Hpyr g9056.t1 pdhb
49/48
N0.HOG0010997 Aips g817.t1 pdhb
Root
N0.HOG0010567 Bbet g1593.t1 pdhb
N0.HOG0010567 Aips g19897.t1 pdhb
35.2/34
N0.HOG0010567 Tsyl g14904.t1 pdhb
91.5/80
N0.HOG0010567 Pmal g16299.t1 pdhb
N0.HOG0010567 Msex g7860.t1 pdhb
88.6/66
94.7/74
N0.HOG0010567 Bmor g13176.t1 pdhb
N0.HOG0010567 Zfil g7062.t1 pdhb
25.3/27
N0.HOG0010567 Hpyr g7432.t1 pdhb
90.2/51
N0.HOG0010567 Sbem g8088.t1 pdhb
63.4/22
85.7/48
N0.HOG0010567 Pint g4370.t1 pdhb
74.3/50
N0.HOG0010567 Mleu g7623.t1 pdhb
85.1/44
N0.HOG0010567 Hmel g8507.t1 pdhb
N0.HOG0010567 Lphl g4348.t1 pdhb
83.5/73
N0.HOG0010567 Lcor g7997.t1 pdhb
91.4/49
85.5/62
N0.HOG0010567 Cvir g20154.t1 pdhb
99.2/100
N0.HOG0010567 Cnem g824.t1 pdhb
89/52
N0.HOG0010567 Pxut g6239.t1 pdhb
100/100
N0.HOG0010567 Pmac g17640.t1 pdhb
84.2/52
N0.HOG0010567 Dple g3471.t1 pdhb
86.7/70
N0.HOG0010567 Ccro g14645.t1 pdhb

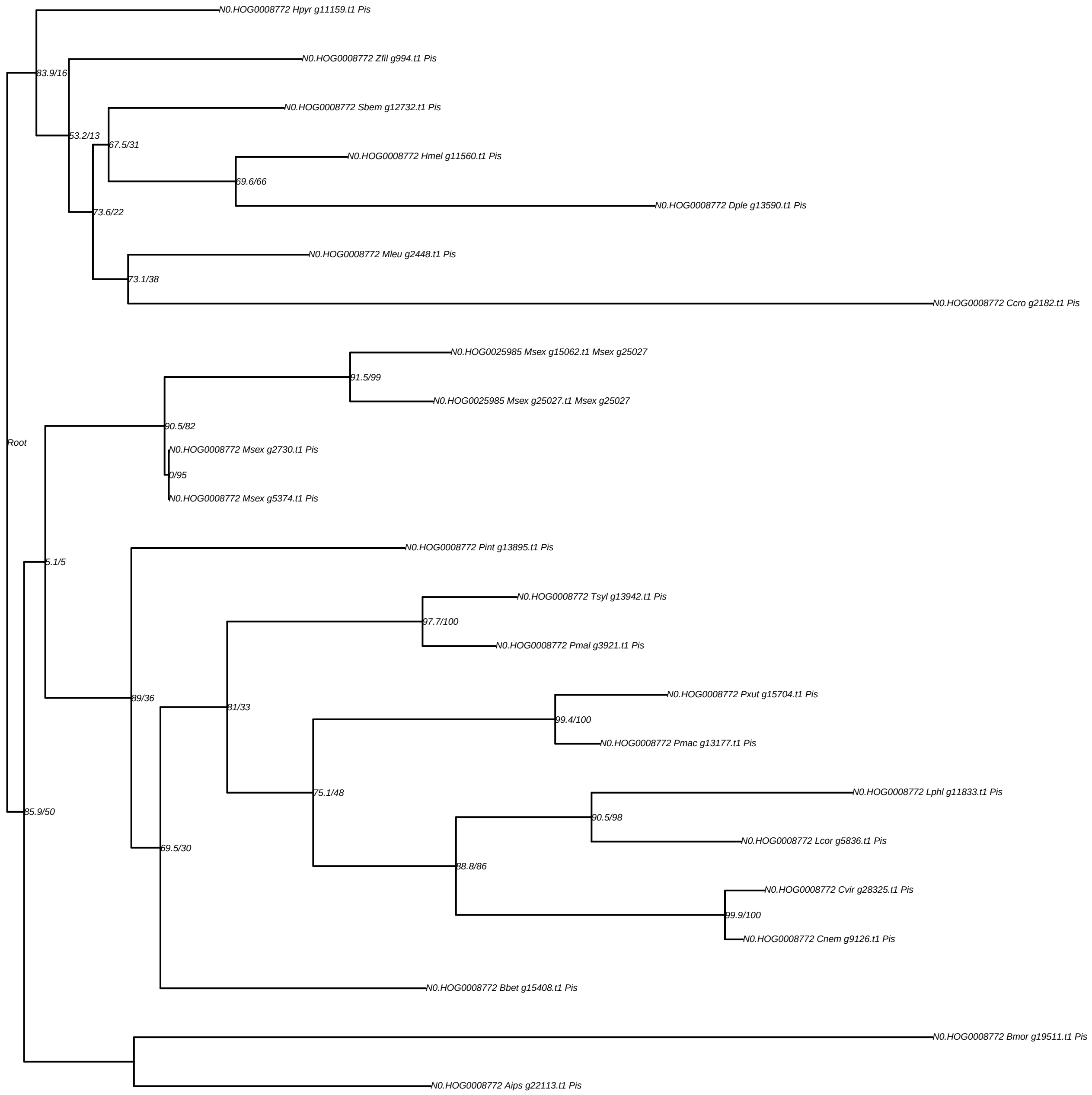
N0.HOG0008772 Hpyr g11159.t1 Pis
83.9/16
N0.HOG0008772 Zfil g994.t1 Pis
N0.HOG0008772 Sbem g12732.t1 Pis
53.2/13
67.5/31
N0.HOG0008772 Hmel g11560.t1 Pis
69.6/66
N0.HOG0008772 Dple g13590.t1 Pis
73.6/22
N0.HOG0008772 Mleu g2448.t1 Pis
73.1/38
N0.HOG0008772 Ccro g2182.t1 Pis
N0.HOG0025985 Msex g15062.t1 Msex g25027
91.5/99
N0.HOG0025985 Msex g25027.t1 Msex g25027
90.5/82
Root
N0.HOG0008772 Msex g2730.t1 Pis
0/95
N0.HOG0008772 Msex g5374.t1 Pis
N0.HOG0008772 Pint g13895.t1 Pis
5.1/5
N0.HOG0008772 Tsyl g13942.t1 Pis
97.7/100
N0.HOG0008772 Pmal g3921.t1 Pis
N0.HOG0008772 Pxut g15704.t1 Pis
89/36
81/33
99.4/100
N0.HOG0008772 Pmac g13177.t1 Pis
N0.HOG0008772 Lphl g11833.t1 Pis
75.1/48
85.9/50
90.5/98
N0.HOG0008772 Lcor g5836.t1 Pis
69.5/30
88.8/86
N0.HOG0008772 Cvir g28325.t1 Pis
99.9/100
N0.HOG0008772 Cnem g9126.t1 Pis
N0.HOG0008772 Bbet g15408.t1 Pis
N0.HOG0008772 Bmor g19511.t1 Pis
N0.HOG0008772 Aips g22113.t1 Pis

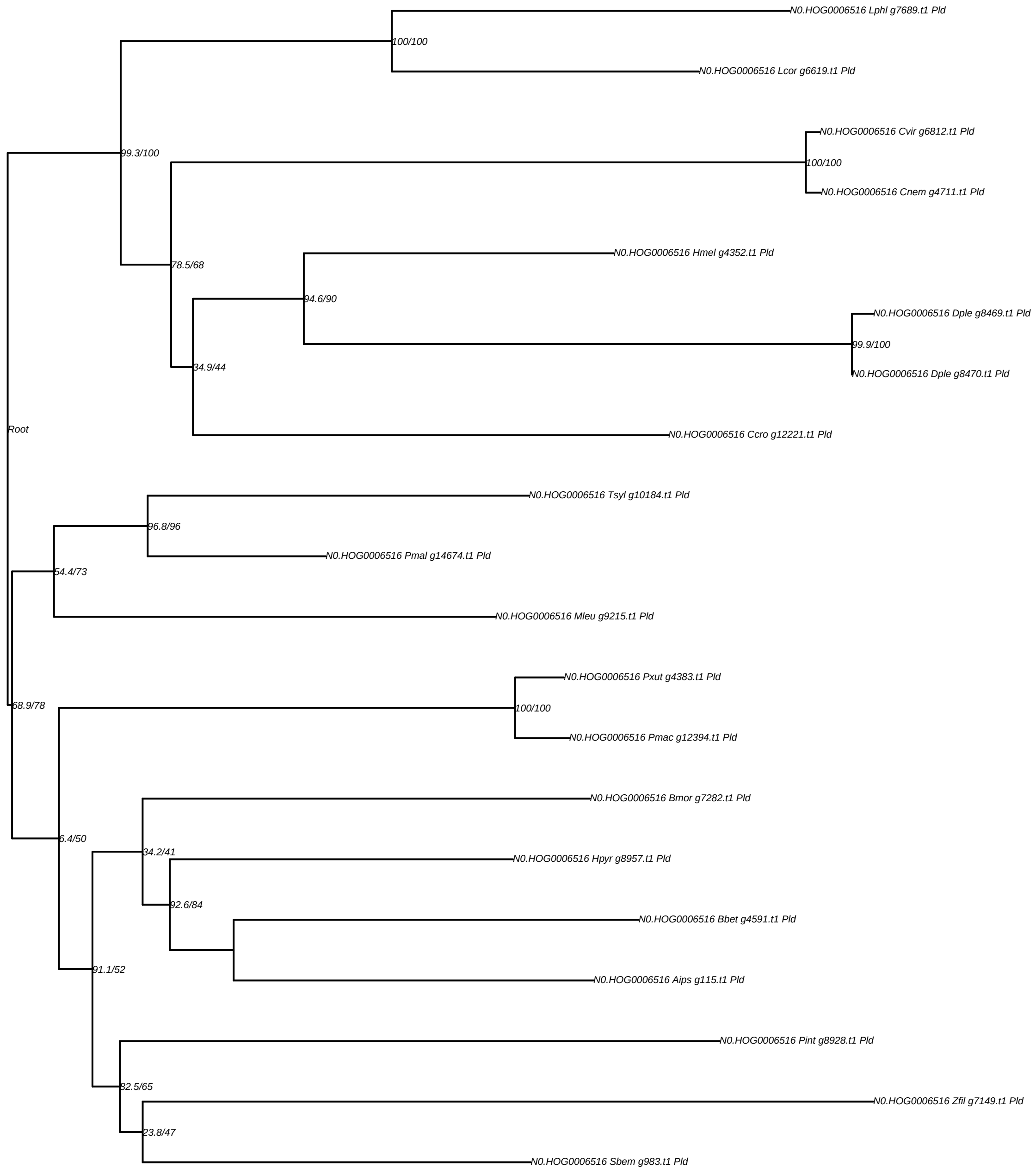
N0.HOG0006516 Lphl g7689.t1 Pld
100/100
N0.HOG0006516 Lcor g6619.t1 Pld
N0.HOG0006516 Cvir g6812.t1 Pld
99.3/100
100/100
N0.HOG0006516 Cnem g4711.t1 Pld
N0.HOG0006516 Hmel g4352.t1 Pld
78.5/68
94.6/90
N0.HOG0006516 Dple g8469.t1 Pld
99.9/100
34.9/44
N0.HOG0006516 Dple g8470.t1 Pld
Root
N0.HOG0006516 Ccro g12221.t1 Pld
N0.HOG0006516 Tsyl g10184.t1 Pld
96.8/96
N0.HOG0006516 Pmal g14674.t1 Pld
54.4/73
N0.HOG0006516 Mleu g9215.t1 Pld
N0.HOG0006516 Pxut g4383.t1 Pld
68.9/78
100/100
N0.HOG0006516 Pmac g12394.t1 Pld
N0.HOG0006516 Bmor g7282.t1 Pld
6.4/50
34.2/41
N0.HOG0006516 Hpyr g8957.t1 Pld
92.6/84
N0.HOG0006516 Bbet g4591.t1 Pld
91.1/52
N0.HOG0006516 Aips g115.t1 Pld
N0.HOG0006516 Pint g8928.t1 Pld
82.5/65
N0.HOG0006516 Zfil g7149.t1 Pld
23.8/47
N0.HOG0006516 Sbem g983.t1 Pld

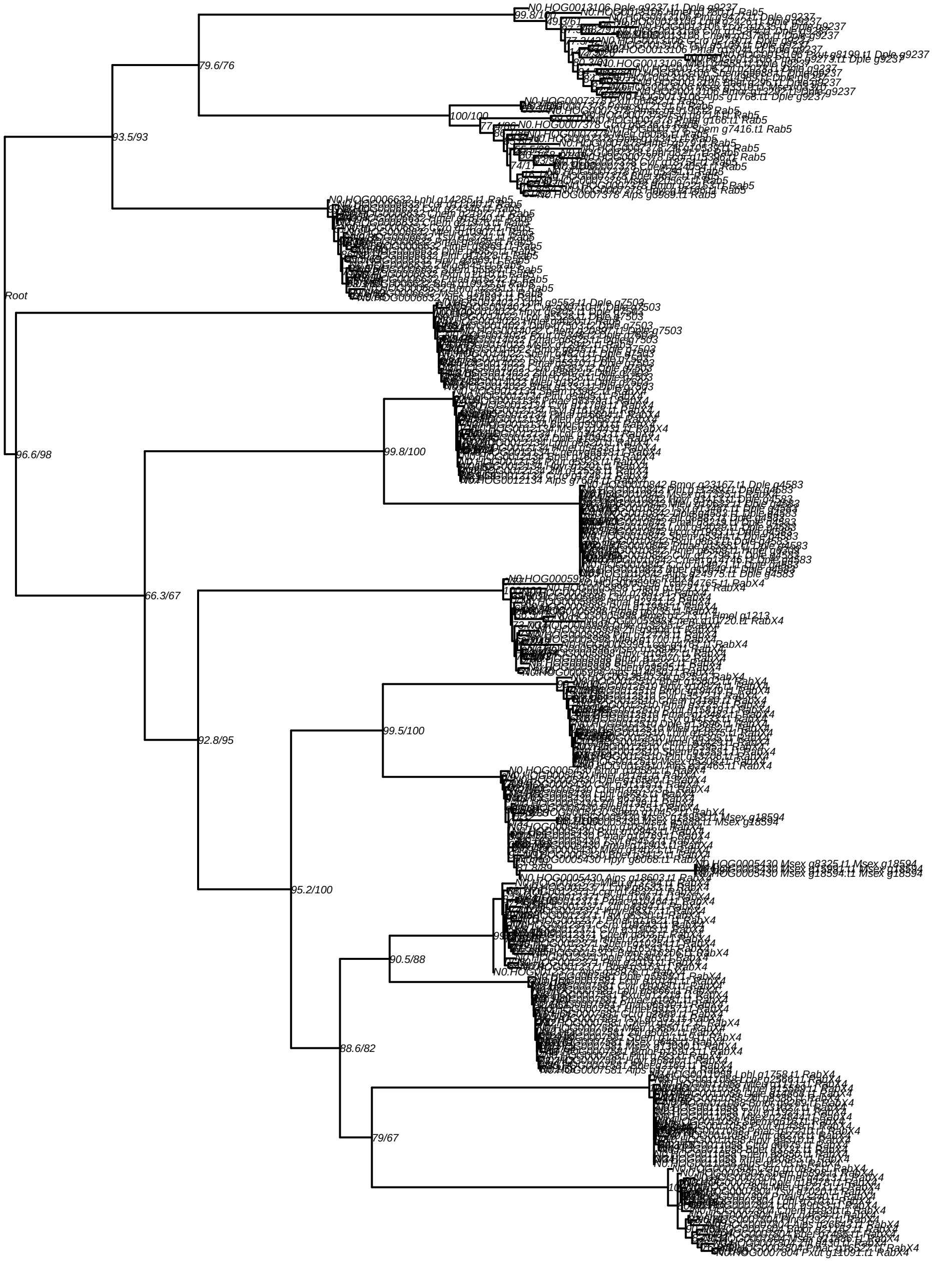
Root
93.5/93
79.6/76
96.6/98
66.3/67
99.8/100
92.8/95
99.5/100
95.2/100
90.5/88
88.6/82
79/67
100/100
N0.HOG0013106 Dple g9237.t1 Dple g9237
N0.HOG0006632 Lphl g14285.t1 Rab5
N0.HOG0007378 Aips g6969.t1 Rab5
N0.HOG0005430 Msex g13594.t1 Msex g18594
N0.HOG0007804 Pxut g11091.t1 RabX4

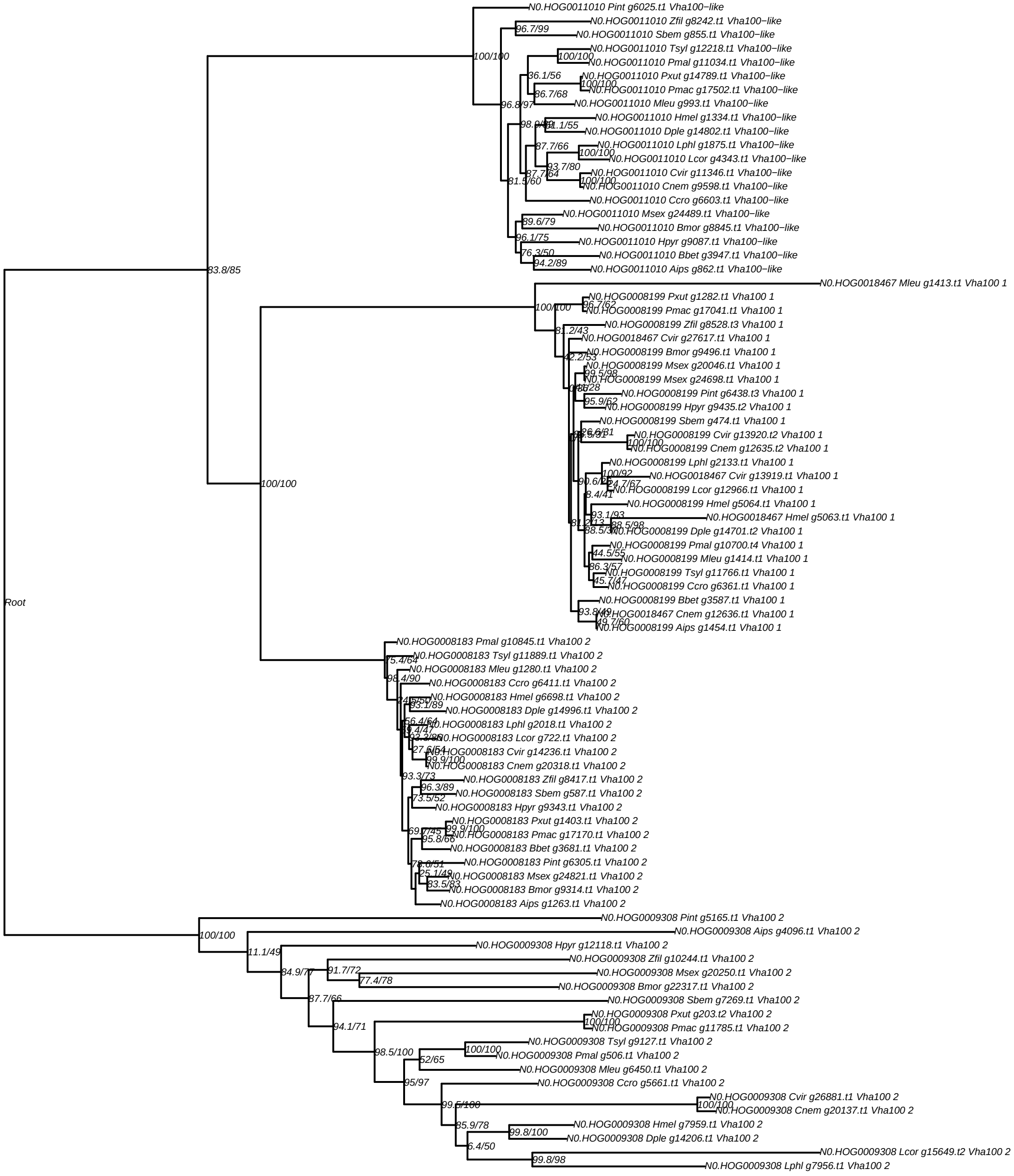
N0.HOG0011010 Pint g6025.t1 Vha100−like
N0.HOG0011010 Zfil g8242.t1 Vha100−like
N0.HOG0011010 Sbem g855.t1 Vha100−like
N0.HOG0011010 Tsyl g12218.t1 Vha100−like
N0.HOG0011010 Pmal g11034.t1 Vha100−like
N0.HOG0011010 Pxut g14789.t1 Vha100−like
N0.HOG0011010 Pmac g17502.t1 Vha100−like
N0.HOG0011010 Mleu g993.t1 Vha100−like
N0.HOG0011010 Hmel g1334.t1 Vha100−like
N0.HOG0011010 Dple g14802.t1 Vha100−like
N0.HOG0011010 Lphl g1875.t1 Vha100−like
N0.HOG0011010 Lcor g4343.t1 Vha100−like
N0.HOG0011010 Cvir g11346.t1 Vha100−like
N0.HOG0011010 Cnem g9598.t1 Vha100−like
N0.HOG0011010 Ccro g6603.t1 Vha100−like
N0.HOG0011010 Msex g24489.t1 Vha100−like
N0.HOG0011010 Bmor g8845.t1 Vha100−like
N0.HOG0011010 Hpyr g9087.t1 Vha100−like
N0.HOG0011010 Bbet g3947.t1 Vha100−like
N0.HOG0011010 Aips g862.t1 Vha100−like
N0.HOG0018467 Mleu g1413.t1 Vha100 1
N0.HOG0008199 Pxut g1282.t1 Vha100 1
N0.HOG0008199 Pmac g17041.t1 Vha100 1
N0.HOG0008199 Zfil g8528.t3 Vha100 1
N0.HOG0018467 Cvir g27617.t1 Vha100 1
N0.HOG0008199 Bmor g9496.t1 Vha100 1
N0.HOG0008199 Msex g20046.t1 Vha100 1
N0.HOG0008199 Msex g24698.t1 Vha100 1
N0.HOG0008199 Pint g6438.t3 Vha100 1
N0.HOG0008199 Hpyr g9435.t2 Vha100 1
N0.HOG0008199 Sbem g474.t1 Vha100 1
N0.HOG0008199 Cvir g13920.t2 Vha100 1
N0.HOG0008199 Cnem g12635.t2 Vha100 1
N0.HOG0008199 Lphl g2133.t1 Vha100 1
N0.HOG0018467 Cvir g13919.t1 Vha100 1
N0.HOG0008199 Lcor g12966.t1 Vha100 1
N0.HOG0008199 Hmel g5064.t1 Vha100 1
N0.HOG0018467 Hmel g5063.t1 Vha100 1
N0.HOG0008199 Dple g14701.t2 Vha100 1
N0.HOG0008199 Pmal g10700.t4 Vha100 1
N0.HOG0008199 Mleu g1414.t1 Vha100 1
N0.HOG0008199 Tsyl g11766.t1 Vha100 1
N0.HOG0008199 Ccro g6361.t1 Vha100 1
N0.HOG0008199 Bbet g3587.t1 Vha100 1
N0.HOG0018467 Cnem g12636.t1 Vha100 1
N0.HOG0008199 Aips g1454.t1 Vha100 1
N0.HOG0008183 Pmal g10845.t1 Vha100 2
N0.HOG0008183 Tsyl g11889.t1 Vha100 2
N0.HOG0008183 Mleu g1280.t1 Vha100 2
N0.HOG0008183 Ccro g6411.t1 Vha100 2
N0.HOG0008183 Hmel g6698.t1 Vha100 2
N0.HOG0008183 Dple g14996.t1 Vha100 2
N0.HOG0008183 Lphl g2018.t1 Vha100 2
N0.HOG0008183 Lcor g722.t1 Vha100 2
N0.HOG0008183 Cvir g14236.t1 Vha100 2
N0.HOG0008183 Cnem g20318.t1 Vha100 2
N0.HOG0008183 Zfil g8417.t1 Vha100 2
N0.HOG0008183 Sbem g587.t1 Vha100 2
N0.HOG0008183 Hpyr g9343.t1 Vha100 2
N0.HOG0008183 Pxut g1403.t1 Vha100 2
N0.HOG0008183 Pmac g17170.t1 Vha100 2
N0.HOG0008183 Bbet g3681.t1 Vha100 2
N0.HOG0008183 Pint g6305.t1 Vha100 2
N0.HOG0008183 Msex g24821.t1 Vha100 2
N0.HOG0008183 Bmor g9314.t1 Vha100 2
N0.HOG0008183 Aips g1263.t1 Vha100 2
N0.HOG0009308 Pint g5165.t1 Vha100 2
N0.HOG0009308 Aips g4096.t1 Vha100 2
N0.HOG0009308 Hpyr g12118.t1 Vha100 2
N0.HOG0009308 Zfil g10244.t1 Vha100 2
N0.HOG0009308 Msex g20250.t1 Vha100 2
N0.HOG0009308 Bmor g22317.t1 Vha100 2
N0.HOG0009308 Sbem g7269.t1 Vha100 2
N0.HOG0009308 Pxut g203.t2 Vha100 2
N0.HOG0009308 Pmac g11785.t1 Vha100 2
N0.HOG0009308 Tsyl g9127.t1 Vha100 2
N0.HOG0009308 Pmal g506.t1 Vha100 2
N0.HOG0009308 Mleu g6450.t1 Vha100 2
N0.HOG0009308 Ccro g5661.t1 Vha100 2
N0.HOG0009308 Cvir g26881.t1 Vha100 2
N0.HOG0009308 Cnem g20137.t1 Vha100 2
N0.HOG0009308 Hmel g7959.t1 Vha100 2
N0.HOG0009308 Dple g14206.t1 Vha100 2
N0.HOG0009308 Lcor g15649.t2 Vha100 2
N0.HOG0009308 Lphl g7956.t1 Vha100 2
Root

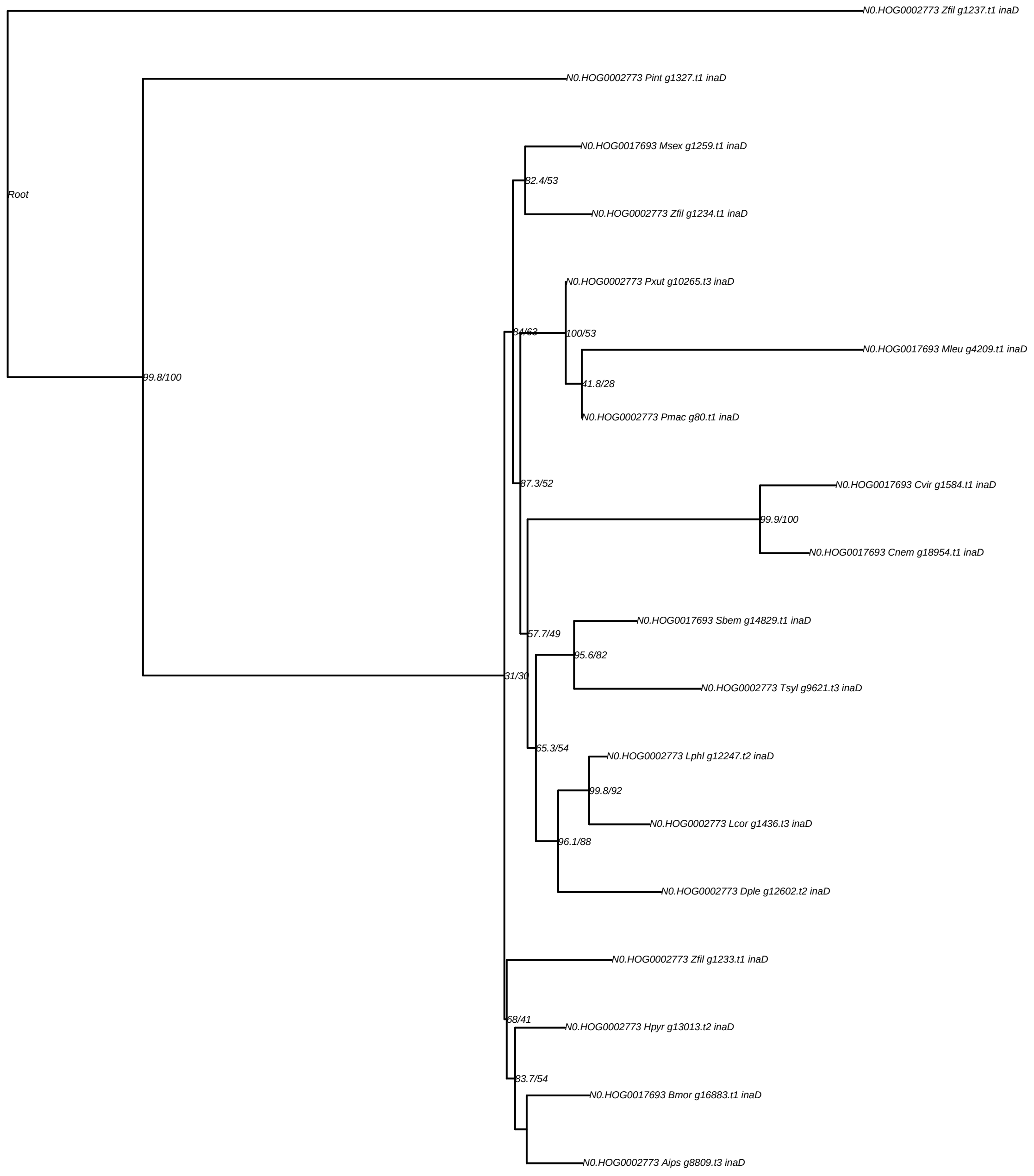
N0.HOG0002773 Zfil g1237.t1 inaD
Root
N0.HOG0002773 Pint g1327.t1 inaD
N0.HOG0017693 Msex g1259.t1 inaD
82.4/53
N0.HOG0002773 Zfil g1234.t1 inaD
N0.HOG0002773 Pxut g10265.t3 inaD
84/63
100/53
N0.HOG0017693 Mleu g4209.t1 inaD
99.8/100
41.8/28
N0.HOG0002773 Pmac g80.t1 inaD
87.3/52
N0.HOG0017693 Cvir g1584.t1 inaD
99.9/100
N0.HOG0017693 Cnem g18954.t1 inaD
N0.HOG0017693 Sbem g14829.t1 inaD
57.7/49
95.6/82
31/30
N0.HOG0002773 Tsyl g9621.t3 inaD
65.3/54
N0.HOG0002773 Lphl g12247.t2 inaD
99.8/92
N0.HOG0002773 Lcor g1436.t3 inaD
96.1/88
N0.HOG0002773 Dple g12602.t2 inaD
N0.HOG0002773 Zfil g1233.t1 inaD
68/41
N0.HOG0002773 Hpyr g13013.t2 inaD
83.7/54
N0.HOG0017693 Bmor g16883.t1 inaD
N0.HOG0002773 Aips g8809.t3 inaD

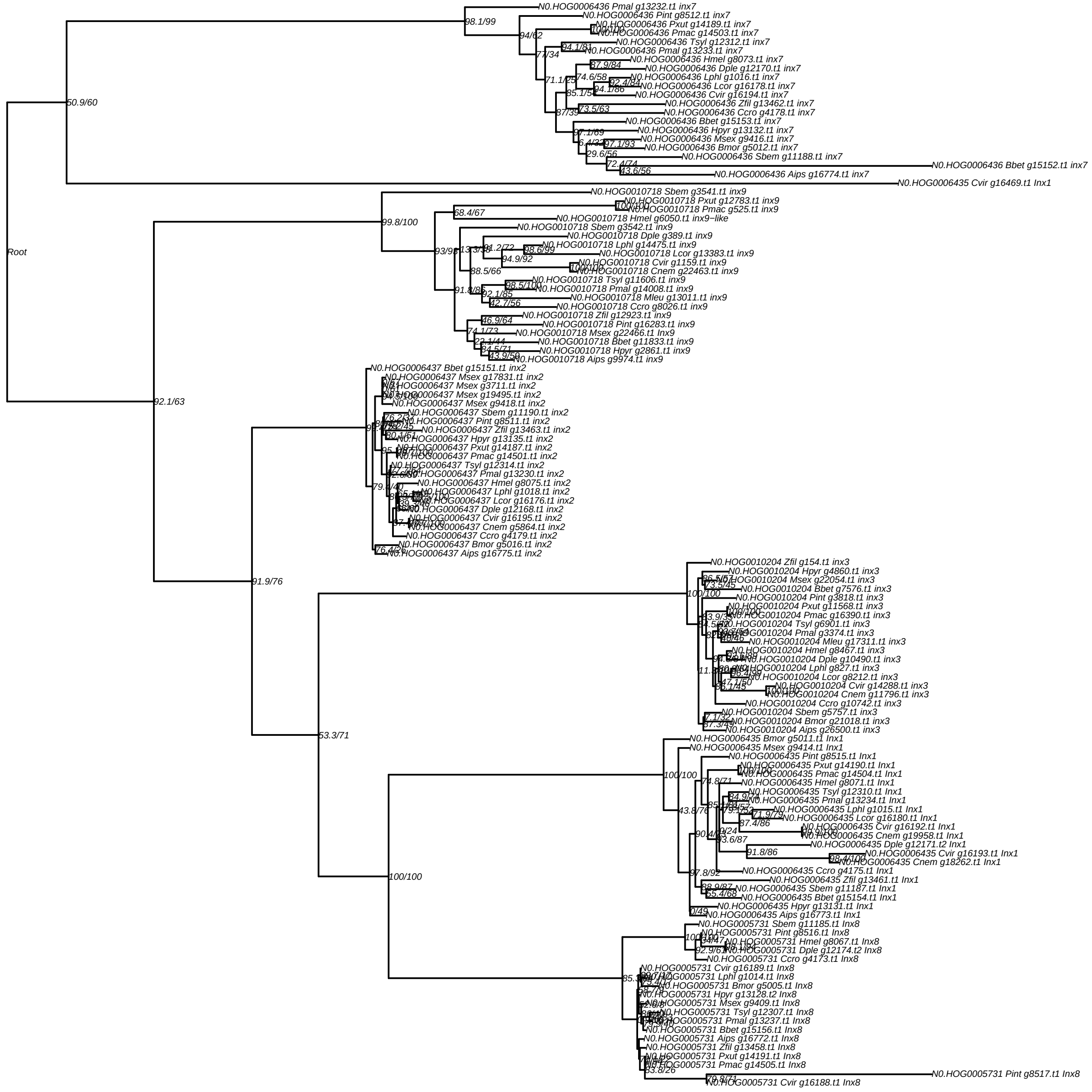
Root
N0.HOG0006436 Pmal g13232.t1 inx7
N0.HOG0006436 Pint g8512.t1 inx7
N0.HOG0006436 Pxut g14189.t1 inx7
N0.HOG0006436 Pmac g14503.t1 inx7
N0.HOG0006436 Tsyl g12312.t1 inx7
N0.HOG0006436 Pmal g13233.t1 inx7
N0.HOG0006436 Hmel g8073.t1 inx7
N0.HOG0006436 Dple g12170.t1 inx7
N0.HOG0006436 Lphl g1016.t1 inx7
N0.HOG0006436 Lcor g16178.t1 inx7
N0.HOG0006436 Cvir g16194.t1 inx7
N0.HOG0006436 Zfil g13462.t1 inx7
N0.HOG0006436 Ccro g4178.t1 inx7
N0.HOG0006436 Bbet g15153.t1 inx7
N0.HOG0006436 Hpyr g13132.t1 inx7
N0.HOG0006436 Msex g9416.t1 inx7
N0.HOG0006436 Bmor g5012.t1 inx7
N0.HOG0006436 Sbem g11188.t1 inx7
N0.HOG0006436 Bbet g15152.t1 inx7
N0.HOG0006436 Aips g16774.t1 inx7
N0.HOG0006435 Cvir g16469.t1 Inx1
N0.HOG0010718 Sbem g3541.t1 inx9
N0.HOG0010718 Pxut g12783.t1 inx9
N0.HOG0010718 Pmac g525.t1 inx9
N0.HOG0010718 Hmel g6050.t1 inx9−like
N0.HOG0010718 Sbem g3542.t1 inx9
N0.HOG0010718 Dple g389.t1 inx9
N0.HOG0010718 Lphl g14475.t1 inx9
N0.HOG0010718 Lcor g13383.t1 inx9
N0.HOG0010718 Cvir g1159.t1 inx9
N0.HOG0010718 Cnem g22463.t1 inx9
N0.HOG0010718 Tsyl g11606.t1 inx9
N0.HOG0010718 Pmal g14008.t1 inx9
N0.HOG0010718 Mleu g13011.t1 inx9
N0.HOG0010718 Ccro g8026.t1 inx9
N0.HOG0010718 Zfil g12923.t1 inx9
N0.HOG0010718 Pint g16283.t1 inx9
N0.HOG0010718 Msex g22466.t1 Inx9
N0.HOG0010718 Bbet g11833.t1 inx9
N0.HOG0010718 Hpyr g2861.t1 inx9
N0.HOG0010718 Aips g9974.t1 inx9
N0.HOG0006437 Bbet g15151.t1 inx2
N0.HOG0006437 Msex g17831.t1 inx2
N0.HOG0006437 Msex g3711.t1 inx2
N0.HOG0006437 Msex g19495.t1 inx2
N0.HOG0006437 Msex g9418.t1 inx2
N0.HOG0006437 Sbem g11190.t1 inx2
N0.HOG0006437 Pint g8511.t1 inx2
N0.HOG0006437 Zfil g13463.t1 inx2
N0.HOG0006437 Hpyr g13135.t1 inx2
N0.HOG0006437 Pxut g14187.t1 inx2
N0.HOG0006437 Pmac g14501.t1 inx2
N0.HOG0006437 Tsyl g12314.t1 inx2
N0.HOG0006437 Pmal g13230.t1 inx2
N0.HOG0006437 Hmel g8075.t1 inx2
N0.HOG0006437 Lphl g1018.t1 inx2
N0.HOG0006437 Lcor g16176.t1 inx2
N0.HOG0006437 Dple g12168.t1 inx2
N0.HOG0006437 Cvir g16195.t1 inx2
N0.HOG0006437 Cnem g5864.t1 inx2
N0.HOG0006437 Ccro g4179.t1 inx2
N0.HOG0006437 Bmor g5016.t1 inx2
N0.HOG0006437 Aips g16775.t1 inx2
N0.HOG0010204 Zfil g154.t1 inx3
N0.HOG0010204 Hpyr g4860.t1 inx3
N0.HOG0010204 Msex g22054.t1 inx3
N0.HOG0010204 Bbet g7576.t1 inx3
N0.HOG0010204 Pint g3818.t1 inx3
N0.HOG0010204 Pxut g11568.t1 inx3
N0.HOG0010204 Pmac g16390.t1 inx3
N0.HOG0010204 Tsyl g6901.t1 inx3
N0.HOG0010204 Pmal g3374.t1 inx3
N0.HOG0010204 Mleu g17311.t1 inx3
N0.HOG0010204 Hmel g8467.t1 inx3
N0.HOG0010204 Dple g10490.t1 inx3
N0.HOG0010204 Lphl g827.t1 inx3
N0.HOG0010204 Lcor g8212.t1 inx3
N0.HOG0010204 Cvir g14288.t1 inx3
N0.HOG0010204 Cnem g11796.t1 inx3
N0.HOG0010204 Ccro g10742.t1 inx3
N0.HOG0010204 Sbem g5757.t1 inx3
N0.HOG0010204 Bmor g21018.t1 inx3
N0.HOG0010204 Aips g26500.t1 inx3
N0.HOG0006435 Bmor g5011.t1 Inx1
N0.HOG0006435 Msex g9414.t1 Inx1
N0.HOG0006435 Pint g8515.t1 Inx1
N0.HOG0006435 Pxut g14190.t1 Inx1
N0.HOG0006435 Pmac g14504.t1 Inx1
N0.HOG0006435 Hmel g8071.t1 Inx1
N0.HOG0006435 Tsyl g12310.t1 Inx1
N0.HOG0006435 Pmal g13234.t1 Inx1
N0.HOG0006435 Lphl g1015.t1 Inx1
N0.HOG0006435 Lcor g16180.t1 Inx1
N0.HOG0006435 Cvir g16192.t1 Inx1
N0.HOG0006435 Cnem g19958.t1 Inx1
N0.HOG0006435 Dple g12171.t2 Inx1
N0.HOG0006435 Cvir g16193.t1 Inx1
N0.HOG0006435 Cnem g18262.t1 Inx1
N0.HOG0006435 Ccro g4175.t1 Inx1
N0.HOG0006435 Zfil g13461.t1 Inx1
N0.HOG0006435 Sbem g11187.t1 Inx1
N0.HOG0006435 Bbet g15154.t1 Inx1
N0.HOG0006435 Hpyr g13131.t1 Inx1
N0.HOG0006435 Aips g16773.t1 Inx1
N0.HOG0005731 Sbem g11185.t1 Inx8
N0.HOG0005731 Pint g8516.t1 Inx8
N0.HOG0005731 Hmel g8067.t1 Inx8
N0.HOG0005731 Dple g12174.t2 Inx8
N0.HOG0005731 Ccro g4173.t1 Inx8
N0.HOG0005731 Cvir g16189.t1 Inx8
N0.HOG0005731 Lphl g1014.t1 Inx8
N0.HOG0005731 Bmor g5005.t1 Inx8
N0.HOG0005731 Hpyr g13128.t2 Inx8
N0.HOG0005731 Msex g9409.t1 Inx8
N0.HOG0005731 Tsyl g12307.t1 Inx8
N0.HOG0005731 Pmal g13237.t1 Inx8
N0.HOG0005731 Bbet g15156.t1 Inx8
N0.HOG0005731 Aips g16772.t1 Inx8
N0.HOG0005731 Zfil g13458.t1 Inx8
N0.HOG0005731 Pxut g14191.t1 Inx8
N0.HOG0005731 Pmac g14505.t1 Inx8
N0.HOG0005731 Pint g8517.t1 Inx8
N0.HOG0005731 Cvir g16188.t1 Inx8

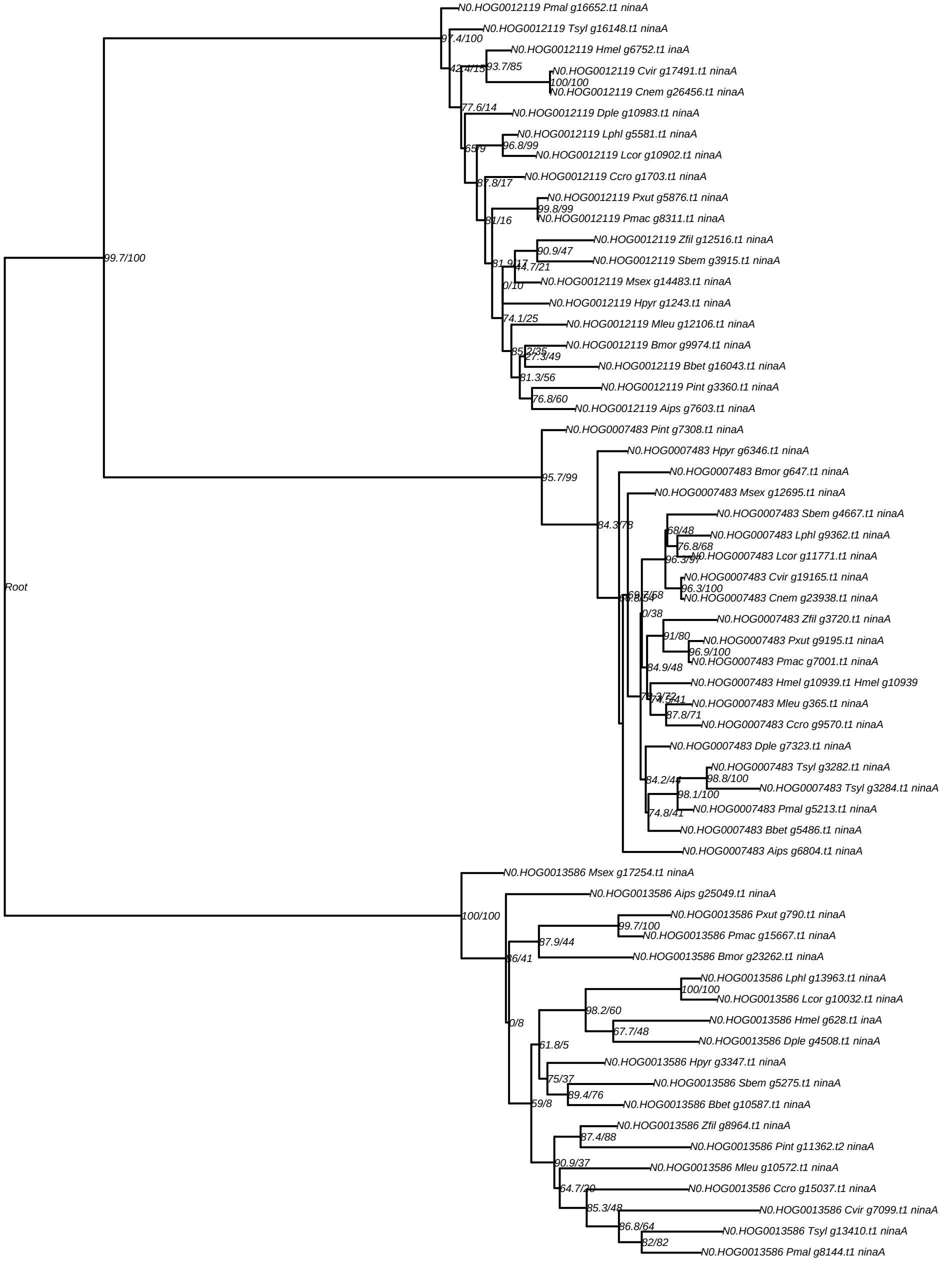

N0.HOG0012119 Pmal g16652.t1 ninaA
N0.HOG0012119 Tsyl g16148.t1 ninaA
97.4/100
N0.HOG0012119 Hmel g6752.t1 inaA
42.4/15
93.7/85
N0.HOG0012119 Cvir g17491.t1 ninaA
100/100
N0.HOG0012119 Cnem g26456.t1 ninaA
77.6/14
N0.HOG0012119 Dple g10983.t1 ninaA
N0.HOG0012119 Lphl g5581.t1 ninaA
65/9
96.8/99
N0.HOG0012119 Lcor g10902.t1 ninaA
N0.HOG0012119 Ccro g1703.t1 ninaA
87.8/17
N0.HOG0012119 Pxut g5876.t1 ninaA
99.8/99
81/16
N0.HOG0012119 Pmac g8311.t1 ninaA
N0.HOG0012119 Zfil g12516.t1 ninaA
90.9/47
99.7/100
81.9/17
44.7/21
N0.HOG0012119 Sbem g3915.t1 ninaA
0/10
N0.HOG0012119 Msex g14483.t1 ninaA
N0.HOG0012119 Hpyr g1243.t1 ninaA
74.1/25
N0.HOG0012119 Mleu g12106.t1 ninaA
N0.HOG0012119 Bmor g9974.t1 ninaA
85.2/35
27.3/49
N0.HOG0012119 Bbet g16043.t1 ninaA
81.3/56
N0.HOG0012119 Pint g3360.t1 ninaA
76.8/60
N0.HOG0012119 Aips g7603.t1 ninaA
N0.HOG0007483 Pint g7308.t1 ninaA
N0.HOG0007483 Hpyr g6346.t1 ninaA
N0.HOG0007483 Bmor g647.t1 ninaA
95.7/99
N0.HOG0007483 Msex g12695.t1 ninaA
N0.HOG0007483 Sbem g4667.t1 ninaA
84.3/78
68/48
N0.HOG0007483 Lphl g9362.t1 ninaA
76.8/68
96.3/97
N0.HOG0007483 Lcor g11771.t1 ninaA
N0.HOG0007483 Cvir g19165.t1 ninaA
Root
96.3/100
69.7/68
N0.HOG0007483 Cnem g23938.t1 ninaA
0/38
N0.HOG0007483 Zfil g3720.t1 ninaA
91/80
N0.HOG0007483 Pxut g9195.t1 ninaA
96.9/100
N0.HOG0007483 Pmac g7001.t1 ninaA
84.9/48
N0.HOG0007483 Hmel g10939.t1 Hmel g10939
74.5/41
N0.HOG0007483 Mleu g365.t1 ninaA
87.8/71
N0.HOG0007483 Ccro g9570.t1 ninaA
N0.HOG0007483 Dple g7323.t1 ninaA
N0.HOG0007483 Tsyl g3282.t1 ninaA
84.2/44
98.8/100
N0.HOG0007483 Tsyl g3284.t1 ninaA
98.1/100
74.8/41
N0.HOG0007483 Pmal g5213.t1 ninaA
N0.HOG0007483 Bbet g5486.t1 ninaA
N0.HOG0007483 Aips g6804.t1 ninaA
N0.HOG0013586 Msex g17254.t1 ninaA
N0.HOG0013586 Aips g25049.t1 ninaA
N0.HOG0013586 Pxut g790.t1 ninaA
100/100
99.7/100
N0.HOG0013586 Pmac g15667.t1 ninaA
87.9/44
N0.HOG0013586 Bmor g23262.t1 ninaA
86/41
N0.HOG0013586 Lphl g13963.t1 ninaA
100/100
N0.HOG0013586 Lcor g10032.t1 ninaA
98.2/60
N0.HOG0013586 Hmel g628.t1 inaA
0/8
67.7/48
N0.HOG0013586 Dple g4508.t1 ninaA
61.8/5
N0.HOG0013586 Hpyr g3347.t1 ninaA
75/37
N0.HOG0013586 Sbem g5275.t1 ninaA
89.4/76
59/8
N0.HOG0013586 Bbet g10587.t1 ninaA
N0.HOG0013586 Zfil g8964.t1 ninaA
87.4/88
N0.HOG0013586 Pint g11362.t2 ninaA
90.9/37
N0.HOG0013586 Mleu g10572.t1 ninaA
64.7/20
N0.HOG0013586 Ccro g15037.t1 ninaA
85.3/48
N0.HOG0013586 Cvir g7099.t1 ninaA
86.8/64
N0.HOG0013586 Tsyl g13410.t1 ninaA
82/82
N0.HOG0013586 Pmal g8144.t1 ninaA

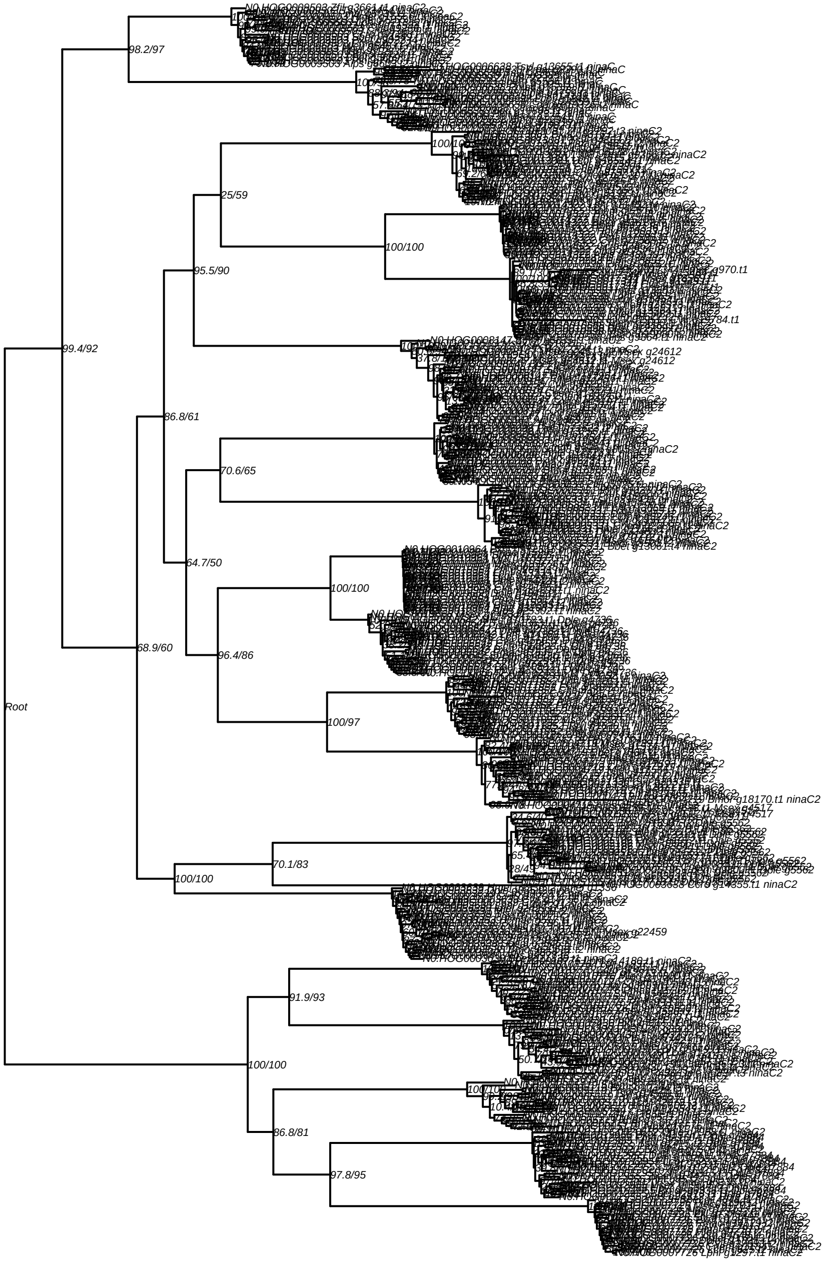

Root
99.4/92
98.2/97
68.9/60
86.8/61
95.5/90
25/59
100/100
64.7/50
70.6/65
96.4/86
100/100
100/97
100/100
70.1/83
100/100
91.9/93
86.8/81
97.8/95

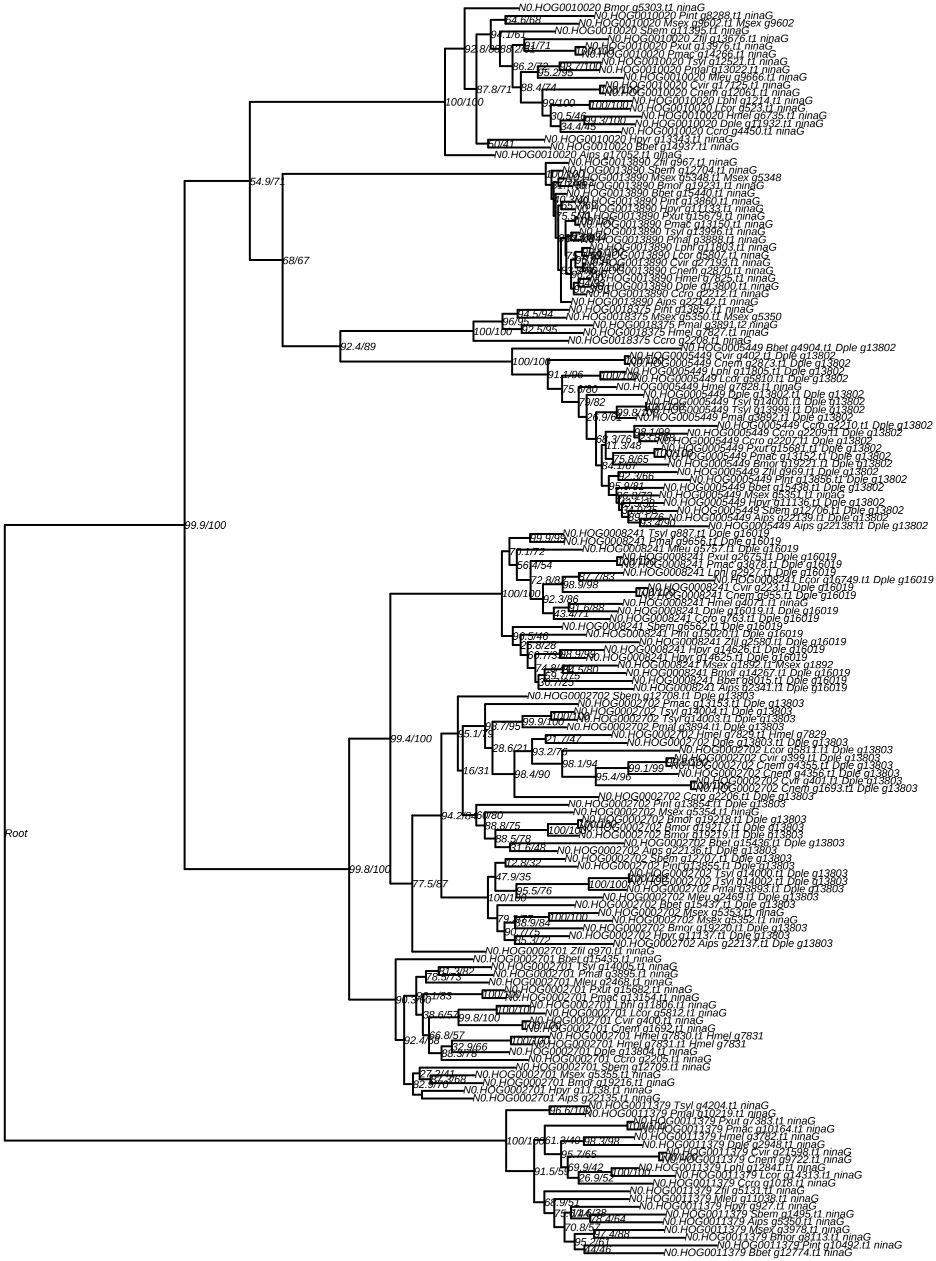
Root
99.9/100
54.9/71
68/67
100/100
92.4/89
99.8/100
99.4/100
77.5/87
N0.HOG0010020 Bmor g5303.t1 ninaG
N0.HOG0010020 Pint g8288.t1 ninaG
N0.HOG0010020 Msex g9602.t1 Msex g9602
N0.HOG0010020 Sbem g11395.t1 ninaG
N0.HOG0010020 Zfil g13676.t1 ninaG
N0.HOG0010020 Pxut g13976.t1 ninaG
N0.HOG0010020 Pmac g14266.t1 ninaG
N0.HOG0010020 Tsyl g12521.t1 ninaG
N0.HOG0010020 Pmal g13022.t1 ninaG
N0.HOG0010020 Mleu g9666.t1 ninaG
N0.HOG0010020 Cvir g17125.t1 ninaG
N0.HOG0010020 Cnem g12061.t1 ninaG
N0.HOG0010020 Lphl g1214.t1 ninaG
N0.HOG0010020 Lcor g523.t1 ninaG
N0.HOG0010020 Hmel g6735.t1 ninaG
N0.HOG0010020 Dple g11932.t1 ninaG
N0.HOG0010020 Ccro g4450.t1 ninaG
N0.HOG0010020 Hpyr g13343.t1 ninaG
N0.HOG0010020 Bbet g14937.t1 ninaG
N0.HOG0010020 Aips g17052.t1 ninaG
N0.HOG0013890 Zfil g967.t1 ninaG
N0.HOG0013890 Sbem g12704.t1 ninaG
N0.HOG0013890 Msex g5348.t1 Msex g5348
N0.HOG0013890 Bmor g19231.t1 ninaG
N0.HOG0013890 Bbet g15440.t1 ninaG
N0.HOG0013890 Pint g13860.t1 ninaG
N0.HOG0013890 Hpyr g11133.t1 ninaG
N0.HOG0013890 Pxut g15679.t1 ninaG
N0.HOG0013890 Pmac g13150.t1 ninaG
N0.HOG0013890 Tsyl g13996.t1 ninaG
N0.HOG0013890 Pmal g3888.t1 ninaG
N0.HOG0013890 Lphl g11803.t1 ninaG
N0.HOG0013890 Lcor g5807.t1 ninaG
N0.HOG0013890 Cvir g27193.t1 ninaG
N0.HOG0013890 Cnem g2870.t1 ninaG
N0.HOG0013890 Hmel g7825.t1 ninaG
N0.HOG0013890 Dple g13800.t1 ninaG
N0.HOG0013890 Ccro g2212.t1 ninaG
N0.HOG0013890 Aips g22142.t1 ninaG
N0.HOG0018375 Pint g13857.t1 ninaG
N0.HOG0018375 Msex g5350.t1 Msex g5350
N0.HOG0018375 Pmal g3891.t2 ninaG
N0.HOG0018375 Hmel g7827.t1 ninaG
N0.HOG0018375 Ccro g2208.t1 ninaG
N0.HOG0005449 Bbet g4904.t1 Dple g13802
N0.HOG0005449 Cvir g402.t1 Dple g13802
N0.HOG0005449 Cnem g2873.t1 Dple g13802
N0.HOG0005449 Lphl g11805.t1 Dple g13802
N0.HOG0005449 Lcor g5810.t1 Dple g13802
N0.HOG0005449 Hmel g7828.t1 ninaG
N0.HOG0005449 Dple g13802.t1 Dple g13802
N0.HOG0005449 Tsyl g14001.t1 Dple g13802
N0.HOG0005449 Tsyl g13999.t1 Dple g13802
N0.HOG0005449 Pmal g3892.t1 Dple g13802
N0.HOG0005449 Ccro g2210.t1 Dple g13802
N0.HOG0005449 Ccro g2209.t1 Dple g13802
N0.HOG0005449 Ccro g2207.t1 Dple g13802
N0.HOG0005449 Pxut g15681.t1 Dple g13802
N0.HOG0005449 Pmac g13152.t1 Dple g13802
N0.HOG0005449 Bmor g19221.t1 Dple g13802
N0.HOG0005449 Zfil g969.t1 Dple g13802
N0.HOG0005449 Pint g13856.t1 Dple g13802
N0.HOG0005449 Bbet g15438.t1 Dple g13802
N0.HOG0005449 Msex g5351.t1 ninaG
N0.HOG0005449 Hpyr g11136.t1 Dple g13802
N0.HOG0005449 Sbem g12706.t1 Dple g13802
N0.HOG0005449 Aips g22139.t1 Dple g13802
N0.HOG0005449 Aips g22138.t1 Dple g13802
N0.HOG0008241 Tsyl g887.t1 Dple g16019
N0.HOG0008241 Pmal g9656.t1 Dple g16019
N0.HOG0008241 Mleu g5757.t1 Dple g16019
N0.HOG0008241 Pxut g2675.t1 Dple g16019
N0.HOG0008241 Pmac g3878.t1 Dple g16019
N0.HOG0008241 Lphl g2927.t1 Dple g16019
N0.HOG0008241 Lcor g16749.t1 Dple g16019
N0.HOG0008241 Cvir g223.t1 Dple g16019
N0.HOG0008241 Cnem g955.t1 Dple g16019
N0.HOG0008241 Hmel g4071.t1 ninaG
N0.HOG0008241 Dple g16019.t1 Dple g16019
N0.HOG0008241 Ccro g763.t1 Dple g16019
N0.HOG0008241 Sbem g6562.t1 Dple g16019
N0.HOG0008241 Pint g15020.t1 Dple g16019
N0.HOG0008241 Zfil g2580.t1 Dple g16019
N0.HOG0008241 Hpyr g14626.t1 Dple g16019
N0.HOG0008241 Hpyr g14625.t1 Dple g16019
N0.HOG0008241 Msex g1892.t1 Msex g1892
N0.HOG0008241 Bmor g14267.t1 Dple g16019
N0.HOG0008241 Bbet g8015.t1 Dple g16019
N0.HOG0008241 Aips g2341.t1 Dple g16019
N0.HOG0002702 Sbem g12708.t1 Dple g13803
N0.HOG0002702 Pmac g13153.t1 Dple g13803
N0.HOG0002702 Tsyl g14004.t1 Dple g13803
N0.HOG0002702 Tsyl g14003.t1 Dple g13803
N0.HOG0002702 Pmal g3894.t1 Dple g13803
N0.HOG0002702 Hmel g7829.t1 Hmel g7829
N0.HOG0002702 Dple g13803.t1 Dple g13803
N0.HOG0002702 Lcor g5811.t1 Dple g13803
N0.HOG0002702 Cvir g399.t1 Dple g13803
N0.HOG0002702 Cnem g4355.t1 Dple g13803
N0.HOG0002702 Cnem g4356.t1 Dple g13803
N0.HOG0002702 Cvir g401.t1 Dple g13803
N0.HOG0002702 Cnem g1693.t1 Dple g13803
N0.HOG0002702 Ccro g2206.t1 Dple g13803
N0.HOG0002702 Pint g13854.t1 Dple g13803
N0.HOG0002702 Msex g5354.t1 ninaG
N0.HOG0002702 Bmor g19218.t1 Dple g13803
N0.HOG0002702 Bmor g19217.t1 Dple g13803
N0.HOG0002702 Bmor g19219.t1 Dple g13803
N0.HOG0002702 Bbet g15436.t1 Dple g13803
N0.HOG0002702 Aips g22136.t1 Dple g13803
N0.HOG0002702 Sbem g12707.t1 Dple g13803
N0.HOG0002702 Pint g13855.t1 Dple g13803
N0.HOG0002702 Tsyl g14000.t1 Dple g13803
N0.HOG0002702 Tsyl g14002.t1 Dple g13803
N0.HOG0002702 Pmal g3893.t1 Dple g13803
N0.HOG0002702 Mleu g2469.t1 Dple g13803
N0.HOG0002702 Bbet g15437.t1 Dple g13803
N0.HOG0002702 Msex g5353.t1 ninaG
N0.HOG0002702 Msex g5352.t1 ninaG
N0.HOG0002702 Bmor g19220.t1 Dple g13803
N0.HOG0002702 Hpyr g11137.t1 Dple g13803
N0.HOG0002702 Aips g22137.t1 Dple g13803
N0.HOG0002701 Zfil g970.t1 ninaG
N0.HOG0002701 Bbet g15435.t1 ninaG
N0.HOG0002701 Tsyl g14005.t1 ninaG
N0.HOG0002701 Pmal g3895.t1 ninaG
N0.HOG0002701 Mleu g2468.t1 ninaG
N0.HOG0002701 Pxut g15682.t1 ninaG
N0.HOG0002701 Pmac g13154.t1 ninaG
N0.HOG0002701 Lphl g11806.t1 ninaG
N0.HOG0002701 Lcor g5812.t1 ninaG
N0.HOG0002701 Cvir g400.t1 ninaG
N0.HOG0002701 Cnem g1692.t1 ninaG
N0.HOG0002701 Hmel g7830.t1 Hmel g7831
N0.HOG0002701 Hmel g7831.t1 Hmel g7831
N0.HOG0002701 Dple g13804.t1 ninaG
N0.HOG0002701 Ccro g2205.t1 ninaG
N0.HOG0002701 Sbem g12709.t1 ninaG
N0.HOG0002701 Msex g5355.t1 ninaG
N0.HOG0002701 Bmor g19216.t1 ninaG
N0.HOG0002701 Hpyr g11138.t1 ninaG
N0.HOG0002701 Aips g22135.t1 ninaG
N0.HOG0011379 Tsyl g4204.t1 ninaG
N0.HOG0011379 Pmal g10219.t1 ninaG
N0.HOG0011379 Pxut g7383.t1 ninaG
N0.HOG0011379 Pmac g10164.t1 ninaG
N0.HOG0011379 Hmel g3782.t1 ninaG
N0.HOG0011379 Dple g2948.t1 ninaG
N0.HOG0011379 Cvir g21598.t1 ninaG
N0.HOG0011379 Cnem g9722.t1 ninaG
N0.HOG0011379 Lphl g12841.t1 ninaG
N0.HOG0011379 Lcor g14313.t1 ninaG
N0.HOG0011379 Ccro g1018.t1 ninaG
N0.HOG0011379 Zfil g5131.t1 ninaG
N0.HOG0011379 Mleu g11038.t1 ninaG
N0.HOG0011379 Hpyr g927.t1 ninaG
N0.HOG0011379 Sbem g1495.t1 ninaG
N0.HOG0011379 Aips g5350.t1 ninaG
N0.HOG0011379 Msex g3978.t1 ninaG
N0.HOG0011379 Bmor g8113.t1 ninaG
N0.HOG0011379 Pint g10492.t1 ninaG
N0.HOG0011379 Bbet g12774.t1 ninaG

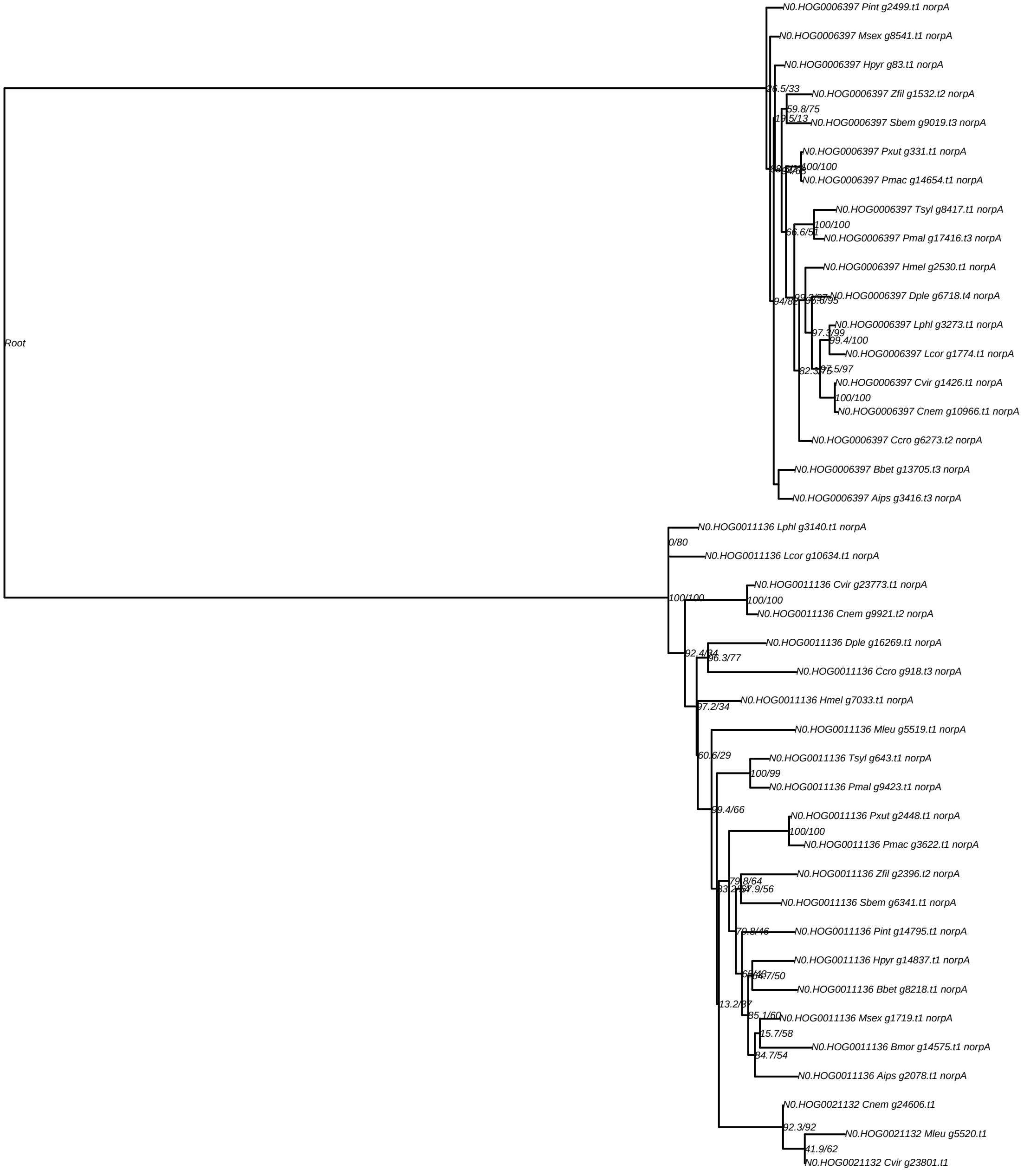

Root
N0.HOG0006397 Pint g2499.t1 norpA
N0.HOG0006397 Msex g8541.t1 norpA
N0.HOG0006397 Hpyr g83.t1 norpA
26.5/33
N0.HOG0006397 Zfil g1532.t2 norpA
59.8/75
N0.HOG0006397 Sbem g9019.t3 norpA
N0.HOG0006397 Pxut g331.t1 norpA
100/100
N0.HOG0006397 Pmac g14654.t1 norpA
N0.HOG0006397 Tsyl g8417.t1 norpA
100/100
N0.HOG0006397 Pmal g17416.t3 norpA
N0.HOG0006397 Hmel g2530.t1 norpA
N0.HOG0006397 Dple g6718.t4 norpA
94/89
N0.HOG0006397 Lphl g3273.t1 norpA
97.3/99
99.4/100
N0.HOG0006397 Lcor g1774.t1 norpA
97.5/97
N0.HOG0006397 Cvir g1426.t1 norpA
100/100
N0.HOG0006397 Cnem g10966.t1 norpA
N0.HOG0006397 Ccro g6273.t2 norpA
N0.HOG0006397 Bbet g13705.t3 norpA
N0.HOG0006397 Aips g3416.t3 norpA
N0.HOG0011136 Lphl g3140.t1 norpA
0/80
N0.HOG0011136 Lcor g10634.t1 norpA
N0.HOG0011136 Cvir g23773.t1 norpA
100/100
100/100
N0.HOG0011136 Cnem g9921.t2 norpA
N0.HOG0011136 Dple g16269.t1 norpA
96.3/77
N0.HOG0011136 Ccro g918.t3 norpA
N0.HOG0011136 Hmel g7033.t1 norpA
97.2/34
N0.HOG0011136 Mleu g5519.t1 norpA
60.6/29
N0.HOG0011136 Tsyl g643.t1 norpA
100/99
N0.HOG0011136 Pmal g9423.t1 norpA
99.4/66
N0.HOG0011136 Pxut g2448.t1 norpA
100/100
N0.HOG0011136 Pmac g3622.t1 norpA
N0.HOG0011136 Zfil g2396.t2 norpA
N0.HOG0011136 Sbem g6341.t1 norpA
N0.HOG0011136 Pint g14795.t1 norpA
N0.HOG0011136 Hpyr g14837.t1 norpA
N0.HOG0011136 Bbet g8218.t1 norpA
13.2/37
85.1/60
N0.HOG0011136 Msex g1719.t1 norpA
15.7/58
N0.HOG0011136 Bmor g14575.t1 norpA
84.7/54
N0.HOG0011136 Aips g2078.t1 norpA
N0.HOG0021132 Cnem g24606.t1
92.3/92
N0.HOG0021132 Mleu g5520.t1
41.9/62
N0.HOG0021132 Cvir g23801.t1

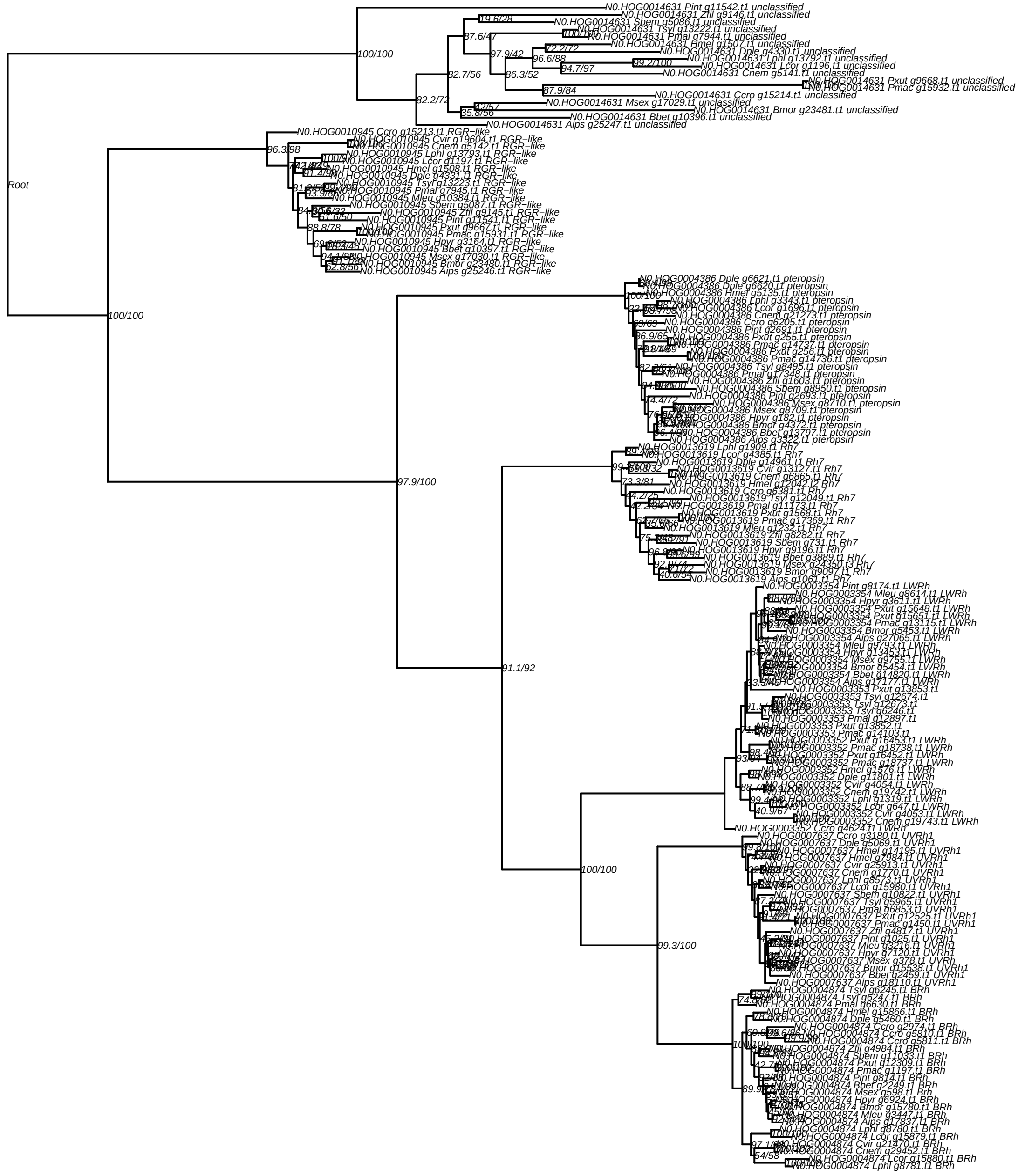
Root
100/100
N0.HOG0014631 Pint g11542.t1 unclassified
N0.HOG0014631 Zfil g9146.t1 unclassified
N0.HOG0014631 Sbem g5086.t1 unclassified
N0.HOG0014631 Tsyl g13222.t1 unclassified
N0.HOG0014631 Pmal g7944.t1 unclassified
N0.HOG0014631 Hmel g1507.t1 unclassified
N0.HOG0014631 Dple g4330.t1 unclassified
N0.HOG0014631 Lphl g13792.t1 unclassified
N0.HOG0014631 Lcor g1196.t1 unclassified
N0.HOG0014631 Cnem g5141.t1 unclassified
N0.HOG0014631 Pxut g9668.t1 unclassified
N0.HOG0014631 Pmac g15932.t1 unclassified
N0.HOG0014631 Ccro g15214.t1 unclassified
N0.HOG0014631 Msex g17029.t1 unclassified
N0.HOG0014631 Bmor g23481.t1 unclassified
N0.HOG0014631 Bbet g10396.t1 unclassified
N0.HOG0014631 Aips g25247.t1 unclassified
N0.HOG0010945 Ccro g15213.t1 RGR−like
N0.HOG0010945 Cvir g19604.t1 RGR−like
N0.HOG0010945 Cnem g5142.t1 RGR−like
N0.HOG0010945 Lphl g13793.t1 RGR−like
N0.HOG0010945 Lcor g1197.t1 RGR−like
N0.HOG0010945 Hmel g1508.t1 RGR−like
N0.HOG0010945 Dple g4331.t1 RGR−like
N0.HOG0010945 Tsyl g13223.t1 RGR−like
N0.HOG0010945 Pmal g7945.t1 RGR−like
N0.HOG0010945 Mleu g10384.t1 RGR−like
N0.HOG0010945 Sbem g5087.t1 RGR−like
N0.HOG0010945 Zfil g9145.t1 RGR−like
N0.HOG0010945 Pint g11541.t1 RGR−like
N0.HOG0010945 Pxut g9667.t1 RGR−like
N0.HOG0010945 Pmac g15931.t1 RGR−like
N0.HOG0010945 Hpyr g3164.t1 RGR−like
N0.HOG0010945 Bbet g10397.t1 RGR−like
N0.HOG0010945 Msex g17030.t1 RGR−like
N0.HOG0010945 Bmor g23480.t1 RGR−like
N0.HOG0010945 Aips g25246.t1 RGR−like
N0.HOG0004386 Dple g6621.t1 pteropsin
N0.HOG0004386 Dple g6620.t1 pteropsin
N0.HOG0004386 Hmel g5135.t1 pteropsin
N0.HOG0004386 Lphl g3343.t1 pteropsin
N0.HOG0004386 Lcor g1696.t1 pteropsin
N0.HOG0004386 Cnem g21273.t1 pteropsin
N0.HOG0004386 Ccro g6205.t1 pteropsin
N0.HOG0004386 Pint g2691.t1 pteropsin
N0.HOG0004386 Pxut g255.t1 pteropsin
N0.HOG0004386 Pmac g14737.t1 pteropsin
N0.HOG0004386 Pxut g256.t1 pteropsin
N0.HOG0004386 Pmac g14736.t1 pteropsin
N0.HOG0004386 Tsyl g8495.t1 pteropsin
N0.HOG0004386 Pmal g17348.t1 pteropsin
N0.HOG0004386 Zfil g1603.t1 pteropsin
N0.HOG0004386 Sbem g8950.t1 pteropsin
N0.HOG0004386 Pint g2693.t1 pteropsin
N0.HOG0004386 Msex g8710.t1 pteropsin
N0.HOG0004386 Msex g8709.t1 pteropsin
N0.HOG0004386 Hpyr g182.t1 pteropsin
N0.HOG0004386 Bmor g4372.t1 pteropsin
N0.HOG0004386 Bbet g13797.t1 pteropsin
N0.HOG0004386 Aips g3322.t1 pteropsin
97.9/100
N0.HOG0013619 Lphl g1909.t1 Rh7
N0.HOG0013619 Lcor g4385.t1 Rh7
N0.HOG0013619 Dple g14961.t1 Rh7
N0.HOG0013619 Cvir g13127.t1 Rh7
N0.HOG0013619 Cnem g6865.t1 Rh7
N0.HOG0013619 Hmel g12042.t2 Rh7
N0.HOG0013619 Ccro g6381.t1 Rh7
N0.HOG0013619 Tsyl g12049.t1 Rh7
N0.HOG0013619 Pmal g11173.t1 Rh7
N0.HOG0013619 Pxut g1568.t1 Rh7
N0.HOG0013619 Pmac g17369.t1 Rh7
N0.HOG0013619 Mleu g1232.t1 Rh7
N0.HOG0013619 Zfil g8282.t1 Rh7
N0.HOG0013619 Sbem g731.t1 Rh7
N0.HOG0013619 Hpyr g9196.t1 Rh7
N0.HOG0013619 Bbet g3889.t1 Rh7
N0.HOG0013619 Msex g24350.t3 Rh7
N0.HOG0013619 Bmor g9097.t1 Rh7
N0.HOG0013619 Aips g1061.t1 Rh7
91.1/92
N0.HOG0003354 Pint g8174.t1 LWRh
N0.HOG0003354 Mleu g8614.t1 LWRh
N0.HOG0003354 Hpyr g3611.t1 LWRh
N0.HOG0003354 Pxut g15648.t1 LWRh
N0.HOG0003354 Pxut g15651.t1 LWRh
N0.HOG0003354 Pmac g13115.t1 LWRh
N0.HOG0003354 Bmor g5453.t1 LWRh
N0.HOG0003354 Aips g27065.t1 LWRh
N0.HOG0003354 Mleu g9793.t1 LWRh
N0.HOG0003354 Hpyr g13453.t1 LWRh
N0.HOG0003354 Msex g9755.t1 LWRh
N0.HOG0003354 Bmor g5454.t1 LWRh
N0.HOG0003354 Bbet g14820.t1 LWRh
N0.HOG0003354 Aips g17177.t1 LWRh
N0.HOG0003353 Pxut g13853.t1
N0.HOG0003353 Tsyl g12674.t1
N0.HOG0003353 Tsyl g12673.t1
N0.HOG0003353 Tsyl g6246.t1
N0.HOG0003353 Pmal g12897.t1
N0.HOG0003353 Pxut g13852.t1
N0.HOG0003353 Pmac g14103.t1
N0.HOG0003352 Pxut g16453.t1 LWRh
N0.HOG0003352 Pmac g18738.t1 LWRh
N0.HOG0003352 Pxut g16452.t1 LWRh
N0.HOG0003352 Pmac g18737.t1 LWRh
N0.HOG0003352 Hmel g1576.t1 LWRh
N0.HOG0003352 Dple g11801.t1 LWRh
N0.HOG0003352 Cvir g4054.t1 LWRh
N0.HOG0003352 Cnem g19742.t1 LWRh
N0.HOG0003352 Lphl g1319.t1 LWRh
N0.HOG0003352 Lcor g647.t1 LWRh
N0.HOG0003352 Cvir g4053.t1 LWRh
N0.HOG0003352 Cnem g19743.t1 LWRh
N0.HOG0003352 Ccro g4624.t1 LWRh
100/100
N0.HOG0007637 Ccro g3180.t1 UVRh1
N0.HOG0007637 Dple g5069.t1 UVRh1
N0.HOG0007637 Hmel g14195.t1 UVRh1
N0.HOG0007637 Hmel g7984.t1 UVRh1
N0.HOG0007637 Cvir g25913.t1 UVRh1
N0.HOG0007637 Cnem g1770.t1 UVRh1
N0.HOG0007637 Lphl g8573.t1 UVRh1
N0.HOG0007637 Lcor g15980.t1 UVRh1
N0.HOG0007637 Sbem g10822.t1 UVRh1
N0.HOG0007637 Tsyl g5965.t1 UVRh1
N0.HOG0007637 Pmal g6853.t1 UVRh1
N0.HOG0007637 Pxut g12525.t1 UVRh1
N0.HOG0007637 Pmac g1450.t1 UVRh1
N0.HOG0007637 Zfil g4817.t1 UVRh1
N0.HOG0007637 Pint g1025.t1 UVRh1
N0.HOG0007637 Mleu g3216.t1 UVRh1
N0.HOG0007637 Hpyr g7120.t1 UVRh1
N0.HOG0007637 Msex g378.t1 UVRh1
N0.HOG0007637 Bmor g15538.t1 UVRh1
N0.HOG0007637 Bbet g2459.t1 UVRh1
N0.HOG0007637 Aips g18110.t1 UVRh1
99.3/100
N0.HOG0004874 Tsyl g6245.t1 BRh
N0.HOG0004874 Tsyl g6247.t1 BRh
N0.HOG0004874 Pmal g6630.t1 BRh
N0.HOG0004874 Hmel g15866.t1 BRh
N0.HOG0004874 Dple g5460.t1 BRh
N0.HOG0004874 Ccro g2974.t1 BRh
N0.HOG0004874 Ccro g5810.t1 BRh
N0.HOG0004874 Ccro g5811.t1 BRh
N0.HOG0004874 Zfil g4984.t1 BRh
N0.HOG0004874 Sbem g11033.t1 BRh
N0.HOG0004874 Pxut g12309.t1 BRh
N0.HOG0004874 Pmac g1197.t1 BRh
N0.HOG0004874 Pint g814.t1 BRh
N0.HOG0004874 Bbet g2249.t1 BRh
N0.HOG0004874 Msex g598.t1 BRh
N0.HOG0004874 Hpyr g6924.t1 BRh
N0.HOG0004874 Bmor g15780.t1 BRh
N0.HOG0004874 Mleu g3447.t1 BRh
N0.HOG0004874 Aips g17837.t1 BRh
N0.HOG0004874 Lphl g8780.t1 BRh
N0.HOG0004874 Lcor g15879.t1 BRh
N0.HOG0004874 Cvir g21470.t1 BRh
N0.HOG0004874 Cnem g29452.t1 BRh
N0.HOG0004874 Lcor g15880.t1 BRh
N0.HOG0004874 Lphl g8781.t1 BRh

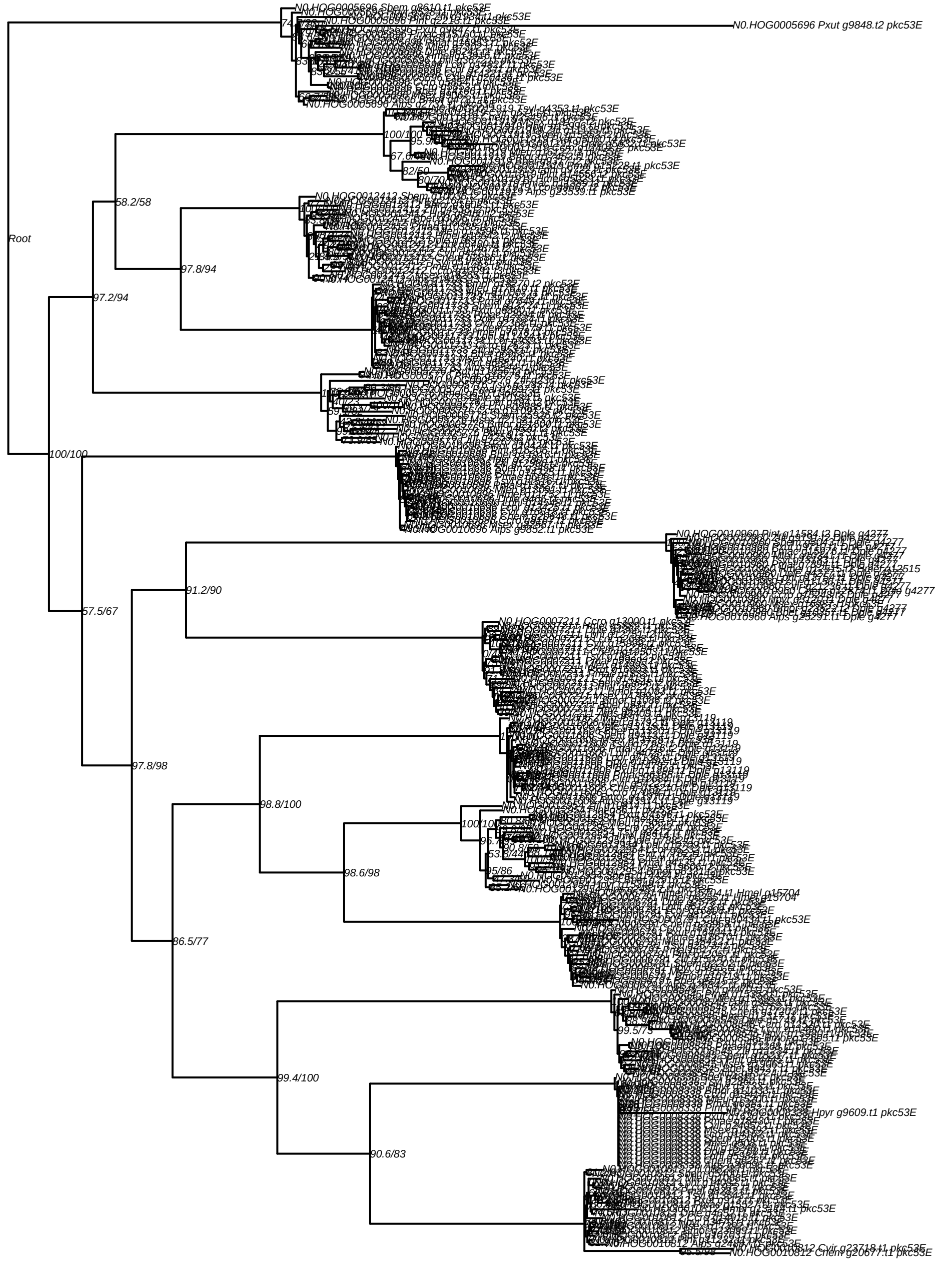

Root
N0.HOG0005696 Sbem g8610.t1 pkc53E
N0.HOG0005696 Pxut g9848.t2 pkc53E
N0.HOG0011919 Tsyl g4353.t1 pkc53E
100/100
58.2/58
97.8/94
97.2/94
100/100
57.5/67
91.2/90
97.8/98
86.5/77
98.8/100
98.6/98
99.4/100
90.6/83
N0.HOG0010960 Pint g11584.t2 Dple g4277
N0.HOG0010960 Alps g25291.t1 Dple g4277
N0.HOG0007211 Ccro g13000.t1 pkc53E
N0.HOG0010696 Alps g9852.t1 pkc53E
N0.HOG0008338 Hpyr g9609.t1 pkc53E
N0.HOG0010812 Cvir g23718.t1 pkc53E
N0.HOG0010812 Chem g20677.t1 pkc53E

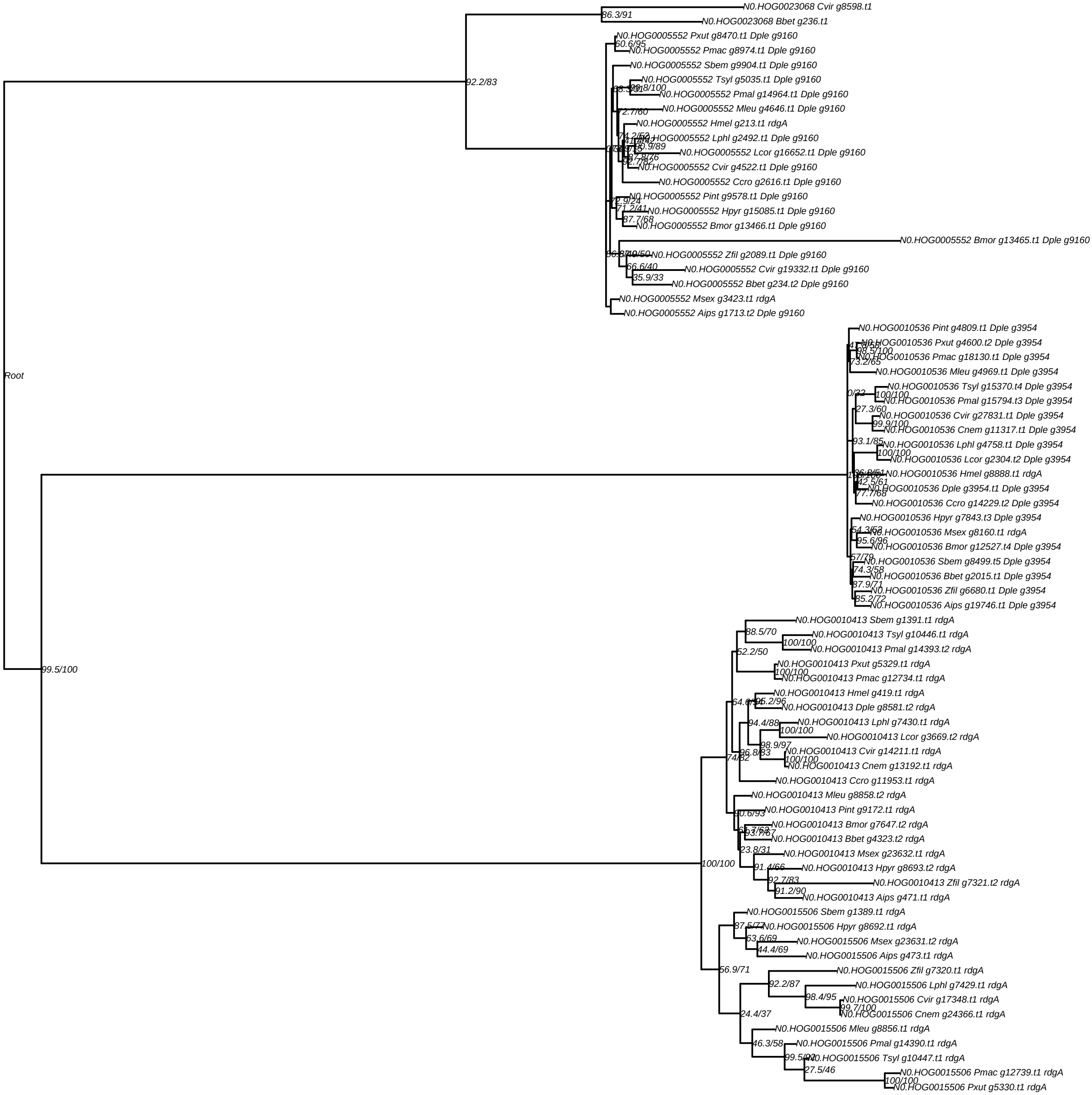
N0.HOG0023068 Cvir g8598.t1
86.3/91
N0.HOG0023068 Bbet g236.t1
N0.HOG0005552 Pxut g8470.t1 Dple g9160
60.6/95
N0.HOG0005552 Pmac g8974.t1 Dple g9160
N0.HOG0005552 Sbem g9904.t1 Dple g9160
N0.HOG0005552 Tsyl g5035.t1 Dple g9160
92.2/83
N0.HOG0005552 Pmal g14964.t1 Dple g9160
72.7/60
N0.HOG0005552 Mleu g4646.t1 Dple g9160
N0.HOG0005552 Hmel g213.t1 rdgA
N0.HOG0005552 Lphl g2492.t1 Dple g9160
N0.HOG0005552 Lcor g16652.t1 Dple g9160
N0.HOG0005552 Cvir g4522.t1 Dple g9160
N0.HOG0005552 Ccro g2616.t1 Dple g9160
N0.HOG0005552 Pint g9578.t1 Dple g9160
N0.HOG0005552 Hpyr g15085.t1 Dple g9160
87.7/68
N0.HOG0005552 Bmor g13466.t1 Dple g9160
N0.HOG0005552 Bmor g13465.t1 Dple g9160
N0.HOG0005552 Zfil g2089.t1 Dple g9160
66.6/40
N0.HOG0005552 Cvir g19332.t1 Dple g9160
35.9/33
N0.HOG0005552 Bbet g234.t2 Dple g9160
N0.HOG0005552 Msex g3423.t1 rdgA
N0.HOG0005552 Aips g1713.t2 Dple g9160
N0.HOG0010536 Pint g4809.t1 Dple g3954
N0.HOG0010536 Pxut g4600.t2 Dple g3954
N0.HOG0010536 Pmac g18130.t1 Dple g3954
73.2/65
N0.HOG0010536 Mleu g4969.t1 Dple g3954
Root
N0.HOG0010536 Tsyl g15370.t4 Dple g3954
0/32
100/100
N0.HOG0010536 Pmal g15794.t3 Dple g3954
27.3/60
N0.HOG0010536 Cvir g27831.t1 Dple g3954
99.9/100
N0.HOG0010536 Cnem g11317.t1 Dple g3954
93.1/85
N0.HOG0010536 Lphl g4758.t1 Dple g3954
100/100
N0.HOG0010536 Lcor g2304.t2 Dple g3954
N0.HOG0010536 Hmel g8888.t1 rdgA
42.5/61
N0.HOG0010536 Dple g3954.t1 Dple g3954
77.7/68
N0.HOG0010536 Ccro g14229.t2 Dple g3954
N0.HOG0010536 Hpyr g7843.t3 Dple g3954
N0.HOG0010536 Msex g8160.t1 rdgA
95.6/96
N0.HOG0010536 Bmor g12527.t4 Dple g3954
57/79
N0.HOG0010536 Sbem g8499.t5 Dple g3954
74.3/58
N0.HOG0010536 Bbet g2015.t1 Dple g3954
87.9/71
N0.HOG0010536 Zfil g6680.t1 Dple g3954
85.2/72
N0.HOG0010536 Aips g19746.t1 Dple g3954
N0.HOG0010413 Sbem g1391.t1 rdgA
88.5/70
N0.HOG0010413 Tsyl g10446.t1 rdgA
100/100
N0.HOG0010413 Pmal g14393.t2 rdgA
52.2/50
99.5/100
N0.HOG0010413 Pxut g5329.t1 rdgA
100/100
N0.HOG0010413 Pmac g12734.t1 rdgA
N0.HOG0010413 Hmel g419.t1 rdgA
N0.HOG0010413 Dple g8581.t2 rdgA
94.4/88
N0.HOG0010413 Lphl g7430.t1 rdgA
100/100
N0.HOG0010413 Lcor g3669.t2 rdgA
98.9/97
N0.HOG0010413 Cvir g14211.t1 rdgA
96.8/83
100/100
74/82
N0.HOG0010413 Cnem g13192.t1 rdgA
N0.HOG0010413 Ccro g11953.t1 rdgA
N0.HOG0010413 Mleu g8858.t2 rdgA
90.6/93
N0.HOG0010413 Pint g9172.t1 rdgA
N0.HOG0010413 Bmor g7647.t2 rdgA
N0.HOG0010413 Bbet g4323.t2 rdgA
23.8/31
N0.HOG0010413 Msex g23632.t1 rdgA
100/100
91.4/66
N0.HOG0010413 Hpyr g8693.t2 rdgA
92.7/83
N0.HOG0010413 Zfil g7321.t2 rdgA
91.2/90
N0.HOG0010413 Aips g471.t1 rdgA
N0.HOG0015506 Sbem g1389.t1 rdgA
N0.HOG0015506 Hpyr g8692.t1 rdgA
63.6/69
N0.HOG0015506 Msex g23631.t2 rdgA
44.4/69
N0.HOG0015506 Aips g473.t1 rdgA
56.9/71
N0.HOG0015506 Zfil g7320.t1 rdgA
92.2/87
N0.HOG0015506 Lphl g7429.t1 rdgA
98.4/95
N0.HOG0015506 Cvir g17348.t1 rdgA
99.7/100
24.4/37
N0.HOG0015506 Cnem g24366.t1 rdgA
N0.HOG0015506 Mleu g8856.t1 rdgA
46.3/58
N0.HOG0015506 Pmal g14390.t1 rdgA
N0.HOG0015506 Tsyl g10447.t1 rdgA
27.5/46
N0.HOG0015506 Pmac g12739.t1 rdgA
100/100
N0.HOG0015506 Pxut g5330.t1 rdgA

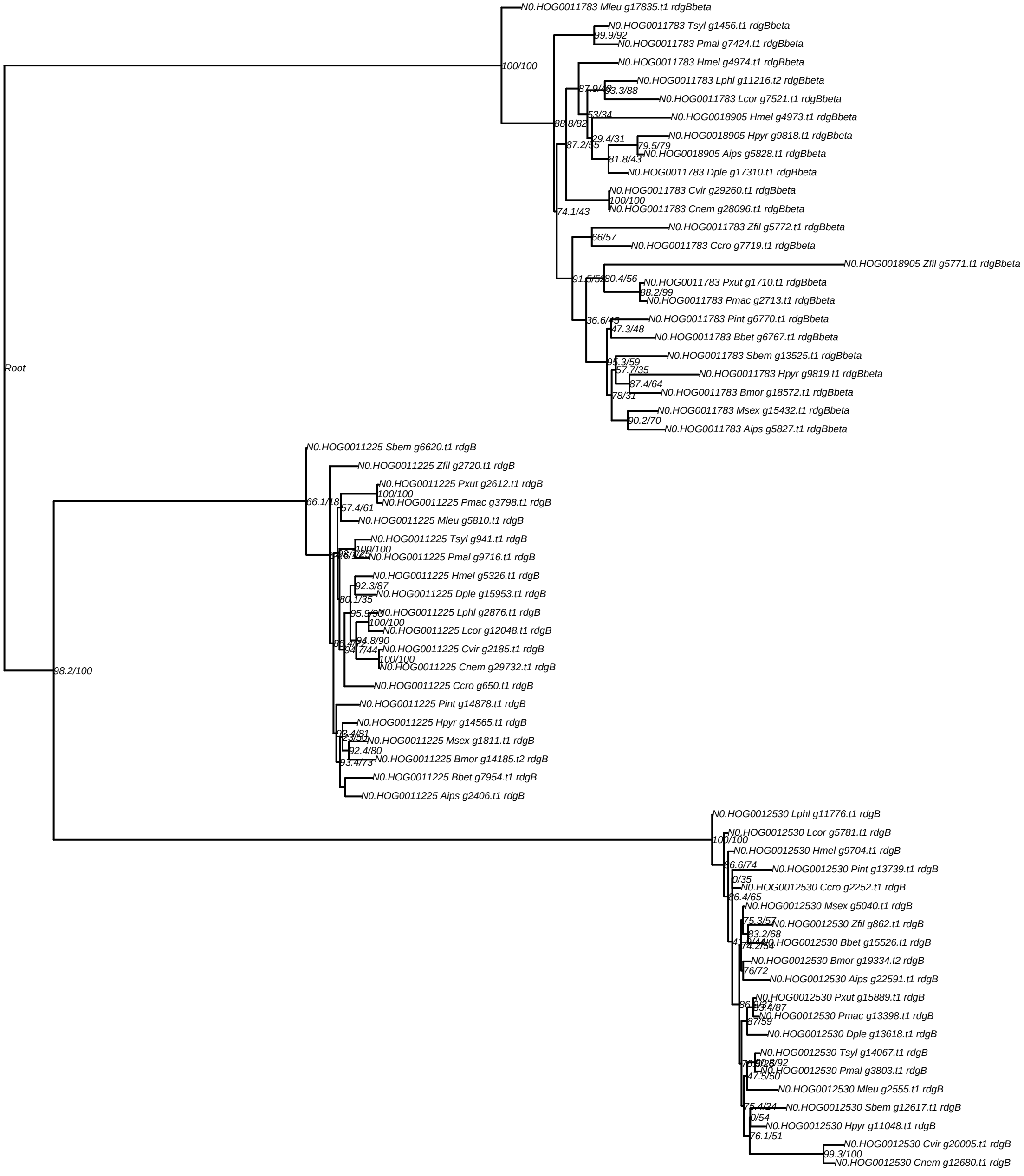
Root
100/100
98.2/100
N0.HOG0011783 Mleu g17835.t1 rdgBbeta
N0.HOG0011783 Tsyl g1456.t1 rdgBbeta
N0.HOG0011783 Pmal g7424.t1 rdgBbeta
N0.HOG0011783 Hmel g4974.t1 rdgBbeta
N0.HOG0011783 Lphl g11216.t2 rdgBbeta
N0.HOG0011783 Lcor g7521.t1 rdgBbeta
N0.HOG0018905 Hmel g4973.t1 rdgBbeta
N0.HOG0018905 Hpyr g9818.t1 rdgBbeta
N0.HOG0018905 Aips g5828.t1 rdgBbeta
N0.HOG0011783 Dple g17310.t1 rdgBbeta
N0.HOG0011783 Cvir g29260.t1 rdgBbeta
N0.HOG0011783 Cnem g28096.t1 rdgBbeta
N0.HOG0011783 Zfil g5772.t1 rdgBbeta
N0.HOG0011783 Ccro g7719.t1 rdgBbeta
N0.HOG0018905 Zfil g5771.t1 rdgBbeta
N0.HOG0011783 Pxut g1710.t1 rdgBbeta
N0.HOG0011783 Pmac g2713.t1 rdgBbeta
N0.HOG0011783 Pint g6770.t1 rdgBbeta
N0.HOG0011783 Bbet g6767.t1 rdgBbeta
N0.HOG0011783 Sbem g13525.t1 rdgBbeta
N0.HOG0011783 Hpyr g9819.t1 rdgBbeta
N0.HOG0011783 Bmor g18572.t1 rdgBbeta
N0.HOG0011783 Msex g15432.t1 rdgBbeta
N0.HOG0011783 Aips g5827.t1 rdgBbeta
N0.HOG0011225 Sbem g6620.t1 rdgB
N0.HOG0011225 Zfil g2720.t1 rdgB
N0.HOG0011225 Pxut g2612.t1 rdgB
N0.HOG0011225 Pmac g3798.t1 rdgB
N0.HOG0011225 Mleu g5810.t1 rdgB
N0.HOG0011225 Tsyl g941.t1 rdgB
N0.HOG0011225 Pmal g9716.t1 rdgB
N0.HOG0011225 Hmel g5326.t1 rdgB
N0.HOG0011225 Dple g15953.t1 rdgB
N0.HOG0011225 Lphl g2876.t1 rdgB
N0.HOG0011225 Lcor g12048.t1 rdgB
N0.HOG0011225 Cvir g2185.t1 rdgB
N0.HOG0011225 Cnem g29732.t1 rdgB
N0.HOG0011225 Ccro g650.t1 rdgB
N0.HOG0011225 Pint g14878.t1 rdgB
N0.HOG0011225 Hpyr g14565.t1 rdgB
N0.HOG0011225 Msex g1811.t1 rdgB
N0.HOG0011225 Bmor g14185.t2 rdgB
N0.HOG0011225 Bbet g7954.t1 rdgB
N0.HOG0011225 Aips g2406.t1 rdgB
N0.HOG0012530 Lphl g11776.t1 rdgB
N0.HOG0012530 Lcor g5781.t1 rdgB
N0.HOG0012530 Hmel g9704.t1 rdgB
N0.HOG0012530 Pint g13739.t1 rdgB
N0.HOG0012530 Ccro g2252.t1 rdgB
N0.HOG0012530 Msex g5040.t1 rdgB
N0.HOG0012530 Zfil g862.t1 rdgB
N0.HOG0012530 Bbet g15526.t1 rdgB
N0.HOG0012530 Bmor g19334.t2 rdgB
N0.HOG0012530 Aips g22591.t1 rdgB
N0.HOG0012530 Pxut g15889.t1 rdgB
N0.HOG0012530 Pmac g13398.t1 rdgB
N0.HOG0012530 Dple g13618.t1 rdgB
N0.HOG0012530 Tsyl g14067.t1 rdgB
N0.HOG0012530 Pmal g3803.t1 rdgB
N0.HOG0012530 Mleu g2555.t1 rdgB
N0.HOG0012530 Sbem g12617.t1 rdgB
N0.HOG0012530 Hpyr g11048.t1 rdgB
N0.HOG0012530 Cvir g20005.t1 rdgB
N0.HOG0012530 Cnem g12680.t1 rdgB

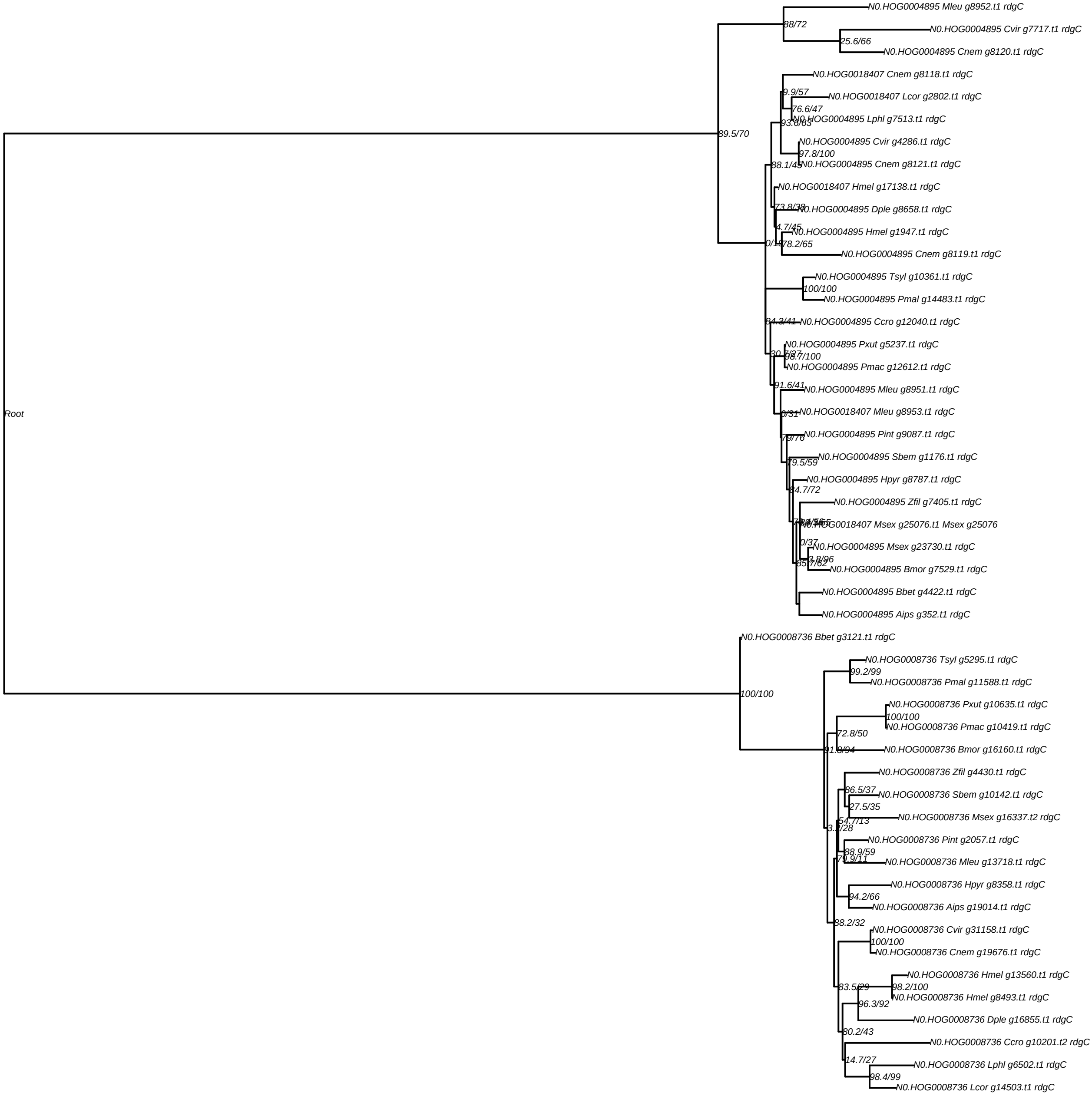

Root
89.5/70
88/72
N0.HOG0004895 Mleu g8952.t1 rdgC
25.6/66
N0.HOG0004895 Cvir g7717.t1 rdgC
N0.HOG0004895 Cnem g8120.t1 rdgC
N0.HOG0018407 Cnem g8118.t1 rdgC
9.9/57
N0.HOG0018407 Lcor g2802.t1 rdgC
76.6/47
N0.HOG0004895 Lphl g7513.t1 rdgC
N0.HOG0004895 Cvir g4286.t1 rdgC
97.8/100
N0.HOG0004895 Cnem g8121.t1 rdgC
N0.HOG0018407 Hmel g17138.t1 rdgC
N0.HOG0004895 Dple g8658.t1 rdgC
4.7/45
N0.HOG0004895 Hmel g1947.t1 rdgC
78.2/65
N0.HOG0004895 Cnem g8119.t1 rdgC
N0.HOG0004895 Tsyl g10361.t1 rdgC
100/100
N0.HOG0004895 Pmal g14483.t1 rdgC
N0.HOG0004895 Ccro g12040.t1 rdgC
N0.HOG0004895 Pxut g5237.t1 rdgC
N0.HOG0004895 Pmac g12612.t1 rdgC
91.6/41
N0.HOG0004895 Mleu g8951.t1 rdgC
N0.HOG0018407 Mleu g8953.t1 rdgC
N0.HOG0004895 Pint g9087.t1 rdgC
N0.HOG0004895 Sbem g1176.t1 rdgC
79.5/59
N0.HOG0004895 Hpyr g8787.t1 rdgC
N0.HOG0004895 Zfil g7405.t1 rdgC
N0.HOG0018407 Msex g25076.t1 Msex g25076
0/37
N0.HOG0004895 Msex g23730.t1 rdgC
N0.HOG0004895 Bmor g7529.t1 rdgC
N0.HOG0004895 Bbet g4422.t1 rdgC
N0.HOG0004895 Aips g352.t1 rdgC
100/100
N0.HOG0008736 Bbet g3121.t1 rdgC
N0.HOG0008736 Tsyl g5295.t1 rdgC
99.2/99
N0.HOG0008736 Pmal g11588.t1 rdgC
N0.HOG0008736 Pxut g10635.t1 rdgC
100/100
N0.HOG0008736 Pmac g10419.t1 rdgC
72.8/50
N0.HOG0008736 Bmor g16160.t1 rdgC
N0.HOG0008736 Zfil g4430.t1 rdgC
86.5/37
N0.HOG0008736 Sbem g10142.t1 rdgC
27.5/35
N0.HOG0008736 Msex g16337.t2 rdgC
54.7/13
N0.HOG0008736 Pint g2057.t1 rdgC
88.9/59
N0.HOG0008736 Mleu g13718.t1 rdgC
N0.HOG0008736 Hpyr g8358.t1 rdgC
94.2/66
N0.HOG0008736 Aips g19014.t1 rdgC
88.2/32
N0.HOG0008736 Cvir g31158.t1 rdgC
100/100
N0.HOG0008736 Cnem g19676.t1 rdgC
N0.HOG0008736 Hmel g13560.t1 rdgC
98.2/100
83.5/29
N0.HOG0008736 Hmel g8493.t1 rdgC
96.3/92
N0.HOG0008736 Dple g16855.t1 rdgC
80.2/43
N0.HOG0008736 Ccro g10201.t2 rdgC
14.7/27
N0.HOG0008736 Lphl g6502.t1 rdgC
98.4/99
N0.HOG0008736 Lcor g14503.t1 rdgC

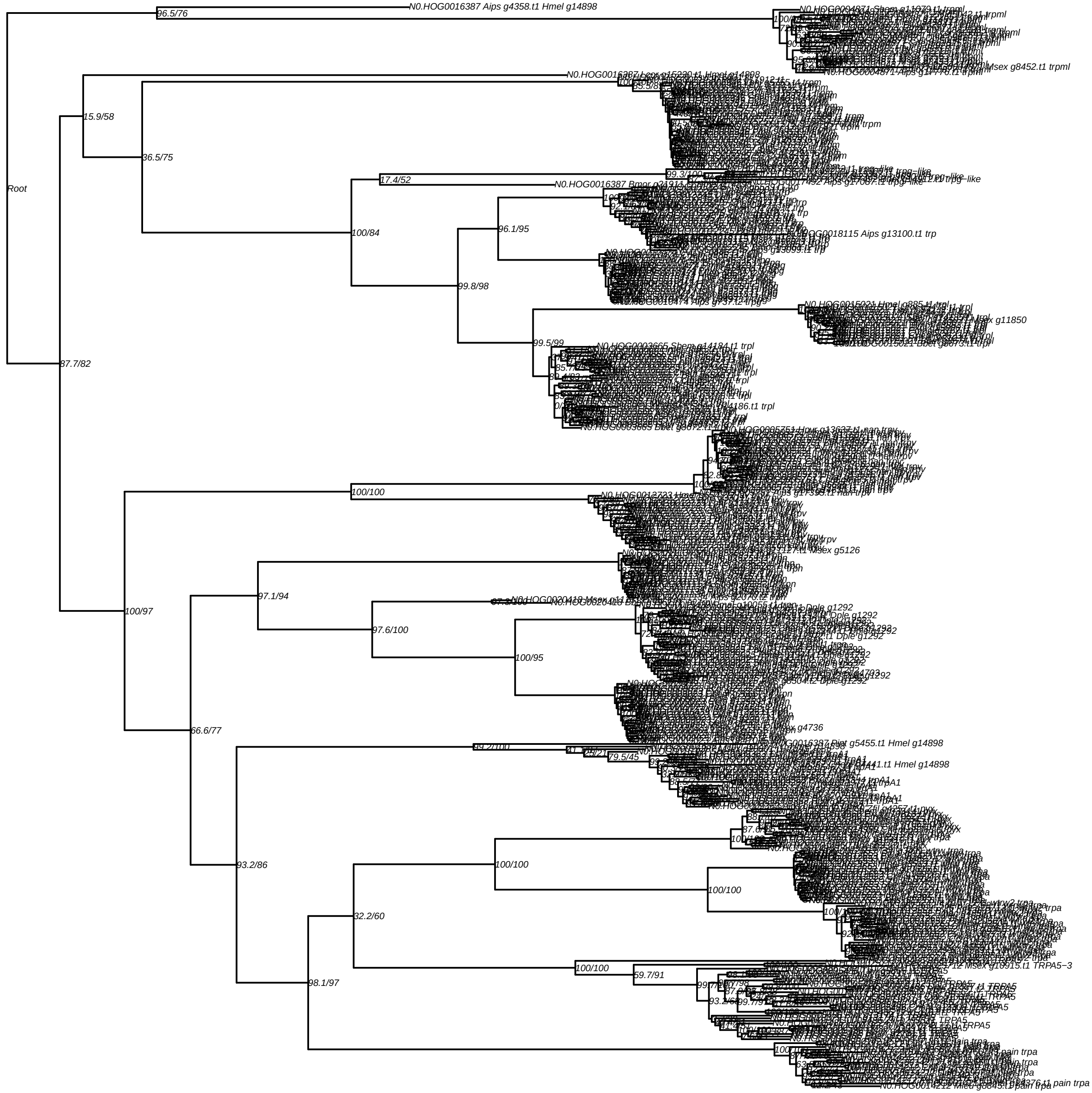
Root
N0.HOG0016387 Aips g4358.t1 Hmel g14898
96.5/76
15.9/58
36.5/75
87.7/82
17.4/52
99.3/100
100/84
96.1/95
N0.HOG0018115 Aips g13100.t1 trp
99.8/98
99.5/99
N0.HOG0003665 Shem g14184.t1 trpl
N0.HOG0003665 Bbet g8672.t1 trpl
N0.HOG0015021 Bbet g8673.t1 trpl
N0.HOG0004871 Aips g17776.t1 trpml
100/100
100/97
97.1/94
97.6/100
100/95
66.6/77
99.2/100
79.5/45
93.2/86
100/100
100/100
32.2/60
100/100
59.7/91
98.1/97
N0.HOG0016387 Pint g5455.t1 Hmel g14898
N0.HOG0014212 Mleu g6845.t1 pain trpa

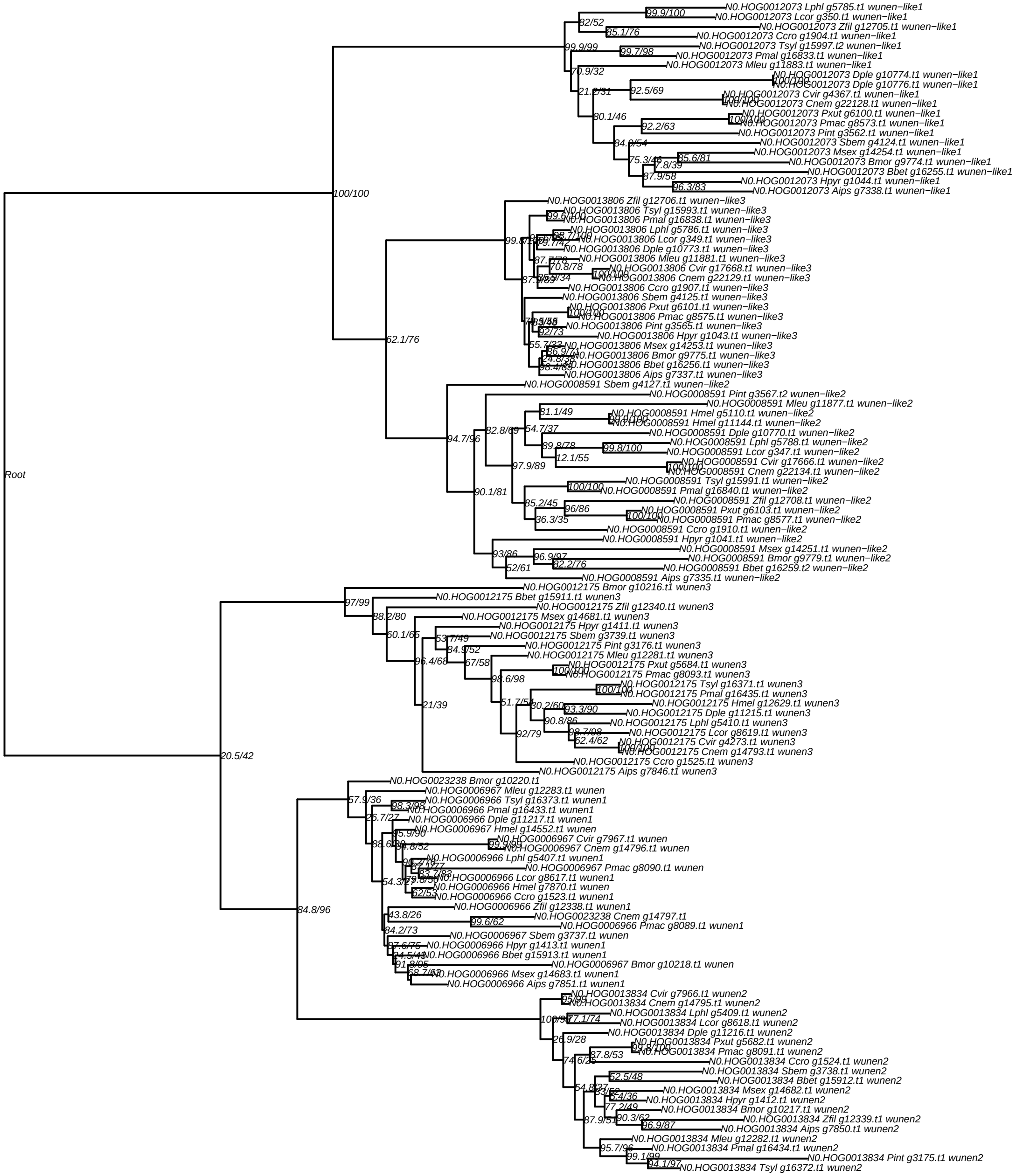
Root
N0.HOG0012073 Lphl g5785.t1 wunen−like1
N0.HOG0012073 Lcor g350.t1 wunen−like1
N0.HOG0012073 Zfil g12705.t1 wunen−like1
N0.HOG0012073 Ccro g1904.t1 wunen−like1
N0.HOG0012073 Tsyl g15997.t2 wunen−like1
N0.HOG0012073 Pmal g16833.t1 wunen−like1
N0.HOG0012073 Mleu g11883.t1 wunen−like1
N0.HOG0012073 Dple g10774.t1 wunen−like1
N0.HOG0012073 Dple g10776.t1 wunen−like1
N0.HOG0012073 Cvir g4367.t1 wunen−like1
N0.HOG0012073 Cnem g22128.t1 wunen−like1
N0.HOG0012073 Pxut g6100.t1 wunen−like1
N0.HOG0012073 Pmac g8573.t1 wunen−like1
N0.HOG0012073 Pint g3562.t1 wunen−like1
N0.HOG0012073 Sbem g4124.t1 wunen−like1
N0.HOG0012073 Msex g14254.t1 wunen−like1
N0.HOG0012073 Bmor g9774.t1 wunen−like1
N0.HOG0012073 Bbet g16255.t1 wunen−like1
N0.HOG0012073 Hpyr g1044.t1 wunen−like1
N0.HOG0012073 Aips g7338.t1 wunen−like1
N0.HOG0013806 Zfil g12706.t1 wunen−like3
N0.HOG0013806 Tsyl g15993.t1 wunen−like3
N0.HOG0013806 Pmal g16838.t1 wunen−like3
N0.HOG0013806 Lphl g5786.t1 wunen−like3
N0.HOG0013806 Lcor g349.t1 wunen−like3
N0.HOG0013806 Dple g10773.t1 wunen−like3
N0.HOG0013806 Mleu g11881.t1 wunen−like3
N0.HOG0013806 Cvir g17668.t1 wunen−like3
N0.HOG0013806 Cnem g22129.t1 wunen−like3
N0.HOG0013806 Ccro g1907.t1 wunen−like3
N0.HOG0013806 Sbem g4125.t1 wunen−like3
N0.HOG0013806 Pxut g6101.t1 wunen−like3
N0.HOG0013806 Pmac g8575.t1 wunen−like3
N0.HOG0013806 Pint g3565.t1 wunen−like3
N0.HOG0013806 Hpyr g1043.t1 wunen−like3
N0.HOG0013806 Msex g14253.t1 wunen−like3
N0.HOG0013806 Bmor g9775.t1 wunen−like3
N0.HOG0013806 Bbet g16256.t1 wunen−like3
N0.HOG0013806 Aips g7337.t1 wunen−like3
N0.HOG0008591 Sbem g4127.t1 wunen−like2
N0.HOG0008591 Pint g3567.t2 wunen−like2
N0.HOG0008591 Mleu g11877.t1 wunen−like2
N0.HOG0008591 Hmel g5110.t1 wunen−like2
N0.HOG0008591 Hmel g11144.t1 wunen−like2
N0.HOG0008591 Dple g10770.t1 wunen−like2
N0.HOG0008591 Lphl g5788.t1 wunen−like2
N0.HOG0008591 Lcor g347.t1 wunen−like2
N0.HOG0008591 Cvir g17666.t1 wunen−like2
N0.HOG0008591 Cnem g22134.t1 wunen−like2
N0.HOG0008591 Tsyl g15991.t1 wunen−like2
N0.HOG0008591 Pmal g16840.t1 wunen−like2
N0.HOG0008591 Zfil g12708.t1 wunen−like2
N0.HOG0008591 Pxut g6103.t1 wunen−like2
N0.HOG0008591 Pmac g8577.t1 wunen−like2
N0.HOG0008591 Ccro g1910.t1 wunen−like2
N0.HOG0008591 Hpyr g1041.t1 wunen−like2
N0.HOG0008591 Msex g14251.t1 wunen−like2
N0.HOG0008591 Bmor g9779.t1 wunen−like2
N0.HOG0008591 Bbet g16259.t2 wunen−like2
N0.HOG0008591 Aips g7335.t1 wunen−like2
N0.HOG0012175 Bmor g10216.t1 wunen3
N0.HOG0012175 Bbet g15911.t1 wunen3
N0.HOG0012175 Zfil g12340.t1 wunen3
N0.HOG0012175 Msex g14681.t1 wunen3
N0.HOG0012175 Hpyr g1411.t1 wunen3
N0.HOG0012175 Sbem g3739.t1 wunen3
N0.HOG0012175 Pint g3176.t1 wunen3
N0.HOG0012175 Mleu g12281.t1 wunen3
N0.HOG0012175 Pxut g5684.t1 wunen3
N0.HOG0012175 Pmac g8093.t1 wunen3
N0.HOG0012175 Tsyl g16371.t1 wunen3
N0.HOG0012175 Pmal g16435.t1 wunen3
N0.HOG0012175 Hmel g12629.t1 wunen3
N0.HOG0012175 Dple g11215.t1 wunen3
N0.HOG0012175 Lphl g5410.t1 wunen3
N0.HOG0012175 Lcor g8619.t1 wunen3
N0.HOG0012175 Cvir g4273.t1 wunen3
N0.HOG0012175 Cnem g14793.t1 wunen3
N0.HOG0012175 Ccro g1525.t1 wunen3
N0.HOG0012175 Aips g7846.t1 wunen3
N0.HOG0023238 Bmor g10220.t1
N0.HOG0006967 Mleu g12283.t1 wunen
N0.HOG0006966 Tsyl g16373.t1 wunen1
N0.HOG0006966 Pmal g16433.t1 wunen1
N0.HOG0006966 Dple g11217.t1 wunen1
N0.HOG0006967 Hmel g14552.t1 wunen
N0.HOG0006967 Cvir g7967.t1 wunen
N0.HOG0006967 Cnem g14796.t1 wunen
N0.HOG0006966 Lphl g5407.t1 wunen1
N0.HOG0006967 Pmac g8090.t1 wunen
N0.HOG0006966 Lcor g8617.t1 wunen1
N0.HOG0006966 Hmel g7870.t1 wunen1
N0.HOG0006966 Ccro g1523.t1 wunen1
N0.HOG0006966 Zfil g12338.t1 wunen1
N0.HOG0023238 Cnem g14797.t1
N0.HOG0006966 Pmac g8089.t1 wunen1
N0.HOG0006967 Sbem g3737.t1 wunen
N0.HOG0006966 Hpyr g1413.t1 wunen1
N0.HOG0006966 Bbet g15913.t1 wunen1
N0.HOG0006967 Bmor g10218.t1 wunen
N0.HOG0006966 Msex g14683.t1 wunen1
N0.HOG0006966 Aips g7851.t1 wunen1
N0.HOG0013834 Cvir g7966.t1 wunen2
N0.HOG0013834 Cnem g14795.t1 wunen2
N0.HOG0013834 Lphl g5409.t1 wunen2
N0.HOG0013834 Lcor g8618.t1 wunen2
N0.HOG0013834 Dple g11216.t1 wunen2
N0.HOG0013834 Pxut g5682.t1 wunen2
N0.HOG0013834 Pmac g8091.t1 wunen2
N0.HOG0013834 Ccro g1524.t1 wunen2
N0.HOG0013834 Sbem g3738.t1 wunen2
N0.HOG0013834 Bbet g15912.t1 wunen2
N0.HOG0013834 Msex g14682.t1 wunen2
N0.HOG0013834 Hpyr g1412.t1 wunen2
N0.HOG0013834 Bmor g10217.t1 wunen2
N0.HOG0013834 Zfil g12339.t1 wunen2
N0.HOG0013834 Aips g7850.t1 wunen2
N0.HOG0013834 Mleu g12282.t1 wunen2
N0.HOG0013834 Pmal g16434.t1 wunen2
N0.HOG0013834 Pint g3175.t1 wunen2
N0.HOG0013834 Tsyl g16372.t1 wunen2
