## Supplementary figures and images for "New genome reveals molecular signatures of adaptation to nocturnality in moth-like butterflies (Hedylidae)"

### Supplementary Data 11: Opsin gene trees

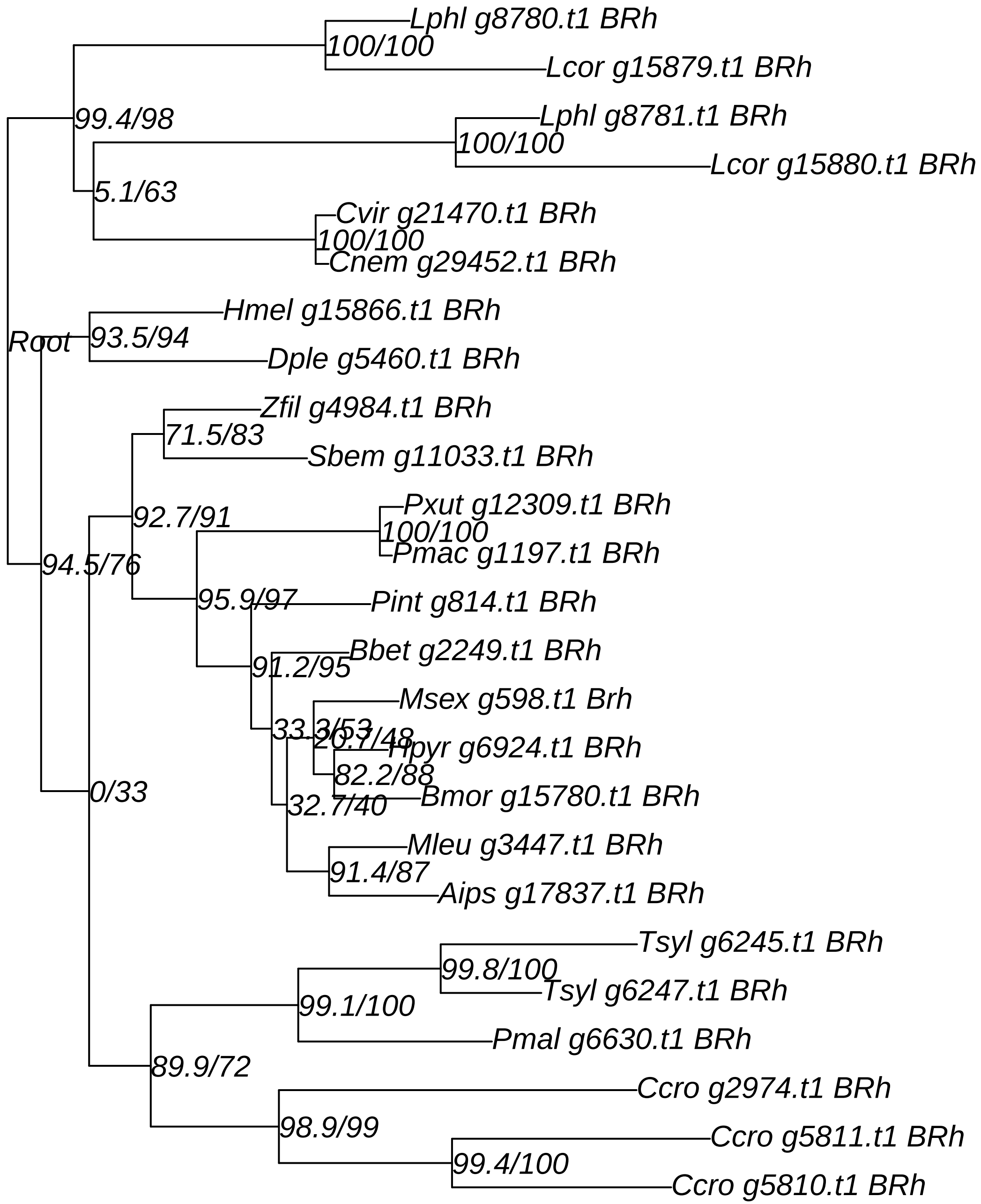

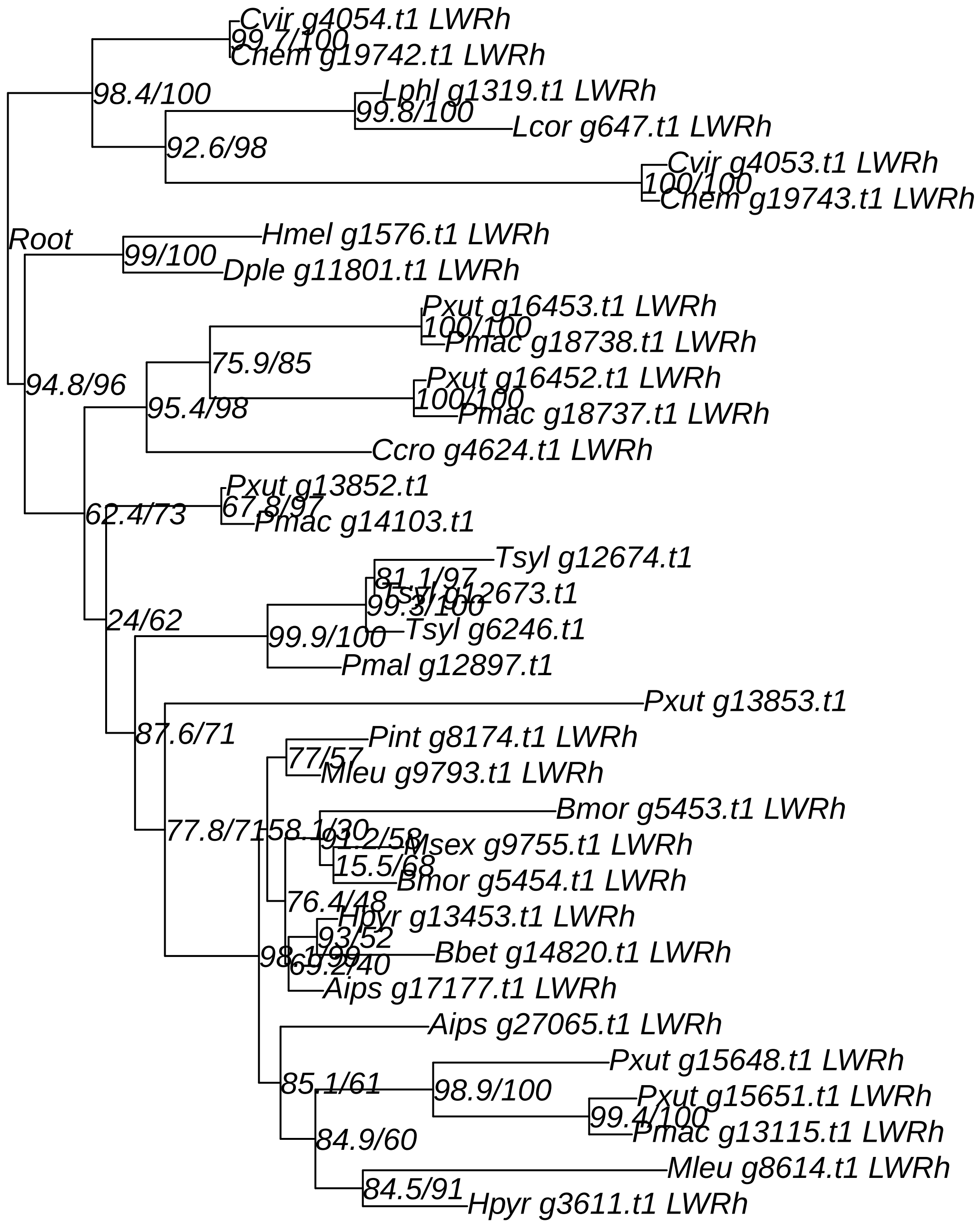

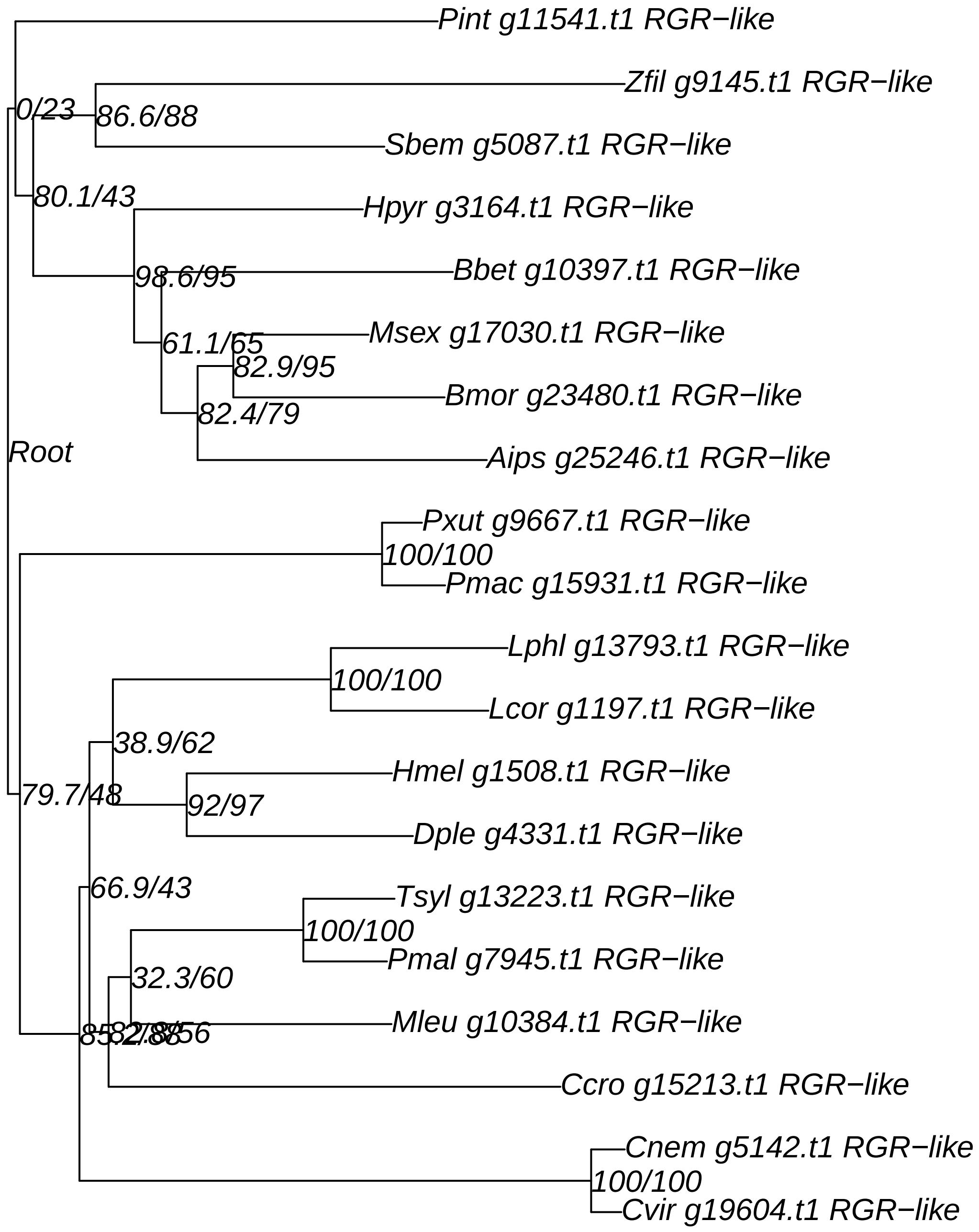

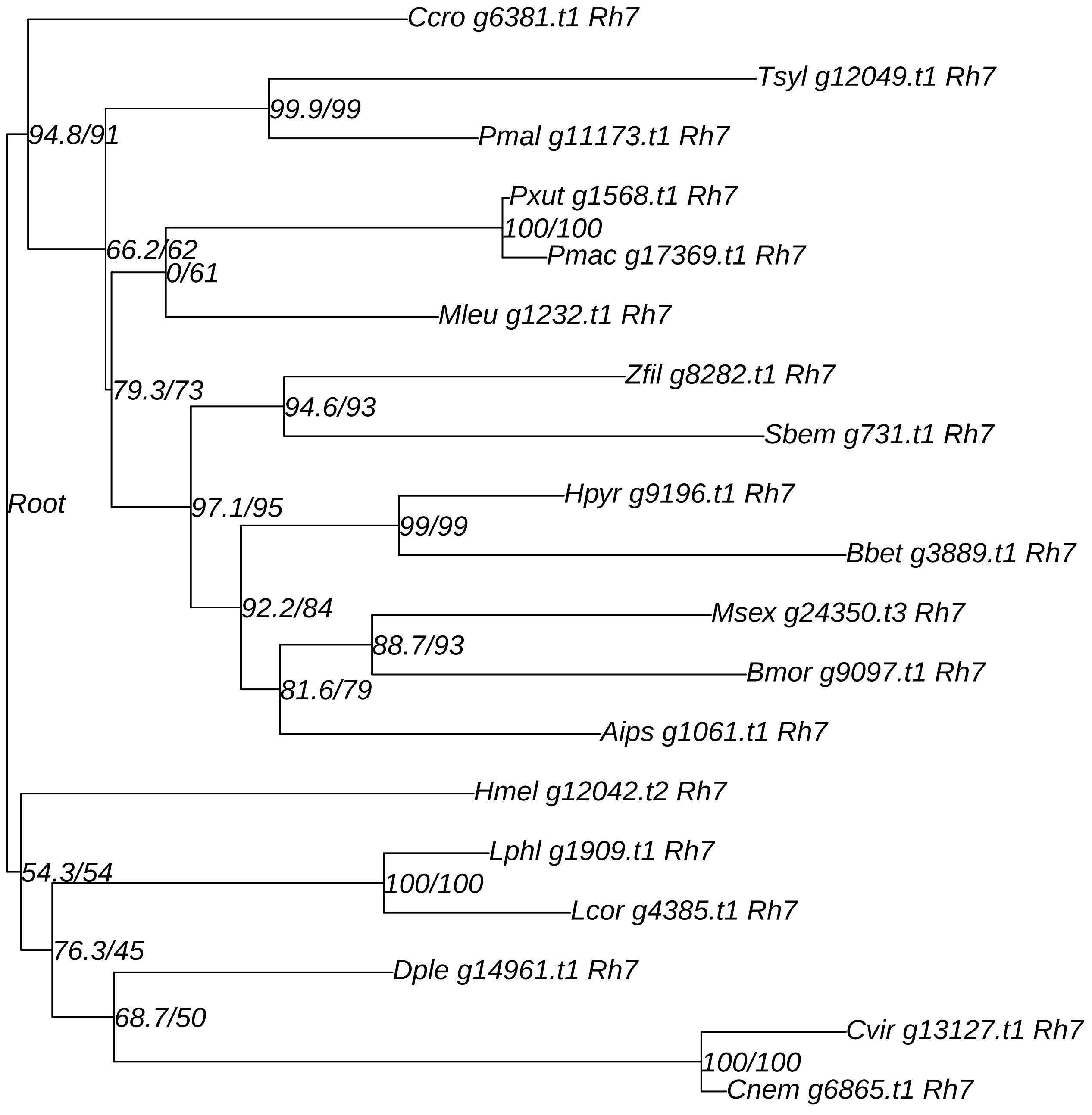

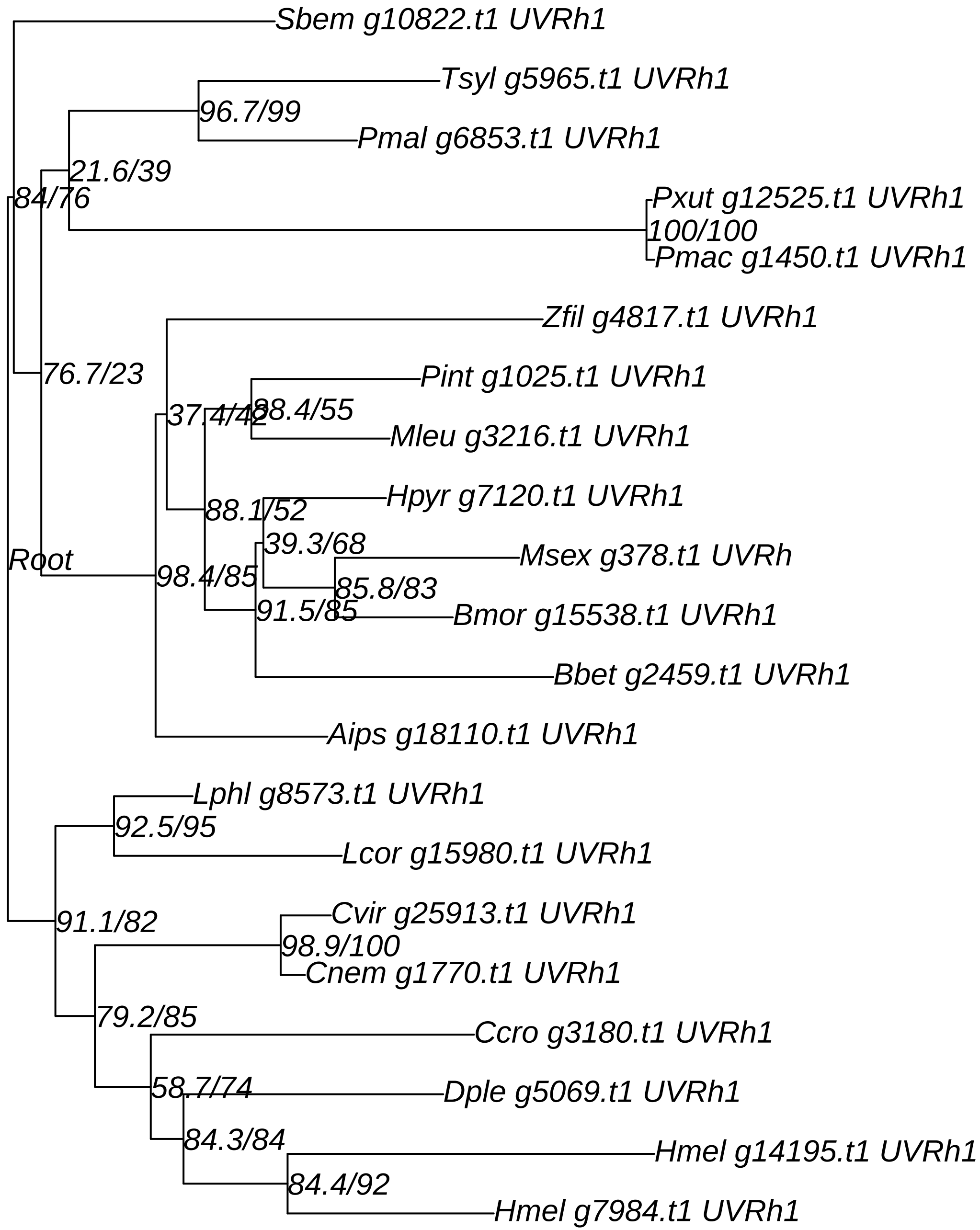

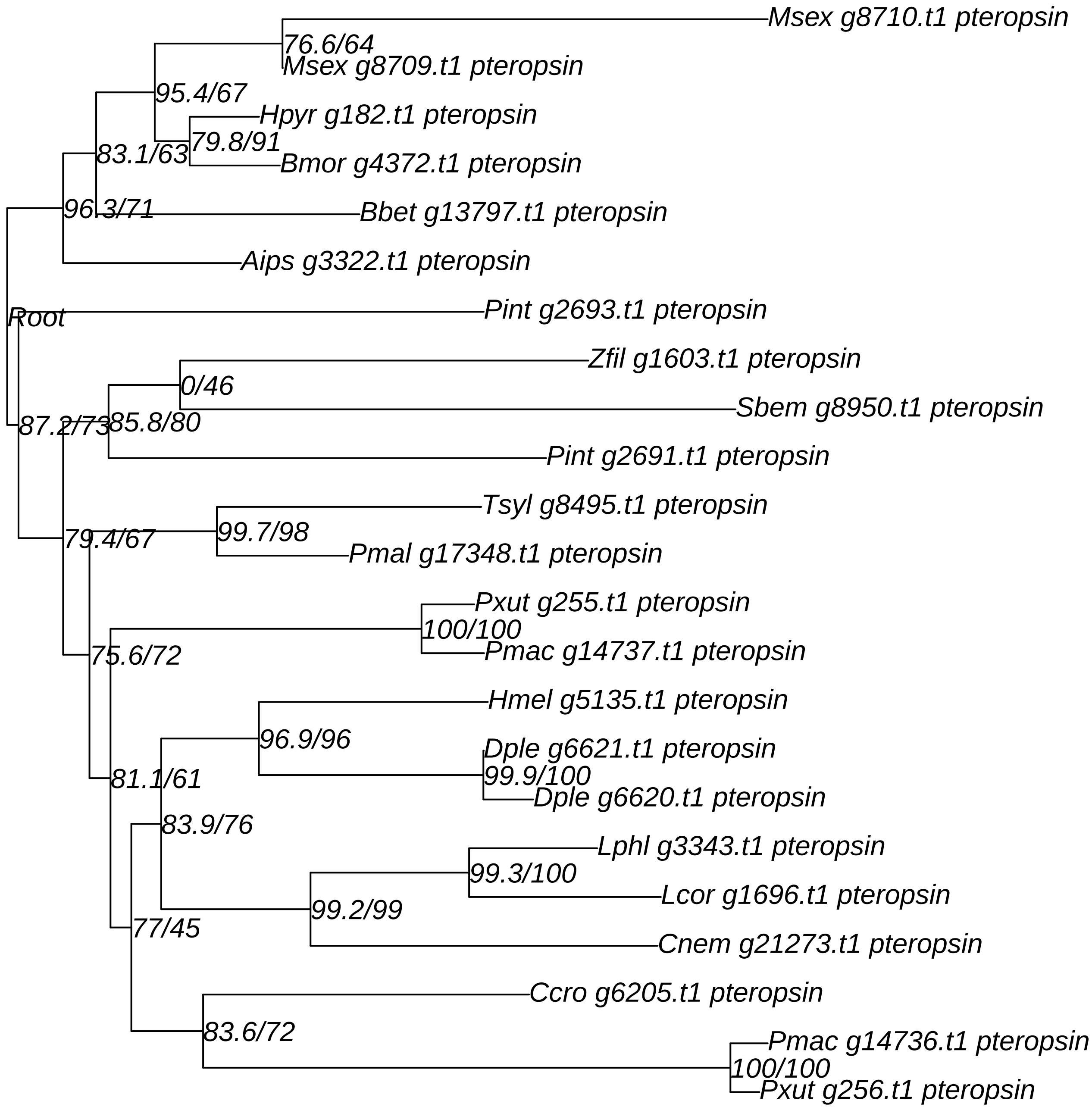

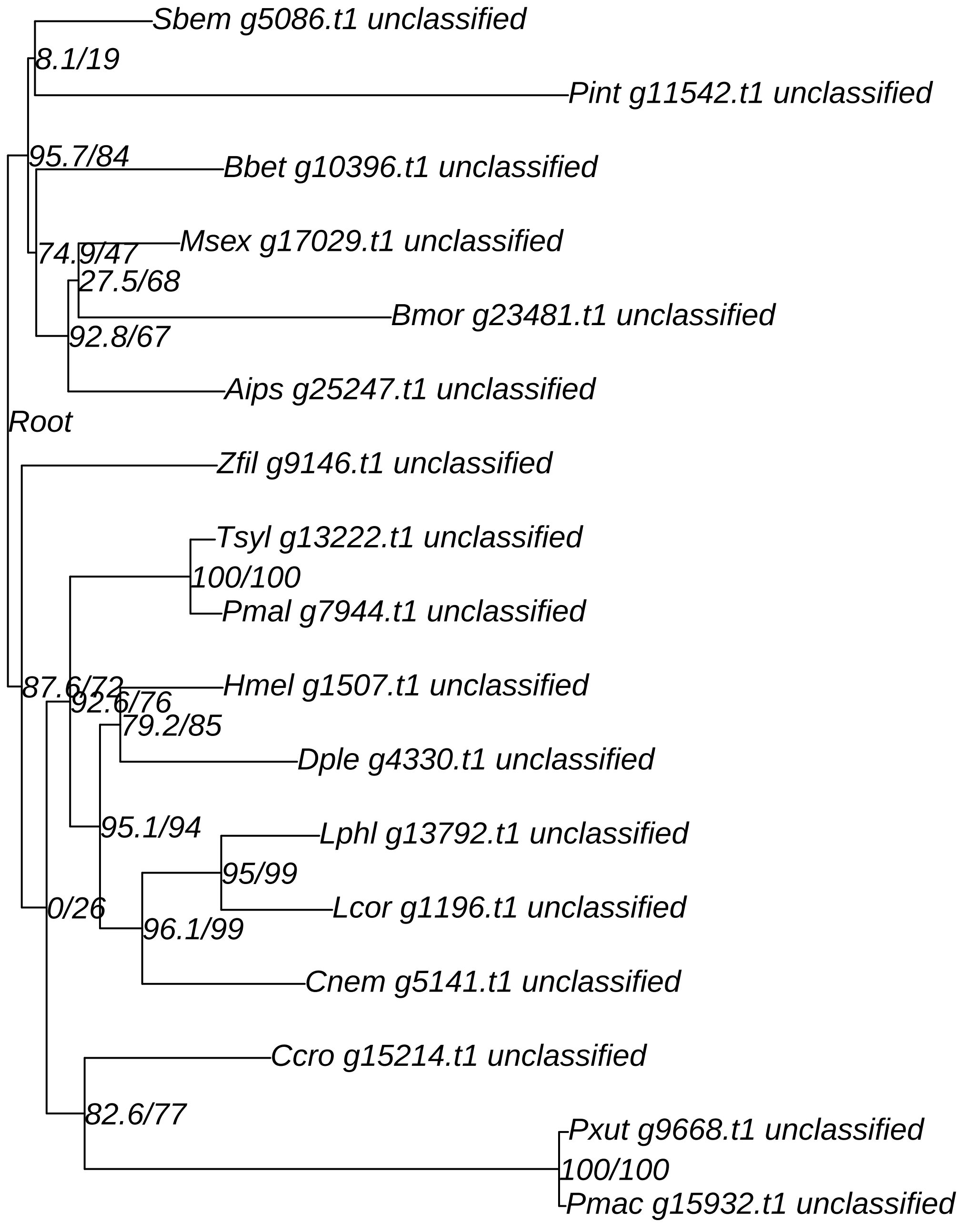

### Supplementary Data 13: Reconciled gene trees ThirdKind

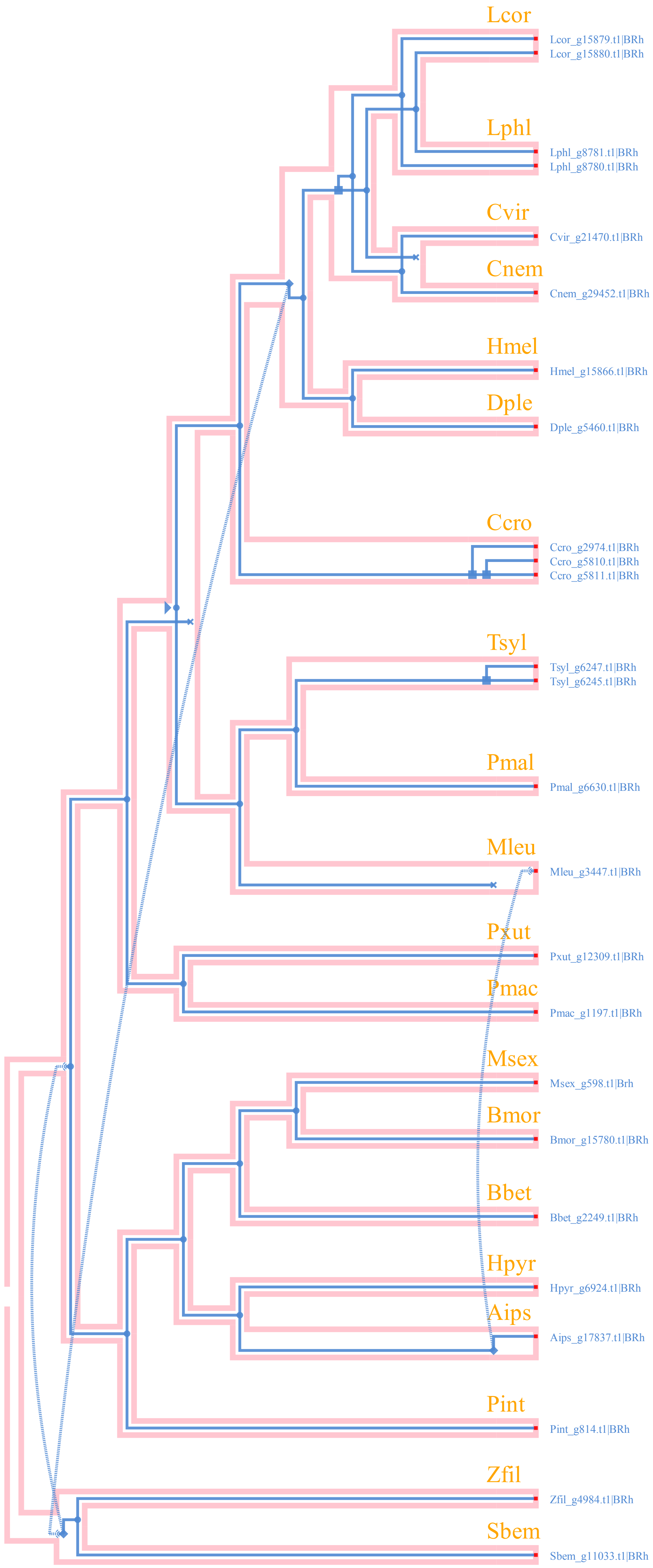

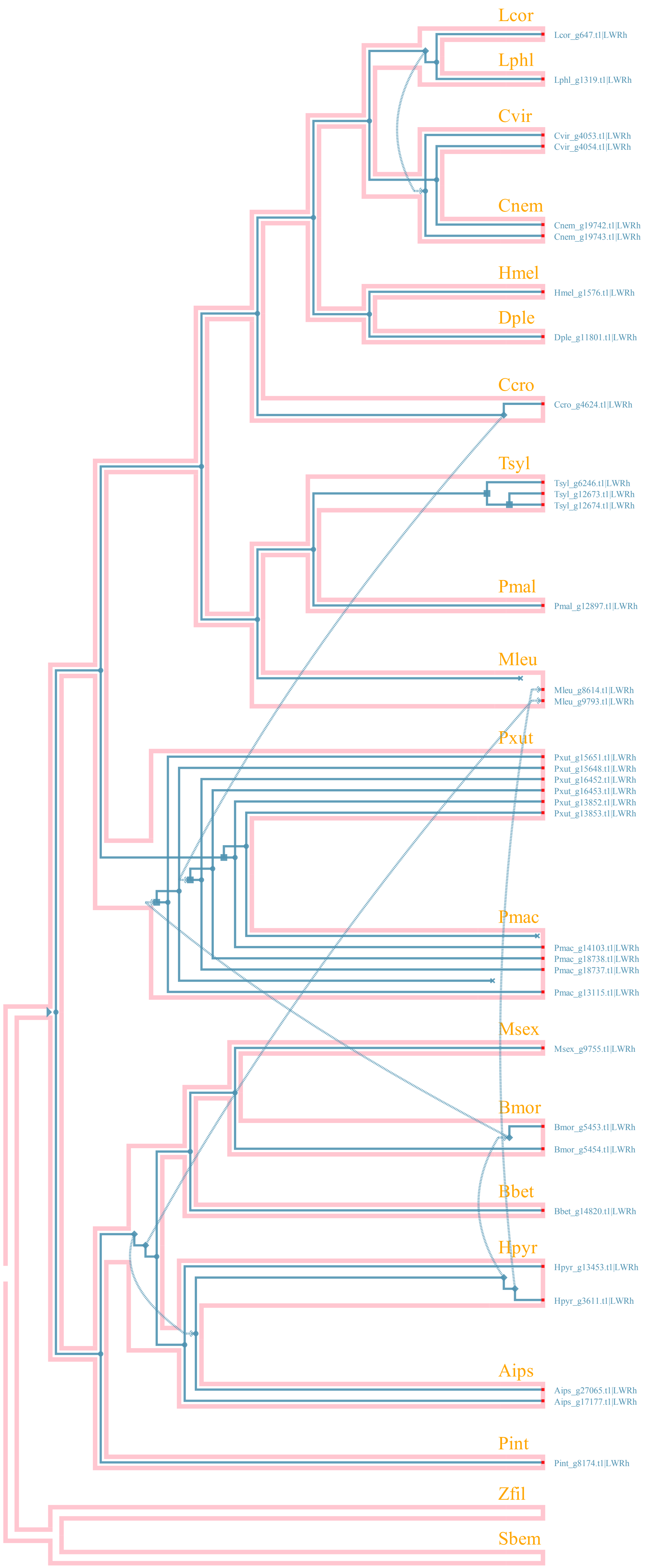

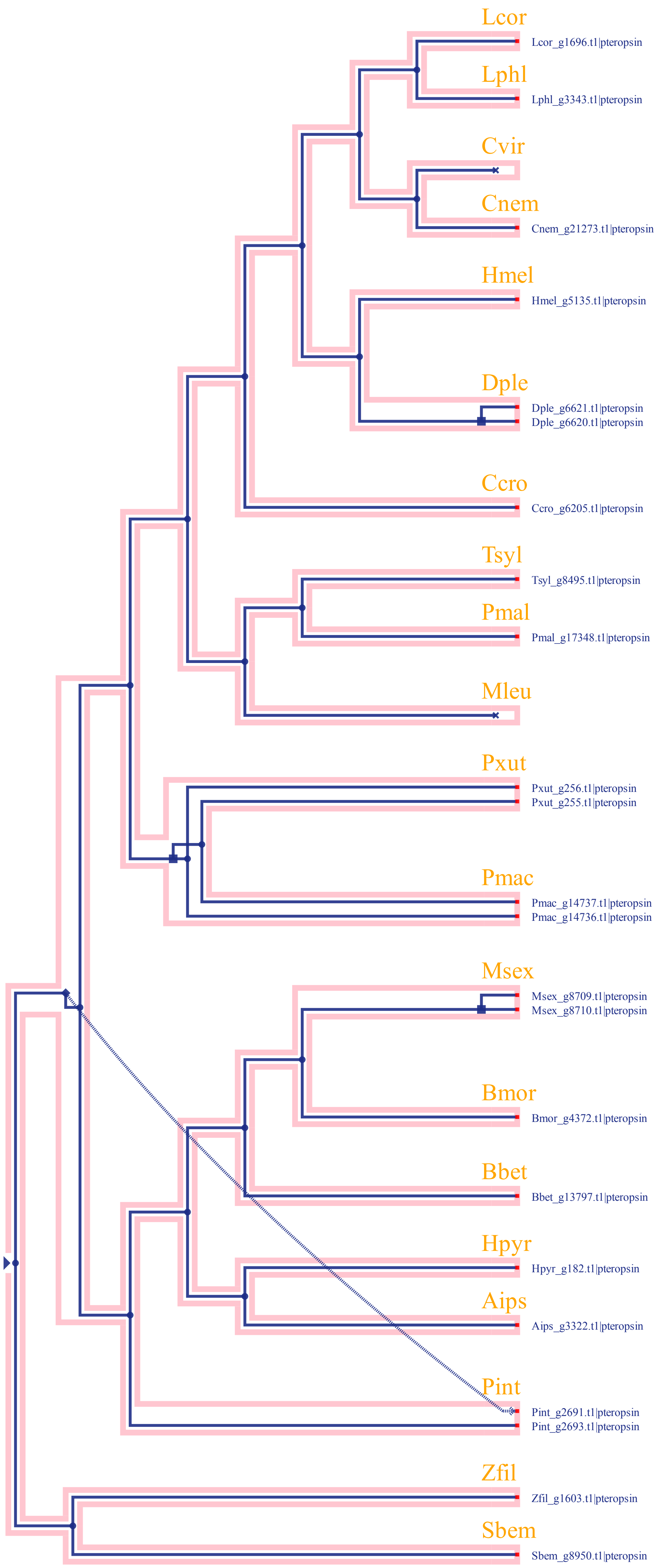

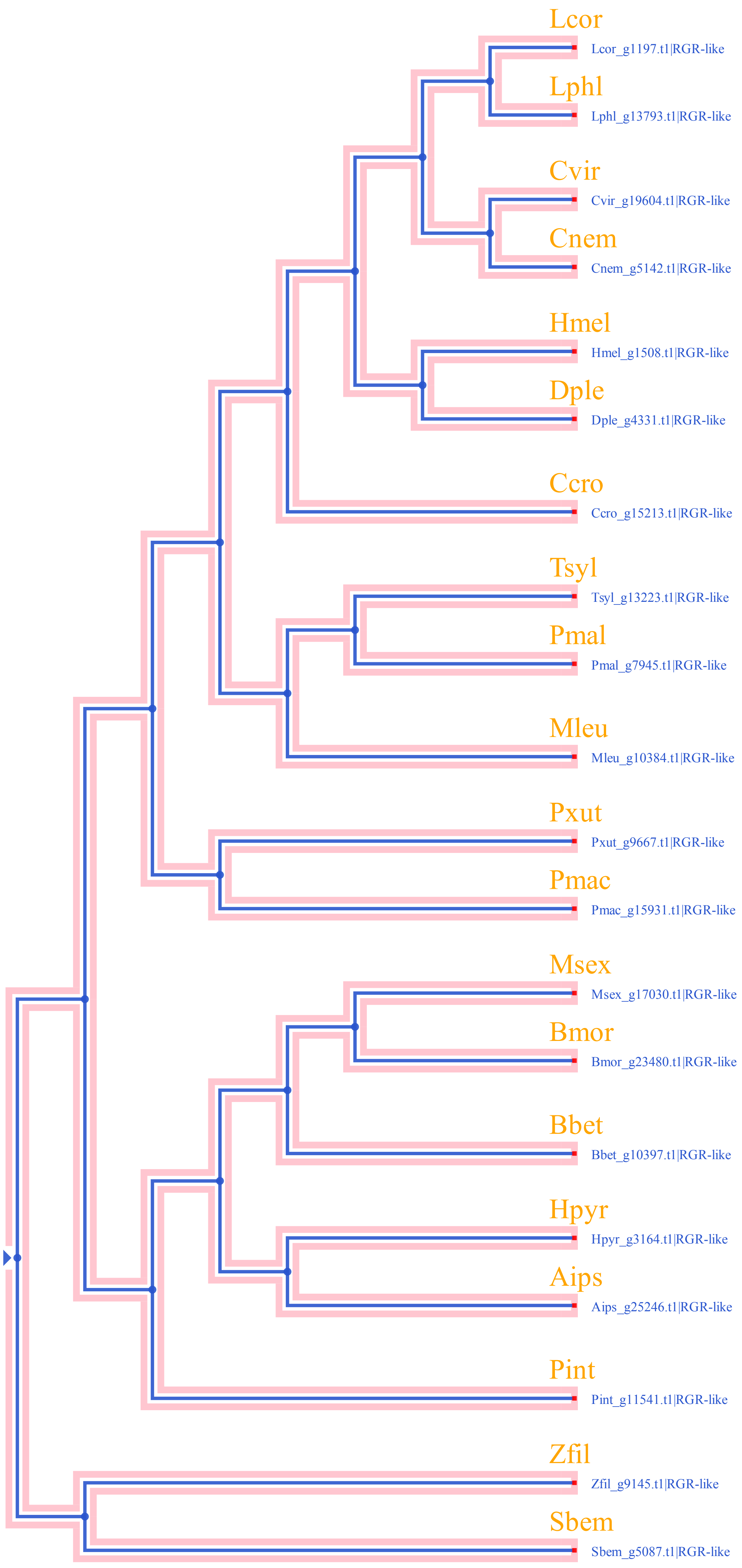

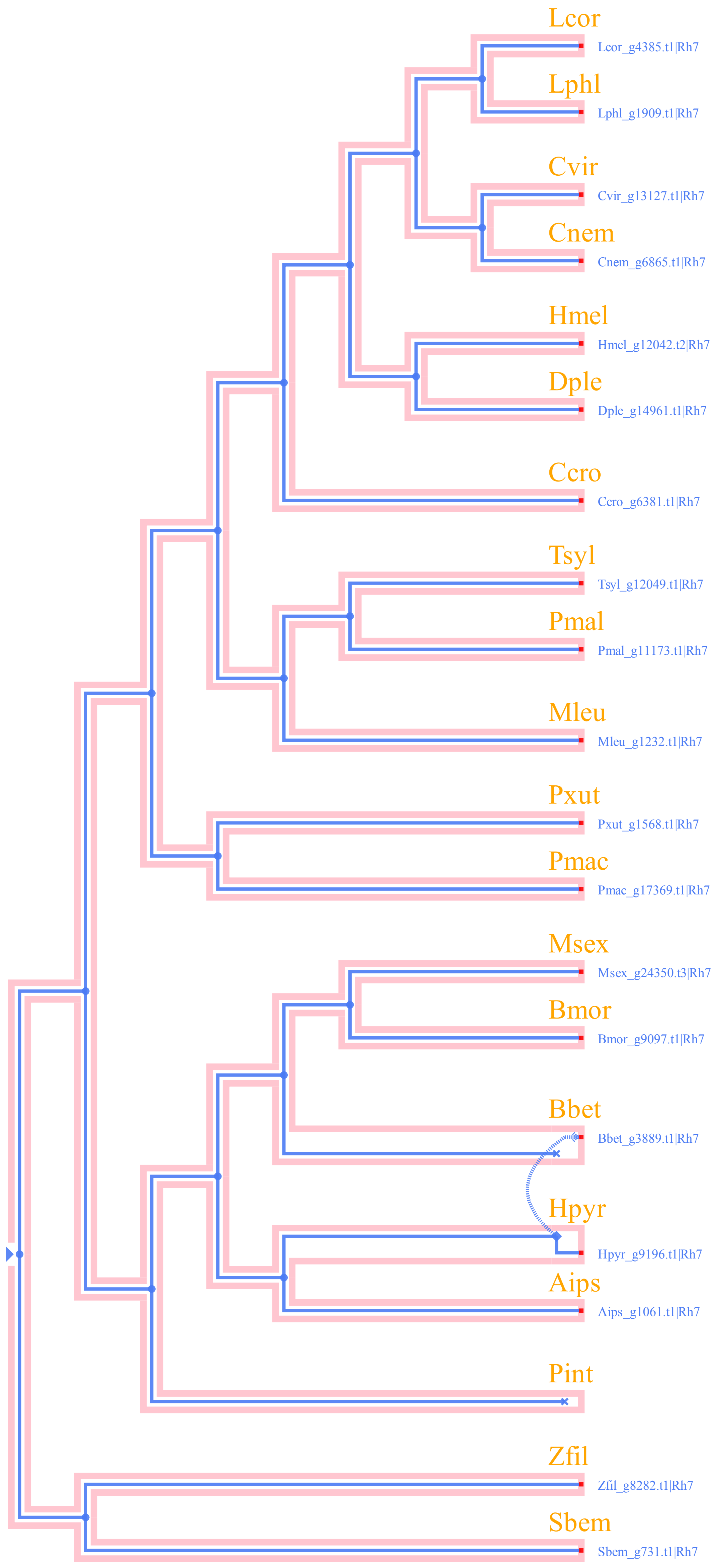

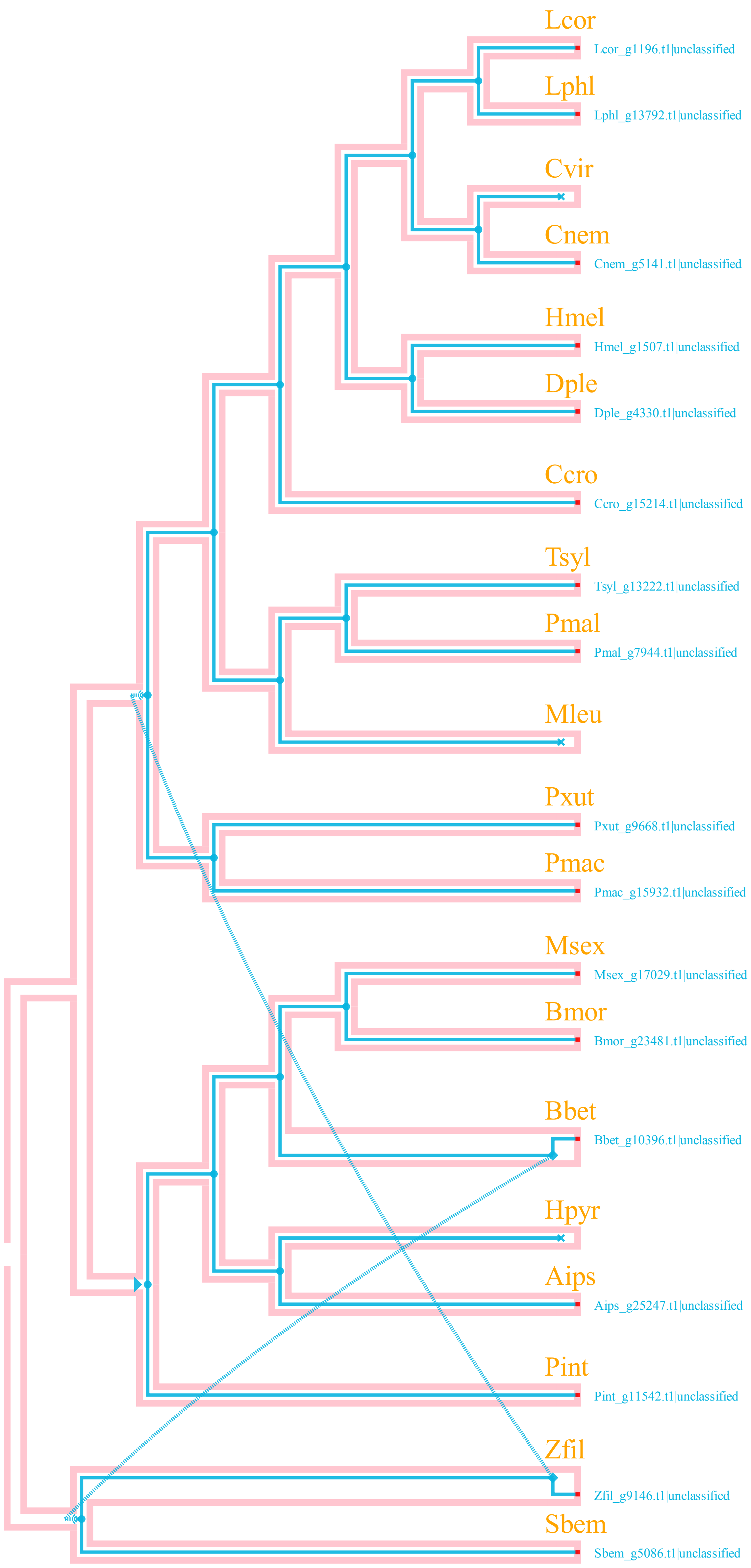

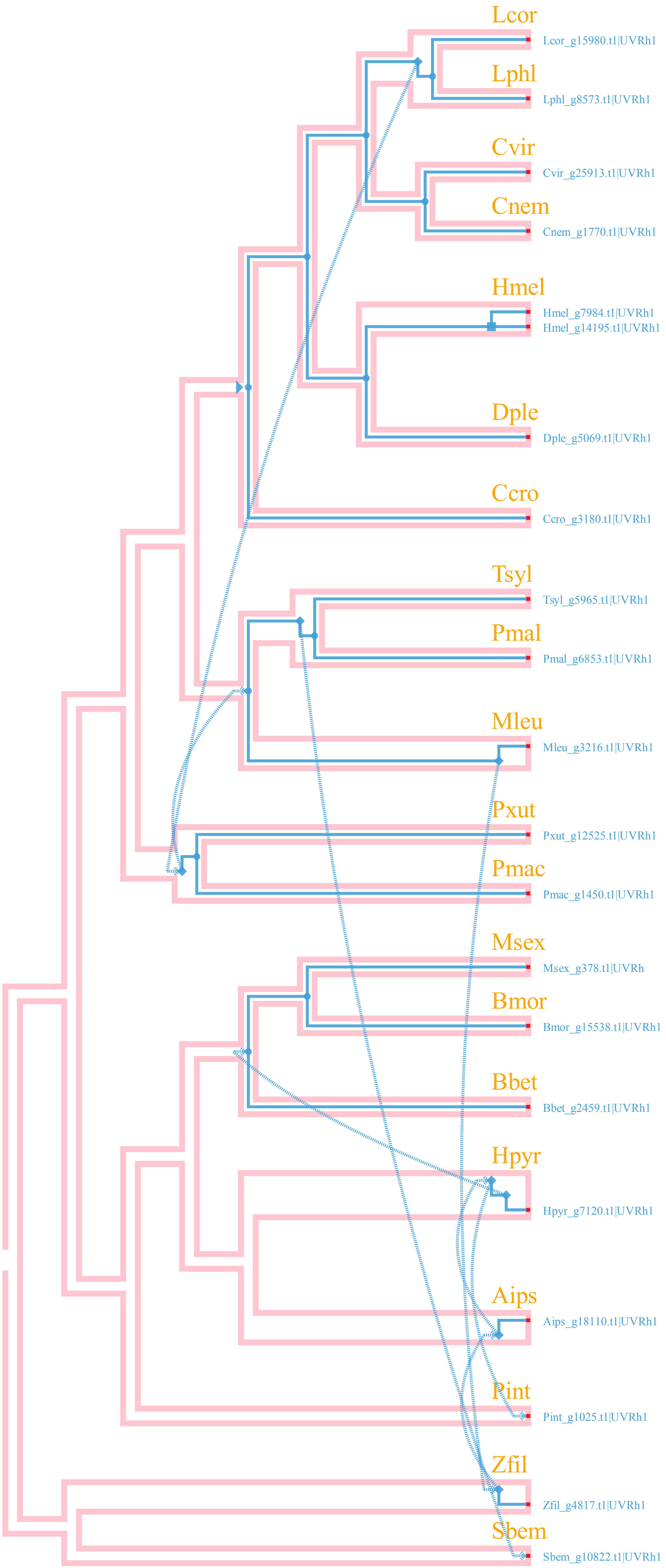
